# Footprints of human migration in the population structure of wild wine yeast

**DOI:** 10.1101/2024.08.08.607167

**Authors:** Jacqueline J. Peña, Eduardo FC Scopel, Audrey K. Ward, Douda Bensasson

## Abstract

Humans have a long history of fermenting food and beverages that led to domestication of the wine yeast, *Saccharomyces cerevisiae.* Despite their tight companionship with humans, yeast species that are domesticated or pathogenic can also live on trees. Here we used over 300 genomes of *S. cerevisiae* from oaks and other trees to determine whether tree-associated populations are genetically distinct from domesticated lineages and estimate the timing of forest lineage divergence. We found populations on trees are highly structured within Europe, Japan, and North America. Approximate estimates of when forest lineages diverged out of Asia and into North America and Europe coincide with the end of the last ice age, the spread of agriculture, and the onset of fermentation by humans. It appears that migration from human-associated environments to trees is ongoing. Indeed, patterns of ancestry in the genomes of three recent migrants from the trees of North America to Europe could be explained by the human response to the Great French Wine Blight. Our results suggest that human-assisted migration affects forest populations, albeit rarely. Such migration events may even have shaped the global distribution of *S. cerevisiae*. Given the potential for lasting impacts due to yeast migration between human and natural environments, it seems important to understand the evolution of human commensals and pathogens in wild niches.

## INTRODUCTION

Since the last ice age, humans have transitioned from a hunter-gatherer to a sedentary lifestyle and developed new technologies for preserving food including fermentation (McGovern, 2003). The wine yeast, *Saccharomyces cerevisiae,* is a driver of such fermentations and is used to produce beer, wine, sake, cocoa, and coffee (Marsit *et al*. 2017). The earliest archaeological evidence of fermented rice, honey, and fruit in China dates to 7,000 BCE (McGovern *et al*. 2004). Fermented beer was first discovered in ancient Sumerian vessels from 6,000 BCE (Michel *et al*. 1992) and there is evidence for wine production between 6,000 and 4,000 BCE in Iran, the Caucasus, and Mesopotamia (Pretorius, 2000; McGovern, 2003). Wine production then spread throughout the Mediterranean and was prevalent across Europe and Northern Africa by 500 BCE (Pretorius, 2000).

Today, the population genetics of *S. cerevisiae* shows imprints of domestication with several genetic lineages associated with distinct domestication events (Almeida *et al*. 2015; Duan *et al*. 2018; Fay *et al*. 2019; Fay & Benavides, 2005; Gallone *et al*. 2016; Gayevskiy *et al*. 2016; Legras *et al*. 2007; Legras *et al*. 2018; Liti *et al*. 2009; Peter *et al*. 2018; Schacherer *et al*. 2009). Deep sampling for wild strains from Chinese and Taiwanese forests revealed high levels of lineage diversity compared to all other lineages, and the current consensus is that East Asian forests likely harbored the ancestral source populations that gave rise to all global *S. cerevisiae* lineages (Duan *et al*. 2018; Lee *et al*. 2022; Wang *et al*. 2012). Even outside Asia, sampling of *S. cerevisiae* from natural environments shows a wild side to this human-associated yeast species; there are wild genetic lineages that are distinct from known domesticated lineages (Almeida *et al*. 2015; Cromie *et al*. 2013; Fay & Benavides, 2005; Han *et al*. 2021; Liti *et al*. 2009; Peter *et al*. 2018; Tilakaratna & Bensasson, 2017).

How much has human activity affected the ecology and evolution of wild populations of human-associated yeast species? Human pathogenic yeast such as *Candida albicans*, *Candida glabrata*, *Candida parapsilosis*, and *Candida tropicalis* can be isolated from trees and other plant habitats (Bensasson *et al*. 2019; Opulente *et al*. 2019; Robinson *et al*. 2016) and other forest yeast species are associated with humans (Boynton & Greig, 2014; Mozzachiodi *et al*. 2022). Here, we make use of the extensive genome data available for *S. cerevisiae* and focus on the population structure and phylogenetic relationships of wild *S. cerevisiae* from a single ecological niche. By studying strains from oaks and other trees, we characterize populations in an ancestral niche while avoiding the complications of genetic admixture more commonly seen in *S. cerevisiae* from fruit, flowers, and insects (Hyma & Fay, 2013; Tilakaratna & Bensasson, 2017). Specifically, we identified tree-associated populations and estimated the timing of wild yeast migration events. We show that (i) tree-associated *S. cerevisiae* populations are highly structured, (ii) the worldwide spread of forest populations out of Asia probably occurred since the Last Ice Age, and lastly (iii) human-assisted migration is ongoing and may include migration from the USA to Europe since the Great French Wine Blight.

## MATERIALS AND METHODS

### Yeast strains and genome data

Whole-genome sequences for tree-associated strains were compiled from publicly available data (N = 295; Table S1) (Almeida *et al*. 2015; Barbosa *et al*. 2016; Bergström *et al*. 2014; Duan *et al*. 2018; Fay *et al*. 2019; Gayevskiy *et al*. 2016; Han *et al*. 2021; Pontes *et al*. 2020; Skelly *et al*. 2013; Song *et al*. 2015; Strope *et al*. 2015; Yue *et al*. 2017). We defined *S. cerevisiae* tree-associated strains as those isolated from tree bark, exudate and leaves from trees or litter, and we also included strains from any soil. Metadata was compiled for each genome sequence to include geographical origin, ecological substrate, and previously reported genetic clade associated with the strain (Table S1). New whole-genome sequence data was generated for tree-associated strains from Indiana and Kentucky (N = 7; Osburn *et al*. 2018), North Carolina (N = 9; Diezmann & Dietrich 2009), Europe (N = 4; Robinson *et al*. 2016) and for new *S. cerevisiae* strains from the bark of white oak (*Quercus alba)* and live oak (*Q. virginiana*) from Georgia, Florida, Pennsylvania, and North Carolina (N = 15; Bensasson lab). DNA was extracted from single yeast colonies using Promega Wizard® Genomic DNA purification kit following the manufacturer’s protocol for yeast except that only 75 units of lyticase (Sigma) were used in an overnight incubation at 37°C. For the generation of genome data from 22 strains from the Bensasson and Osburn labs, paired-end Illumina libraries were generated by the Georgia Genomics and Bioinformatics Core using the purePlex DNA Library Preparation Kit (GGBC Project #5256) or the Nextera DNA-Seq Library Protocol (GGBC Project #5881). Paired end sequencing was performed on the Illumina NextSeq2000 platform (2 × 150 bp). The remaining strains were sequenced at the University of Manchester as described in Almeida *et al*. (2015). Genome data is available on NCBI-SRA under project number PRJNA1090965 and includes a further 11 strains from Pennsylvania that are monosporic derivatives of previously studied strains (Table S1; Sniegowski *et al*. 2002).

To examine how tree-associated strains genetically cluster with non-tree strains, we constructed a reference panel of strains to represent published clades (1,034 strains from 42 clades; Duan *et al*. 2018; Peter *et al*. 2018). These reference-panel strains are different from tree-associated strains because they have been isolated from the human body (clinical), fermentation (e.g., wine and beer), baking, bioethanol, crops (e.g., sugar cane), decaying wood, fruit, flowers, insects, mushrooms, and water (e.g., sewers and oceans).

### Read mapping and base calling

Paired-end and single-end genomic Illumina reads were downloaded from the European Bioinformatics Institute (https://www.ebi.ac.uk/) or generated in this study. Reads were mapped to the *S. cerevisiae* reference genome, S288c (SacCer_Apr2011/sacCer3 from UCSC), using Burrows-Wheeler Aligner (bwa-mem, version 0.7.17; Li & Durbin, 2009). We used SAMtools to sort, index, and compress bam files and generated a consensus sequence using the mpileup function with the -I option to exclude indels (version 1.6; Li *et al*. 2009). Next, we used the BCFtools call function with the -c option to generate a consensus sequence (version 1.9) (Li *et al*. 2009) and converted from vcf format to fastq format in SAMtools using the vcfutils.pl vcf2fq command. Lastly, base calls with a phred-scaled quality score of less than 40 were treated as missing data (calls were converted to “N”) using seqtk seq -q 40 in SAMtools.

### Quality filtering steps

We removed tree-associated genome sequences from downstream population structure analysis by excluding duplicate genome sequences for strains already represented in the dataset (N = 32), strains with no geographical information (N = 1), and strains with average read depth below 30× (N = 18). Additionally, genome data were visualized to check for intraspecies cross-contaminated using vcf2allelePlot.pl (Bensasson *et al*. 2019; Scopel *et al*. 2021): a genome was removed (N = 19) if the frequency of unexpected SNP calls was over 5%. For the remaining tree-associated strains, we estimated genome-wide levels of heterozygosity using vcf2allelePlot.pl (Bensasson *et al*. 2019) which estimates the number of heterozygous point substitutions divided by the length of the high-quality genome sequence (phred-scaled quality score over 40). Most tree-associated strains are homozygous (Figure S1). We removed 35 tree-associated strains that are heterozygous (heterozygosity over 0.001) because they are difficult to represent in downstream phylogenetic analyses and may be interclade hybrids. After filtering, 236 tree-associated strains remained for population genetic analyses (Table S1 and Figure S2).

For the reference panel strains, we applied the same quality filtering steps and examined levels of genome-wide heterozygosity, removing 16 out of 42 published clades because all individuals were heterozygous (Figure S1 and S2).

### Population structure and genetic admixture

Whole-genome alignments were generated by concatenating the alignments for all 16 chromosomes into a single multiple-alignment file. Tree-associated strains were compared to reference strains after random selection of three strains per clade from the 26 published clades that remained after quality filtering (Table S2). One strain (BJ6) was randomly assigned to both CHN-IV and Far East Asia clades (N = 77 strains). Ambiguity codes or lowercase base calls were converted to N’s, and ends were filled to align to the same length. A neighbor-joining tree from genetic distances estimated by pairwise comparison of all genome sequences was constructed using MEGA-CC (version 10.0.5; Kumar *et al*. 2012). We used a Tamura-Nei substitution model (Tamura & Nei, 1993) with a gamma distribution and 100 bootstrap replicates. Gaps or missing data were discarded from each pairwise sequence comparison. For visualization, the neighbor-joining tree was rotated using ape (version 5.6.2; Paradis & Schliep, 2018) and further visualized using ggtree (version 3.4.4; Xu *et al*. 2021; Yu *et al*. 2017, 2018).

Population structure and individual ancestry were estimated from SNP allele frequencies using ADMIXTURE (version 1.3.0; Alexander *et al*. 2009). Genome data for all strains was merged into a single alignment in variant call format (vcf) using BCFtools and mitochondrial DNA was removed. Non-variant sites were filtered out using the min-ac 1 function in BCFtools, which retains variants with at least one non-reference allele. Low-quality reads with a Phred-scaled quality score under 40 were removed using the minQ option in VCFtools (version 0.1.16; Danecek *et al*. 2011). Then we converted the single alignment vcf file to text formatted and binary files using PLINK (version 1.9; Purcell *et al*. 2007) for downstream admixture analyses. We assigned strains to distinct populations or genetic clusters (K) through repeated runs of ADMIXTURE. Runs assumed different numbers of genetic clusters from 4 to 40 with five replicates per K. We selected the run with the highest log likelihood value for each K and visualized population structure across different K’s (Figure S3). We used CLUMPAK, specifically ‘Distruct’, to align ancestry proportions (Q matrices) across different values of K (Kopelman *et al*. 2015). ADMIXTURE results were visualized as stacked bar plots using the pophelper R package (version 3.2.1; Francis, 2017; Figure S4). Distinct genetic clusters were verified if they showed monophyletic clades with at least 95% bootstrap support in a neighbor-joining tree (Figure 1A and Table S2). We selected the run with the highest number of verified clusters or clades based on monophyletic groups in the neighbor-joining phylogeny and whether strains grouped by geography.

**Figure 1.**
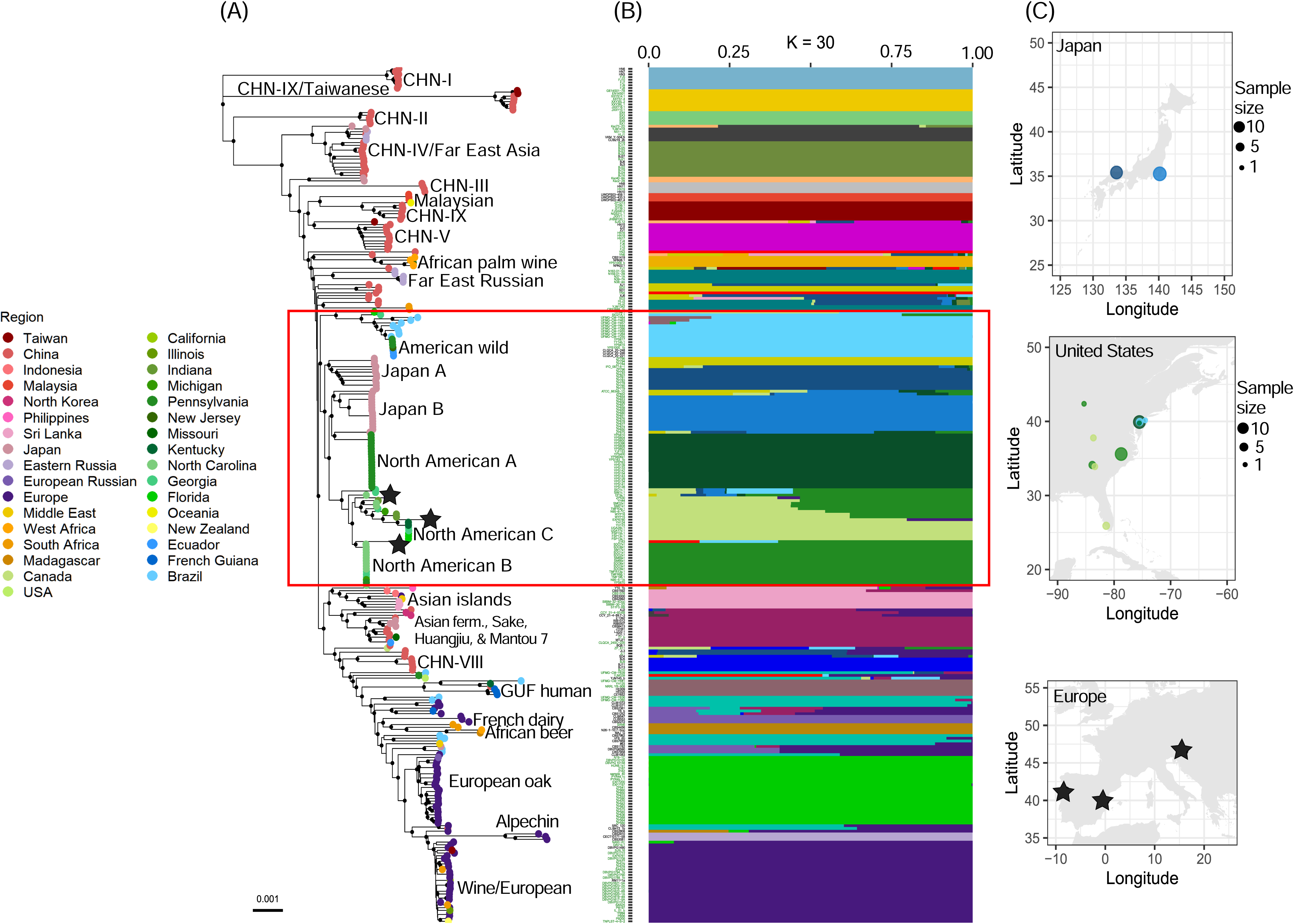
Trees harbor numerous genetically distinct *S. cerevisiae* lineages with population substructure in North America and Japan. **(A)** Whole-genome neighbor-joining tree of 313 strains after excluding heterozygous strains (Table S1). Tree-associated strains are in green and reference panel strains in black. Black circles at nodes indicate bootstrap support > 95%. Colored circles at the tips of the tree show geographical origin. **(B)** ADMIXTURE plot with K = 30 showing the cluster ancestry proportion for each strain. **(C)** Maps showing the geographic source of Japanese, North American, and European tree-associated strains highlighted in the red box. Circle sizes are based on square-root transformed sample sizes Japanese and North American strains and color coded by ancestry from ADMIXTURE plot. We removed North American and Japanese admixed strains (< 90% percent ancestry) from maps for simplicity.

Phylogenomic relationships among strains were further examined using a maximum likelihood tree after excluding admixed strains that showed recent genetic admixture when K = 30 (Figure S5 and Table S2). Admixed strains were defined as individuals whose percent ancestry from a single population is less than 90% in ADMIXTURE results for tree-associated and reference panel strains (Table S2). We included only three reference strains from CHN-IV and Far East Asia, which appear to be the same clade (Figure 1). We used a genome-wide alignment of SNPs mapped to the S228c reference genome to construct a phylogenomic tree with IQtree, ultrafast bootstrapping (version 1.6.12) (Minh *et al*. 2013; Nguyen *et al*. 2015), and a general time reversible model with a gamma distribution to estimate site heterogeneity. The maximum likelihood tree was visualized using ape and ggtree in R (version 4.2.2).

### Population substructure within Europe

Tree-associated strains are well sampled across Europe (N = 51 strains; Table S3) and have not been tested for population substructure. We used the same methods to analyze population substructure and genetic admixture among European tree-associated strains. ADMIXTURE was run by varying K from 2 to 8, and each K was repeated five times (Figure S6 and S7). Then, we constructed a phylogenomic tree using maximum likelihood estimation to infer phylogenetic relationships after removing one admixed strain when K = 4 (ZP541).

### *In silico* chromosome painting

To identify admixture positions within the genome, we used a chromosome painting approach with faChrompaint.pl and a 30 kb window size (Bensasson *et al*. 2019). The *in silico* chromosome painting identifies the most similar sequences in non-overlapping sliding windows to “paint” regions of the genome against a panel of predefined clades. We created a panel of genetically distinct clades based on population structure and phylogenomic analyses, which defined 25 monophyletic clades (Figure S4 and S5) and randomly selected three strains per clade to use as a “backbone” phylogeny (N = 75 backbone strains; Figure S8 and Table S4). For a strain of interest, each 30 kb window was compared to a multiple sequence alignment of backbone strains to assign or “paint” according to the most similar backbone sequence. Genomic regions that were diverged from all other backbone sequences (proportion of differing sites over 0.003) were painted white. This divergence threshold was determined based on within-clade pairwise comparisons for North American and European clades where the proportion of differing sites was below 0.003 for over 90% of 30 kb windows, and most between-clade pairwise comparisons showed divergence over 0.003 (Figure S9 and S10).

### Time divergence analysis

Time divergence analyses were performed on a single non-admixed locus (30-60 kb) from each backbone strain per chromosome (Figure S11). *In silico* chromosome painting analyses of backbone strains confirmed whether each backbone strain matched its primary clade assignment from allele frequency analyses using ADMIXTURE (Figure S8). For downstream time divergence analyses, we removed three backbone strains with less than 50% primary clade assignment using a chromosome painting approach, five backbone strains with over 10% secondary clade assignment and only included a single strain from the outgroup CHN-IX/Taiwanese clade (Table S5). A neighbor-joining tree was constructed for each locus per chromosome using the same methods previously mentioned and the phylogeny was rooted using the CHN-IX/Taiwanese strain (EN14S01) as the outgroup. Neighbor-joining trees and multiple sequence alignments were used to estimate time trees per chromosome using MEGA-CC with the RelTime-ML option (Tamura *et al*. 2018) using a Tamura-Nei substitution model (Tamura & Nei, 1993) with the default setting to consider all sites for branch length calculations (Figure S12).

Using neighbor-joining distance trees and estimates of the *S. cerevisiae* mutation rate, we estimated the approximate timing at which modern tree-associated lineages arose (i) outside Asia; (ii) into North America; and (iii) into Europe. We used genetic distance to estimate the time (T) to the most recent common ancestor (MRCA) in generations per year: T_MRCA_ = k / μ / generations per year; where k is the genetic distance to the MRCA of strains in a clade and μ is the point mutation rate per bp. We used the mutation rate of 1.67 × 10^-10^ point substitutions per site per generation, which was estimated from hundreds of point mutations after genome sequencing of mutation accumulation lines of diploid *S. cerevisiae* (Zhu *et al*. 2014). Under controlled laboratory settings at 30°C, wild diploid *S. cerevisiae* and its sister species *Saccharomyces paradoxus* have an average generation (doubling) time of 65 minutes in the presence of glucose and 125 minutes on nutrient-poor growth media (Kaya *et al*. 2021). Although glucose, fructose and sucrose are present in the bark of trees that harbor yeast (Sampaio & Gonçalves, 2008) it is likely less available than in the lab. In regions with tree-associated *S. cerevisiae*, historic temperatures were usually below 30°C (Table S8). With these considerations in mind, we made a rough estimate of the number of generations per year for yeast on trees assuming: (i) a 90 minute generation time because while the tree niche is likely less nutrient rich than laboratory growth media, some sugars are available; (ii) 12 hours growth per day to account for no growth at lower night time temperatures; (iii) and no growth for six months of the year to account for low temperatures in winter. The resulting estimate is an average of 4 generations per day or 1,460 generations per year. This is much lower than the number of generations possible at 30°C in nutrient rich media in laboratory conditions: 22 per day; 8,086 generations per year. It is also lower than the estimate of 2,920 generations per year used to estimate the age of the wine-associated lineage (Fay & Benavides, 2005), which may be less affected by cold winters and nutrient poor conditions.

## RESULTS

### Tree habitats harbor numerous genetically distinct lineages

Using genome data for 236 strains from oaks and other trees, we examined population structure among wild *S. cerevisiae* in this niche. Phylogenetic analyses of tree-associated strains and a reference panel of 77 strains from published clades (Duan *et al*. 2018; Peter *et al*. 2018) revealed several genetically distinct lineages that only occur on trees from China, Europe, Japan, North America, Russia, and Taiwan (Figure 1, Table S2 and Figure S5). These include previously studied wild lineages such as ‘CHN-IX’, ‘CHN-II,’ and ‘European oak’ (Duan *et al*. 2018; Peter *et al*. 2018), and more tree-associated lineages (see below).

In a larger unfiltered sample of 341 tree-associated strains, we also observed strains from clades that are usually associated with humans (Table S1). For example, there were 28 strains from the ‘Wine/European’ lineage from 21 field sites in 4 continents; 12 ‘South African Beer’ clade strains from 3 South African field sites (Han *et al*. 2021); 6 strains from the ‘Mixed Origin’ clade associated with baking and clinical strains (Peter *et al*. 2018); 3 strains from the ‘French Guiana Human’ clade from 3 continents; and occasional strains from ‘Asian fermentation’, ‘African honey wine’, ‘French dairy’ and ‘African palm wine’ (Table S1). Some of these were too heterozygous for further study (Table S1, Figure S2).

### Population substructure in the forest niches of North America and Japan

Phylogenetic analyses further revealed population substructure within North America and Japan, where each region has multiple genetically distinct populations (Figure 1A). Analysis of allele frequencies using ADMIXTURE confirmed that there are several genetically distinct populations in North America and Japan (Figure 1B). More specifically, there are at least four tree-associated (wild) American *S. cerevisiae* lineages in the eastern United States (Figure 1C) that are well-supported across phylogenetic and ADMIXTURE analyses (Figure 1, S4, and S5). Most wild strains from Pennsylvania are from a previously described lineage (Liti *et al*. 2009), and we refer to it here as ‘North American A’ (Table S1 and Table S2). There are two more North American wild lineages: ‘North American B’ and ‘North American C’, covering a broader geographical region than the North American A lineage (Figure 1C). North American B strains are from Georgia, North Carolina, a singleton strain from Michigan, and a singleton strain from Pennsylvania (Table S1 and Table S2). The North American C lineage occurs in strains from the southeastern United States (Kentucky, Georgia, and Florida). Lastly, wild strains from Ecuador and Brazil cluster with North American strains from Pennsylvania and New Jersey. We are coining this lineage as ‘American wild’ to reflect its Pan-American geography (Figure 1 and Figure S5) though some of these tree-associated strains were previously assigned to an ‘Ecuadorian’ clade (Peter *et al*. 2018) or a ‘Brazil 1’ clade (Barbosa *et al*. 2016).

In Japan, there are at least two wild *S. cerevisiae* populations: one from Hiruzen Highland, ‘Japan A’, and one from Chiba Prefecture, ‘Japan B’ (Figure 1 and Figure S5). Japan A is diverged from all North American lineages and Japan B (Figure 1 and S5). Japan B is most like the North American A lineage (Figure 1 and S5).

### Fine-scale population structure of wild *S. cerevisiae* from Europe

Tree-associated European yeast form two genetically distinct lineages: ‘Wine/European’ and ‘European oak’ (Figure 1 and Figure S5) that were previously known (Almeida *et al*. 2015; Tilakaratna & Bensasson 2017). Although the Wine/European lineage is usually recovered from wine fermentations, it sometimes occurs on trees, especially in vineyards (Gayevskiy *et al*. 2016; Hyma and Fay, 2013; Robinson *et al*. 2016). Initial analyses suggest population substructure within the European oak lineage (Figure 1, Figure S4). To better describe this substructure, we therefore ran separate ADMIXTURE and phylogenomic analyses for all tree-associated strains from Europe (N = 51; Table S3 and Figure 2). These analyses showed population substructure within the European oak lineage that correlates with geography (Figure 2, Figure S6 and S7). Specifically, there is evidence for five sub-lineages from: (i) Portugal and Spain, which we are coining ‘Iberian oak’ with support from both phylogenetic and ADMIXTURE analyses (Figure 2); and phylogenetic analyses suggest other distinct populations in (ii) Italy, (iii) Montenegro, (iv) Greece and Hungary, and (v) the North Caucasus (Figure 2). Additionally, several oak trees harbor strains from the ‘Wine/European’ winemaking lineage (Figure 2).

**Figure 2.**
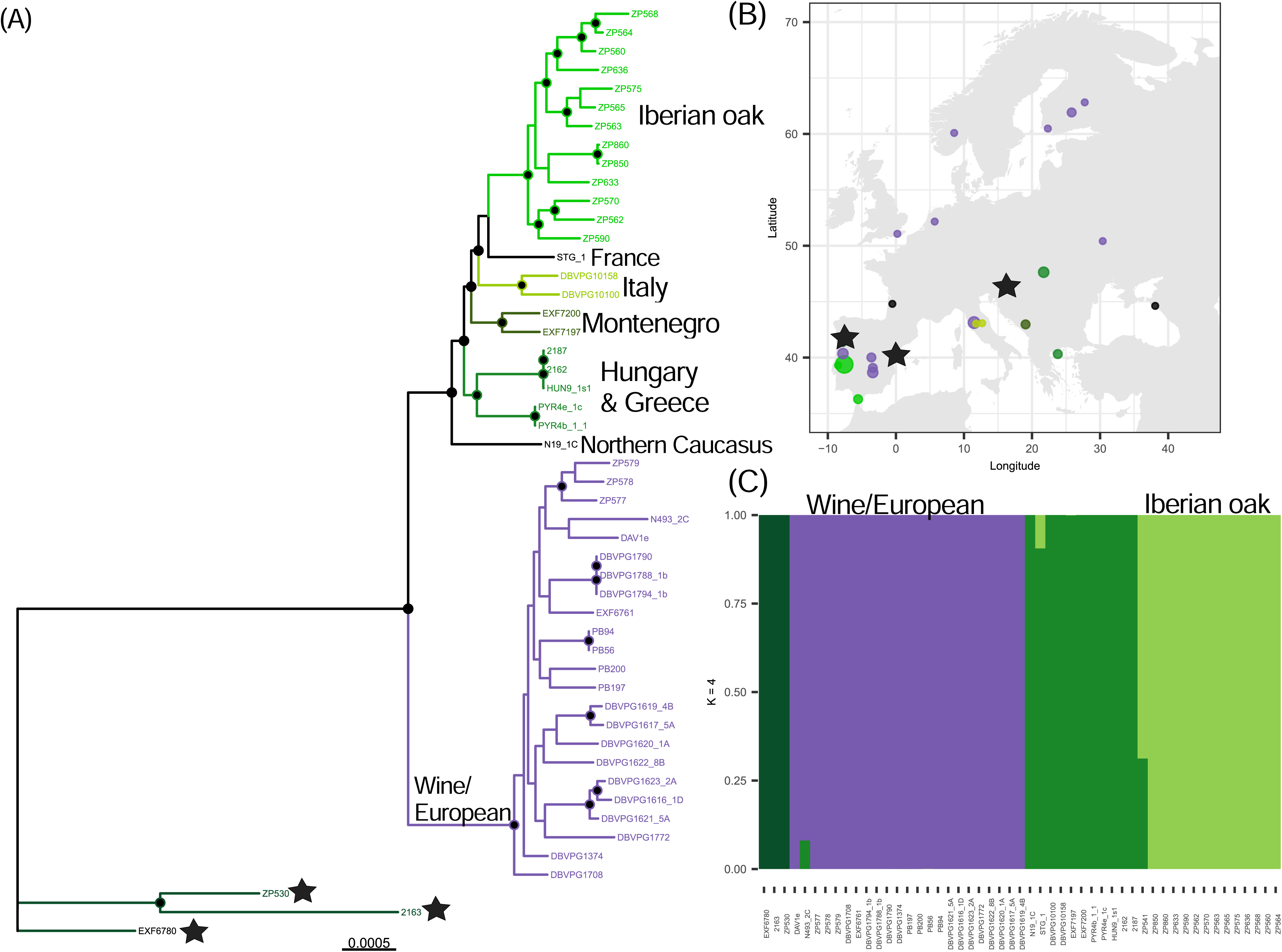
Fine-scale population structure of wild *S. cerevisiae* from Europe. **(A)** Whole-genome maximum likelihood phylogenetic tree of 50 strains after excluding one admixed strain, ZP541 from (C). Black circles at nodes indicate bootstrap support > 95%. Branches are color coded by geography or by ecology. Black stars at tree tips denote strains that genetically cluster with North American strains in Figure 1. **(B)** Map of Europe showing the geographic source of strains, circles are sized by the square-root transformed sample sizes and color coded by branch colors in the phylogenetic tree. Singleton strains from France and Northern Caucasus are colored in black. Black stars denote strains that genetically cluster with North American strains in Figure 1. **(C)** ADMIXTURE plot when K = 4 to examine percent ancestry per individual strain.

### Out-of-Asia migration of forest yeast since the last glacial maximum

We used a relative rate approach (Tamura *et al*. 2018) to estimate the timing of lineage divergences that likely correspond to (i) migration events out of Asia, (ii) into North America, and (iii) the origin of a Wine lineage distinct from European forest lineages. After excluding admixture, phylogenetic analysis of individual loci (30-60 kb) from each chromosome reproduced most genetically distinct clades defined in this study (Table S6 and Figure S12). Using these phylogenetic trees and assuming a mutation rate of 1.67 × 10^-10^ point substitutions per site per generation (Zhu *et al*. 2014) and 4 generations per day (see Methods), we generated rough estimates of divergence times. East Asia is the probable origin for *S. cerevisiae* (Duan *et al*. 2018; Han *et al*. 2021; Wang *et al*. 2012) and we estimate that non-Asian lineages diverged from those only found in Asia in approximately 10,000 BCE (mean = 10304, 95% CI 8754 - 11854 BCE; Figure 3). Out of 16 individual loci, 13 loci reproduced the European oak monophyletic clade and analyses were performed for the Wine-European oak split of these loci (Table S7 and Figure S12). European oak and the domesticated Wine/European lineage diverged around 3,000 BCE (mean = 3363, 95% CI 2248 - 4479 BCE; Figure 3). Out of 16 individual loci, 12 loci clustered the 3 North American clades (A-C), sometimes with Japan (A and B) lineages included. This mostly North American clade appears to diverge from others (Table S7 and Figure S12) around 5,000 BCE (mean = 5376, 95% CI 4550 - 6201 BCE; Figure 3). These estimates suggest global *S. cerevisiae* migrations occurred since the last glacial maximum (Figure 3).

**Figure 3.**
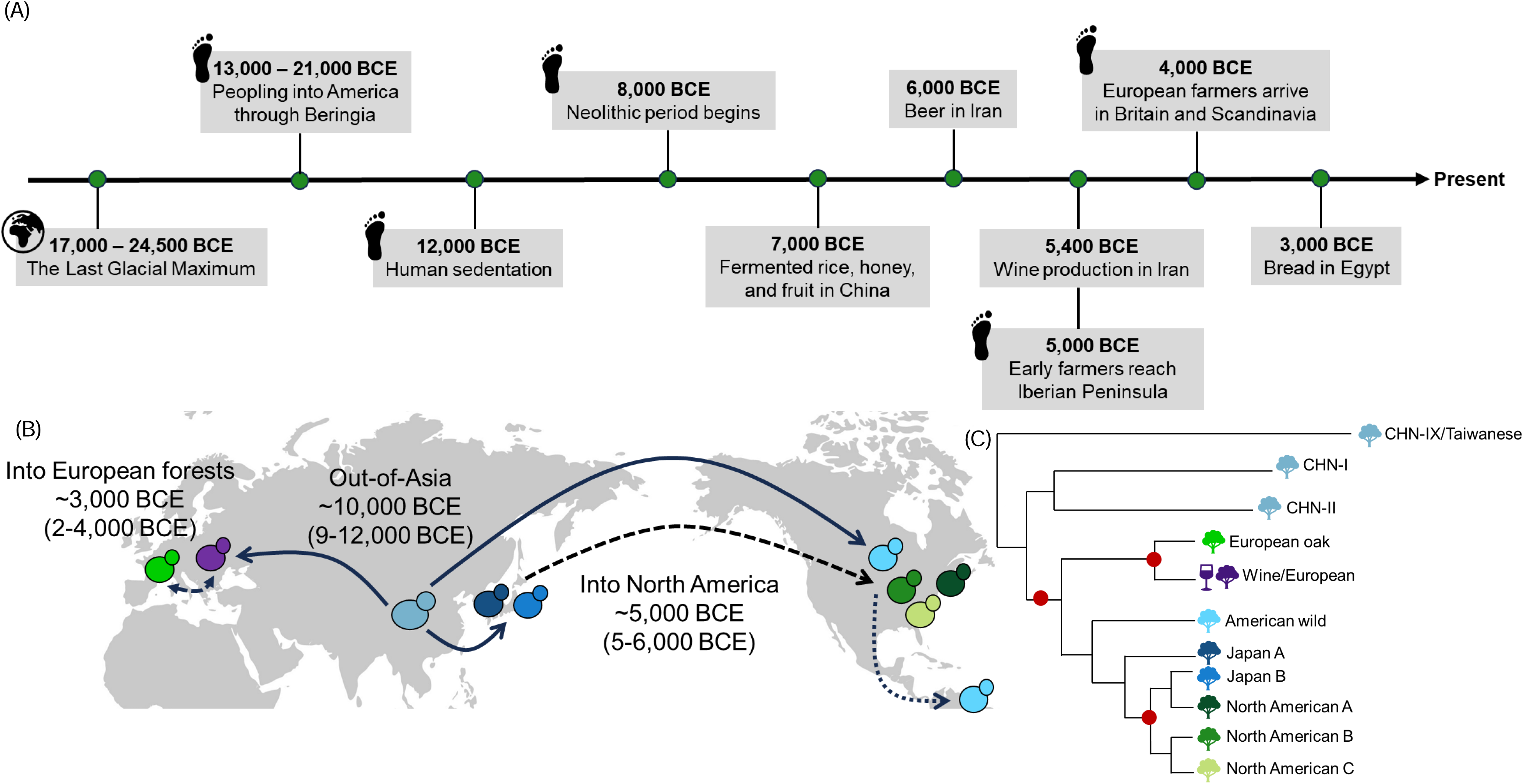
Out-of-Asia migration of forest yeast since the Last Ice Age. **(A)** Timeline showing early archaeological evidence of fermentation and human migration (dates from Clark et al., 2009; Marsit et al., 2017; Nielsen et al., 2017). **(B)** Map showing forest yeast migration events depicted with a red dot at nodes in (C) (i) out of Asia, (ii) into North America, and (iii) into European forests. Lineages are color coded by the lineages in the cladogram in (C). Dashed arrows indicate secondary migration events. **(C)** Cladogram showing phylogenetic relationships of lineages of interest for date estimation.

### Occasional strains in Europe resemble present-day North American lineages

There are three tree-associated strains from Europe (ZP530, 2163, and EXF6780) that resemble North American strains (black stars in Figure 1) and differ from all other European lineages (Figure 2). These genomes are from two different investigations where (i) ZP530 was isolated from chestnut (*Castanea sativa*) from Marão, Campeã, Portugal (Almeida *et al*. 2015), (ii) EXF6780 was isolated from sessile oak (*Quercus petraea*) from Velike Lašče, Kobila hill, Slovenia (Almeida *et al*. 2015) and (iii) 2163 was isolated from Portuguese oak (*Quercus faginea*) from Castellon, Spain (Peter *et al*. 2018). In the phylogenetic analysis, these strains differ from their most closely-related clades (North American B and C in Figure 1A) and appear to show some genetic admixture (Figure 1B). Did these strains arrive on European trees as a result of ancient migration or could their genetic distance from other North American strains be explained by recent admixture? To find out, we “painted” their chromosomes according to the clade of the most closely related strain (Figure 4A). EXF6780, ZP530, and 2163 were compared to our backbone phylogeny (Table S5), which revealed the strains are a mix of three lineages found in North America and that two strains have admixture from the lineage used to ferment grape wine (Figure 4).

**Figure 4.**
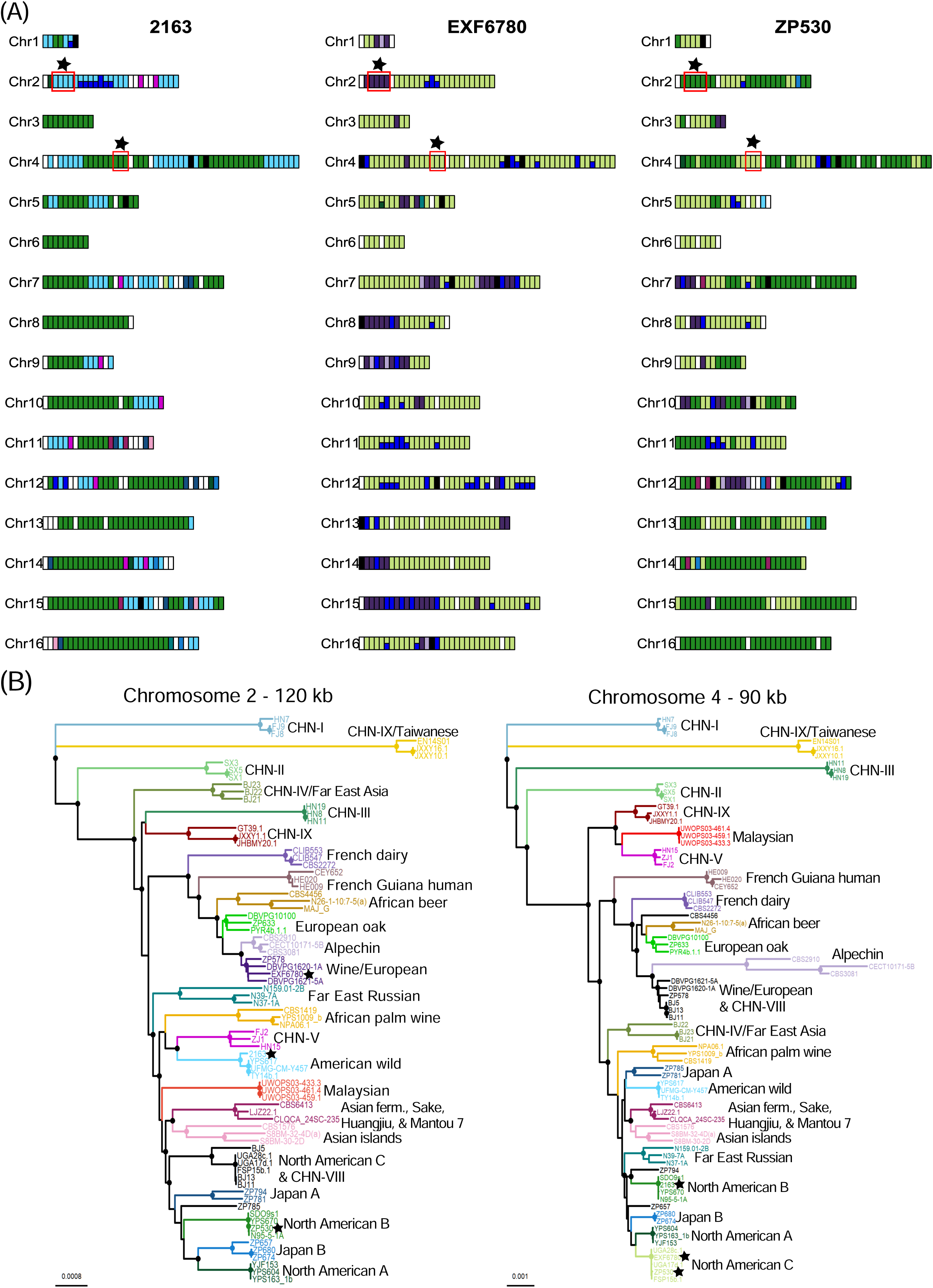
Occasional strains in Europe resemble present-day North American lineages. **(A)** Painted chromosomes of 2163, EXF6780, and ZP530 show admixture between multiple lineages. Genomic regions were “painted” based on the clade assignment of the most similar strain in 30 kb non-overlapping windows. Diverged regions were not colored (white) and were defined as regions that differed by 0.003 from all other strains in the backbone phylogeny. Black colored regions indicate low coverage. Colors are as in (B): North American B is forest green, American wild is light blue, North American C is light green, and Wine/European is dark purple. ZP530 is similar to North American B and North American C lineages. Genomic regions ranging from 90-120 kb were selected to confirm *in silico* chromosome painting results (red boxes with black stars). **(B)** Neighbor-joining phylogenetic trees for two loci. Solid circles at nodes indicate bootstrap support > 95%. Branches are color coded by clade. Phylogenetic analyses show that in the absence of admixture, 2163, EXF6780 and ZP530 (black stars) and very similar to strains from American wild, Wine/European, North American B, and C.

To assess genetic distances from modern North American lineages while accounting for admixture, we selected genomic regions from chromosomes 2 and 4 for phylogenetic analysis that did not show admixture from multiple clades (Figure 4A). Phylogenies with the effects of admixture removed in this way showed that strains 2163, EXF6780, and ZP530 seem almost identical to modern North American lineages across (Figure 4B). It therefore seems most likely that these ‘American European’ strains arrived very recently in Europe, and that admixture among North American and European/Wine lineages (Figure 4) explains their genetic distance from modern strains in the whole-genome phylogeny (Figure 1A).

## DISCUSSION

### Genetic isolation in forest niches

Before molecular methods showed that *S. cerevisiae* lives on trees (Sniegowski *et al*. 2002), people thought it lived only with humans and not in natural environments (Vaughan-Martini & Martini 1995). This resembles thinking for *Candida* pathogenic species before their recent discovery on trees and other plants (Bensasson *et al*. 2019; Opulente *et al*. 2019). Yet our results suggest that *S. cerevisiae* recently colonized woodlands many times and live there in relative isolation from humans. Phylogeographic analysis of forest *S. cerevisiae* populations shows that strains are recognizably from Iberian, French or Italian trees with further lineages occurring in Eastern Europe (Figures 1 and 2). There are at least four forest lineages in North America with regional differences; for example, North American C occurs in the southern USA, and North American A in Pennsylvania. Past analyses also show high population structure in *S. cerevisiae* from the primeval forests of East Asia (Wang *et al*. 2012; Lee *et al*. 2022) and in its sister species, *S. paradoxus* (Hénault *et al*. 2017; Leducq *et al*. 2014). *Saccharomyces* yeast are not air dispersed (Mortimer, 2000), therefore, it is not surprising that they show more population structure than other fungal microbes. Our observations for the tree niche contrast with the broader distribution of domesticated and fruit-associated lineages (Almeida *et al*. 2015; Duan *et al*. 2018; Gallone *et al*. 2016; Gonçalves *et al*. 2016; Lee *et al*. 2022; Legras *et al*. 2007; Legras *et al*. 2018; Peter *et al*. 2018) and are consistent with the proposal that animal-assisted long distance migration is relatively rare in forests (Magwene *et al*. 2011; Tilakaratna & Bensasson 2017).

The evidence for isolated forest populations includes phylogenetic analyses using whole genome, single chromosome, and single locus data (Figures 1A, 2A, 4B, S5, and S12) in addition to analyses of allele frequencies (Figures 1B, 2C, S7). Monophyletic tree-associated clades were reproducible across most chromosomes at the tips of phylogenetic trees, suggesting the fixation of many alleles for each lineage (Figures 1A, 2A, 4B, S5, and S12). Why do tree-associated lineages diverge rapidly? According to past estimates, *S. cerevisiae* reproduces sexually only once in hundreds or thousands of generations (Magwene *et al*. 2011; Lee *et al*. 2022) and the same is true for *S. paradoxus* (Tsai *et al*. 2008). Even when meiosis does occur, tree-associated *S. paradoxus* are almost always selfing (99% of sexual cycles, Tsai *et al*. 2008). Asexual reproduction of *S. cerevisiae* in the forest niche might lead to population bottlenecks and the local fixation of alleles by genetic drift.

### Global spread of *S. cerevisiae* forest populations since the last glacial maximum

Mutation rate estimates for *S. cerevisiae* applied to phylogenetic analyses suggest that the expansion of forest *S. cerevisiae* out of Asia likely occurred in the last 14,000 years (Figure 3 and S12). Divergence among the forest lineages of Europe and America is less deep than among the lineages of Asia (Figure 1A), so our analyses support an East Asian species origin (Wang *et al*. 2012; Lee *et al*. 2022). Averages estimated from phylogenies of 15 loci suggest that *S. cerevisiae* lineages migrated outside Asia around 9,000 - 12,000 BCE (Figure 3 and S12), which was when climate was warming after the last glacial maximum (Clark *et al*. 2009). The forest lineages that first diverged from East Asian populations include those occurring in South America (American wild, French Guiana human; Figure S12). The fine scale population structure within North American forests arose more recently; since the divergence of North American lineages (A-C) from most Asian lineages around 5,000 - 6,000 BCE. The population structure occurring on European trees arose since these lineages diverged from the Wine lineage from approximately 2,000 - 4,000 BCE.

The timing of these yeast migrations seems to roughly coincide with human sedentarization (∼ 12,000 BCE), the origins of agriculture (∼ 8,000 BCE), fermentation practices in Asia (7,000 BCE) and Europe (2,000 - 4,000 BCE) (Figure 3A; Liu *et al*. 2021; Marsit *et al*. 2017; McGovern *et al*. 2004; Nielsen *et al*. 2017). It is therefore possible that humans or their commensals carried yeast with their food as they moved around the world. The alternative, that yeast migrated naturally across the globe as the climate warmed, seems less likely for multiple reasons: (i) tree-associated lineages of *S. cerevisiae* have not been isolated from cool temperate regions (e.g. Canada, northern Europe), despite extensive searches (Charron *et al*. 2014; Johnson *et al*. 2004; Kowallik *et al*. 2015; Robinson *et al*. 2016), so a natural expansion through the cold climate of the Bering land bridge seems unlikely; (ii) human-associated migration can explain the relatively recent origin of European lineages (2,000 - 4,000 BCE), the much later spread to New Zealand in the last 1,000 years and their concentration near New Zealand vineyards (Gayevskiy *et al*. 2016).

Our time estimates are based on the well-studied mutation rate of *S. cerevisiae*. The estimate we use (1.67 × 10^-10^ per base per generation) is the most accurate; from hundreds of point mutations (867) observed in 145 genome sequences from diploid mutation accumulation lines (Zhu *et al*. 2014). Earlier mutation rate estimates did not differ greatly: 2.9 × 10^-10^ from mutations accumulated in a different diploid background (Nishant *et al*. 2010), 3.3 × 10^-10^ in haploids (Lynch *et al*. 2008), or 1.84 × 10^-10^ from reporter assays in haploid strains at individual loci (Drake 1991; Fay & Benavides 2005). The number of generations occurring in natural forest environments is more difficult to measure (Mozzachiodi *et al*. 2022). As in past analysis of wine yeast by Fay and Benavides (ca. 2,920 generations per year; 2005), we assume a lower growth rate than in the laboratory and only 12 hours of growth per day. The rate we use for trees (ca. 1,460 generations per year) assumes slower growth because of fewer nutrients on trees and no growth for 6 months of the year because of low temperatures (see Methods). For some parts of the species range, such as Florida and Georgia, generation times could be underestimated because some of the maximum temperatures in the coldest 6 months (18 - 29°C) and minimum nighttime temperatures in the hottest months (16 - 23°C) are also high enough for yeast growth (Table S8; Sweeney *et al*. 2004). While the fossil record is excellent for estimating divergences among yeast families or genera (Douzery *et al*. 2004; Marcet-Houben & Gabaldon 2015; Shen *et al*. 2018), the most recent fossil for ascomycotans dates to 417 million years ago (Douzery *et al*. 2004), and therefore may not be accurate for intraspecies divergences. Our timings are consistent with past estimates of divergence between wine and East Asian sake strains (9,900 BCE), among wine strains (1,700 BCE; Fay & Benavides 2005), the split between European oak and wine 8,300 BCE - 700 AD (Almeida *et al*. 2015), and the arrival of *S. cerevisiae* in New Zealand (Gayevskiy *et al*. 2016).

### Ongoing migration between human and tree-associated environments

Not all *S. cerevisiae* strains on trees are from tree-associated lineages. Strains from the European grape wine lineage and other human-associated lineages also live on trees (Gayevskiy *et al*. 2016; Hyma & Fay 2013; Robinson *et al*. 2016). Indeed, there were at least 10 migration events from Europe to New Zealand trees that happened in the last 1,000 years; since humans arrived in New Zealand (Gayevskiy *et al*. 2016). Here we observe strains on trees from clades connected with grape and African wines, Asian fermentations, brewing, baking, and clinical strains (Table S1, Figure S2), which suggests transmission between humans and trees is ongoing.

### Footprints of human activity in the genomes of European tree strains

Three strains from trees in Portugal, Spain, and Slovenia had genome sequences that were predominantly from North American forest clades. Chromosome painting of their genomes shows these strains must be descended from three different transatlantic migrants (Figure 4A). These migrations from North America must have been recent because analysis of large (90 and 120 kb) loci shows genomic tracts that look typical of current North American B, C and American wild lineages (Figure 4B). None of these American European strains resemble strains from the North American A lineage, which only occurred in Pennsylvania (Table S1). Interestingly, strains from Portugal and Slovenia (ZP530 and EXF6780) resemble the North American C lineage (Figure 4), which we only observe in the southern USA (Figure 1C).

A potential explanation for the presence of North American tree-associated lineages in Europe is the human response to the Great French Wine Blight. In the 1850s, humans accidentally introduced an insect pest, *Phylloxera*, from North America to Europe that destroyed most European vineyards. Native American vines are naturally resistant to *Phylloxera*. The European wine industry was rescued by the mass import of vines from the southern USA to Europe, and from the late 1800s to the present day most European vines are grafted onto resistant North American grapevine rootstock (Campbell, 2004). In support of this explanation, Portuguese and Slovenian American strains show admixture from the Wine/European lineage, suggesting recent association with vineyards whereas European oak and North American forest lineages rarely show admixture from the wine lineage (Figure 4).

### Conclusion

In summary, our analyses show forests harbor many isolated *S. cerevisiae* populations that are distinct from human-associated lineages. The phylogeographic structure of tree-associated lineages implies that migrants from humans rarely establish in forest niches. Yet even rare events can shape the distribution of a species. The postglacial spread of forest *S. cerevisiae* out of Asia and into North America and Europe suggests that this substantial impact was driven by people. Consistent with this, we also observe footprints of ongoing human-assisted movement of forest yeast. Fungal microbes introduced into forests can transform landscapes when they are mutualists or parasites (Hoeksema *et al*. 2020; Averill *et al*. 2022). For intimate human commensals and occasional pathogens, such as *S. cerevisiae* and *Candida sp.*, it seems important to consider their evolution in non-human environments - especially since environmental fungal microbes may adapt to fungicide use or rising temperatures (Garcia-Solache & Casadevall 2010; Leducq *et al*. 2014, Kang *et al*. 2022, Lockhart *et al*. 2023).

## Acknowledgements

This work was funded by an NSF LSAMP Bridge to the Doctorate Fellowship Program award (grant number: 1702361) to JJP; by the Howard Hughes Medical Institute through the James H. Gilliam Fellowships for Advanced Study program (grant no. GT13544) awarded to JJP and DB; and by a National Science Foundation grant (IOS, no. 1946046) awarded to DB. For some of the new genomes used, we would like to thank the Florida State Parks Service, the Duke Forest Teaching and Research Laboratory, Tyler Arboretum, Thompson Mills Forest, Skidaway Institute of Oceanography, and the University of Georgia for their support of fieldwork, and Diana Ambrocio, Domenic Won and Linda Habersham for strain isolation. Past and current members of the Bensasson lab provided helpful feedback. We would like to acknowledge that most of the research performed for this study was on Cherokee, Yuchi, and Muscogee indigenous lands.

## Data Accessibility and Benefit-Sharing

Genome data is available on NCBI-SRA under project number PRJNA1090965.

## Author Contributions

The research was conceptualized and designed by JJP and DB. AKW, JJP and EFCS performed DNA extractions for genome sequencing. JJP and EFCS developed bioinformatic pipelines, obtained and curated the data. JJP performed analyses and data visualization. JJP and DB wrote the paper with input from EFCS and AKW.

