## Supplemental Figures for "Footprints of human migration in the population structure of wild wine yeast"

**Supplemental Figure 1** Genome-wide levels of heterozygosity of *Saccharomyces cerevisiae* partitioned by tree-associated (N = 172) (A) and reference or non-tree-associated strains (N = 881) (B) after excluding monosporic derivatives. For downstream population structure analyses, we applied a 0.001 heterozygosity cutoff (red vertical line) to exclude strains likely to be inter-clade hybrids, which would obscure phylogenetic relationships.

**Supplemental Figure 2** Genome-wide levels of heterozygosity for *S. cerevisiae* tree-associated (N = 172) (A) and non-tree-associated strains (N = 881) (B) partitioned by published clades after excluding monosporic derivatives. Heterozygous clades (highlighted in red) are where 90% or more of the strains within that clade have a genome-wide level of heterozygosity greater than 0.001 (horizontal red line). Numbers on the plots indicate the number of strains per clade.

**Supplemental Figure 3** Log likelihood values from ADMIXTURE analyses for each cluster (K) across five replicate runs among 313 tree-associated and reference panel (non-tree-associated) S. cerevisiae strains. (A) Boxplot summary of the log likelihood values with five replicate runs for each value of K. (B) The log likelihood values for each replicate run.

**Supplemental Figure 4** Population structure and admixture of *S. cerevisiae* tree-associated and reference panel strains after excluding heterozygous strains (> 0.001 heterozygosity). Lineages were estimated using ADMIXTURE from cluster (K) 4-40 with five replicate runs for each K. We selected the run with the highest loglikelihood value from each K; strains are ordered by their position in the neighbor-joining tree (Figure 1A). Tree-associated strains are highlighted in green text. The ADMIXTURE plot highlighted with a red asterisk (K = 30) is the model used in downstream analyses. It had distinct genetic clusters that matched monophyletic clades in the neighbor-joining tree (>95% bootstrap support).

**Supplemental Figure 5** Phylogenetic relationships of *S. cerevisiae*. Whole-genome phylogeny of tree-associated and reference panel strains after excluding admixed. The phylogeny was constructed using maximum likelihood estimation with IQ-TREE ultrafast bootstrapping (1,000 bootstraps) using a general-time reversible model with a gamma distribution. Filled circles at nodes show monophyletic clades with 100% bootstrap support.

**Supplemental Figure 6** Log likelihood values from ADMIXTURE analyses for each cluster (K) across five replicate runs among 51 wild *S. cerevisiae* from European woodlands. (A) Boxplot summary of the log likelihood values with five replicate runs for each value of K. (B) The log likelihood values for each replicate run.

**Supplemental Figure 7** The population structure and admixture of wild *S. cerevisiae* from Europe. Populations were estimated using ADMIXTURE and with varying cluster counts (K = 2-8) with five replicate runs per each K. Here we show the run that had the highest log likelihood value from each K and strains are ordered by their position in the neighbor-joining tree.

**Supplemental Figure 8** Chromosome painting for 77 strains used as a backbone phylogeny (Table S3). Genomic regions were colored based on the clade assignment of the most similar strain in 30 kb non-overlapping windows. Diverged regions were not colored (white) and were defined as regions with a maximum proportion of sites that differed by 0.003 from all other strains in the backbone phylogeny. Black colored regions indicate low coverage.

**Supplemental Figure 9** Histograms of within-clade and between-clade comparisons for North American A-C (A-C). Within-clade divergences are below the 95th quantile (vertical red line), while between-clade divergences are above 0.003 (vertical red line).

**Supplemental Figure 10** Histograms of within-clade and between-clade comparisons for Wine/European (A) and European oak (B). Within-clade divergences are below the 95th quantile (vertical red line), while between-clade divergences are above 0.003 (vertical red line).

**Supplemental Figure 11** *In silico* chromosome painting results of quality filtered backbone phylogeny strains grouped by chromosome of strains for time divergence analysis (Table S5). Red boxes indicate 60-90 kb genomic regions selected for time divergence analysis when strains are assigned to their primary clade (>50%). Diverged regions are white and were windows that differed by at least 0.003 from all other strains in the backbone phylogeny. Black colored regions indicate low coverage. Genomic regions that are equally similar to strains from multiple clades are colored yellow.

**Supplemental Figure 12** Time divergence results of selected 60-90 kb genomic regions of backbone phylogeny strains for each chromosome. Phylogenetic trees on the left are neighbor-joining trees of the selected genomic region. Black circles at nodes indicate bootstrap support >95%. Clades are labelled in blue text. Time calibrated trees are on the left and were estimated using the RelTime-ML option with default settings (Tamura et al., 2012). We used a CHN-IX/Taiwanese strain (EN14S01) to root the time tree. We calculated the time (T) since the most recent common ancestor (MRCA) in generations per year. Time divergence events are indicated with a red circle at a node for the following events: Out-of-Asia, Wine-European oak split, and North America/Japan split. Time estimates are shown as dates Before Common Era (BCE) (Table S7). Time estimates were not calculated for events when clades did not form monophyletic groups.

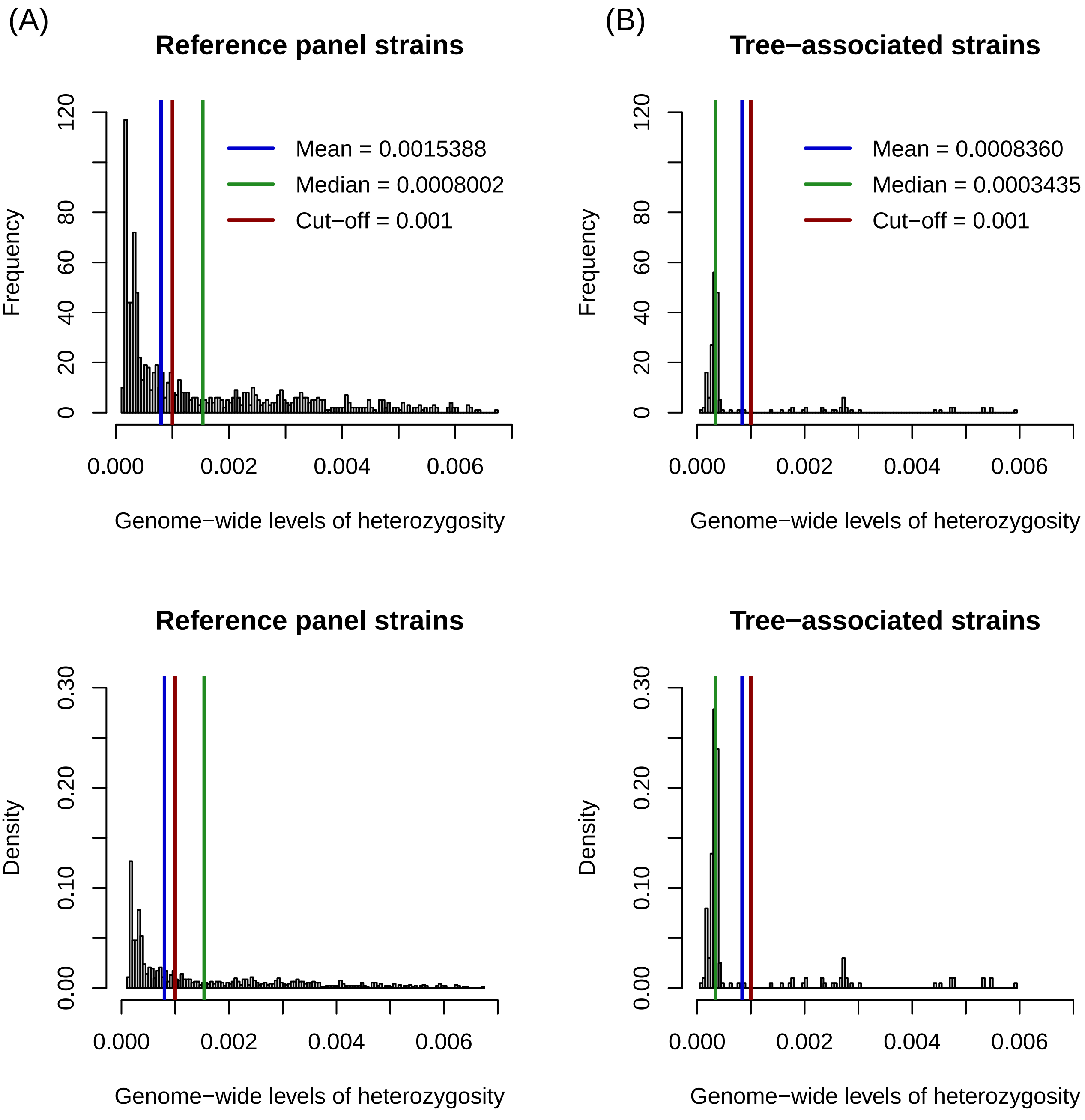
(A)
Reference panel strains
Mean = 0.0015388
Median = 0.0008002
Cut−off = 0.001
Frequency
Genome−wide levels of heterozygosity
(B)
Tree−associated strains
Mean = 0.0008360
Median = 0.0003435
Cut−off = 0.001
Frequency
Genome−wide levels of heterozygosity
Reference panel strains
Density
Genome−wide levels of heterozygosity
Tree−associated strains
Density
Genome−wide levels of heterozygosity

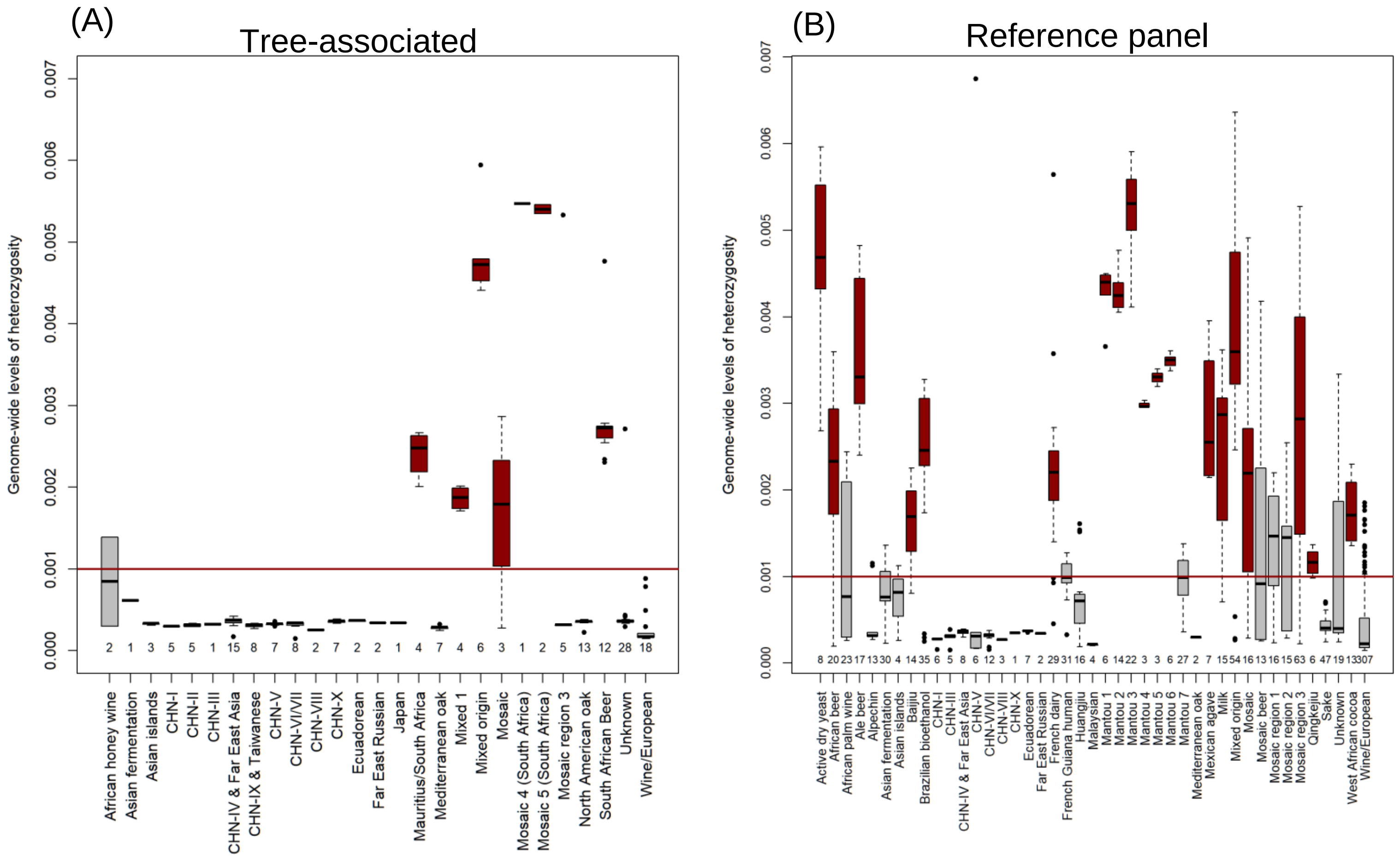
(A)
Tree-associated
Genome-wide levels of heterozygosity
0.000
0.001
0.002
0.003
0.004
0.005
0.006
0.007
2 1 3 5 5 1 15 8 7 8 2 7 2 2 1 4 7 4 6 3 1 2 5 13 12 28 18
African honey wine
Asian fermentation
Asian islands
CHN-I
CHN-II
CHN-III
CHN-IV & Far East Asia
CHN-IX & Taiwanese
CHN-V
CHN-VI/VII
CHN-VIII
CHN-X
Ecuadorean
Far East Russian
Japan
Mauritius/South Africa
Mediterranean oak
Mixed 1
Mixed origin
Mosaic
Mosaic 4 (South Africa)
Mosaic 5 (South Africa)
Mosaic region 3
North American oak
South African Beer
Unknown
Wine/European
(B)
Reference panel
Genome-wide levels of heterozygosity
0.000
0.001
0.002
0.003
0.004
0.005
0.006
0.007
8 20 23 17 13 30 4 14 35 6 5 8 6 12 3 1 7 2 29 31 16 4 6 14 22 3 3 6 27 2 7 15 54 16 13 16 15 63 6 47 19 13 307
Active dry yeast
African beer
African palm wine
Ale beer
Alpechin
Asian fermentation
Asian islands
Baijiu
Brazilian bioethanol
CHN-I
CHN-III
CHN-IV & Far East Asia
CHN-V
CHN-VI/VII
CHN-VIII
CHN-X
Ecuadorean
Far East Russian
French dairy
French Guiana human
Huangjiu
Malaysian
Mantou 1
Mantou 2
Mantou 3
Mantou 4
Mantou 5
Mantou 6
Mantou 7
Mediterranean oak
Mexican agave
Milk
Mixed origin
Mosaic
Mosaic beer
Mosaic region 1
Mosaic region 2
Mosaic region 3
Qingkeju
Sake
Unknown
West African cocoa
Wine/European

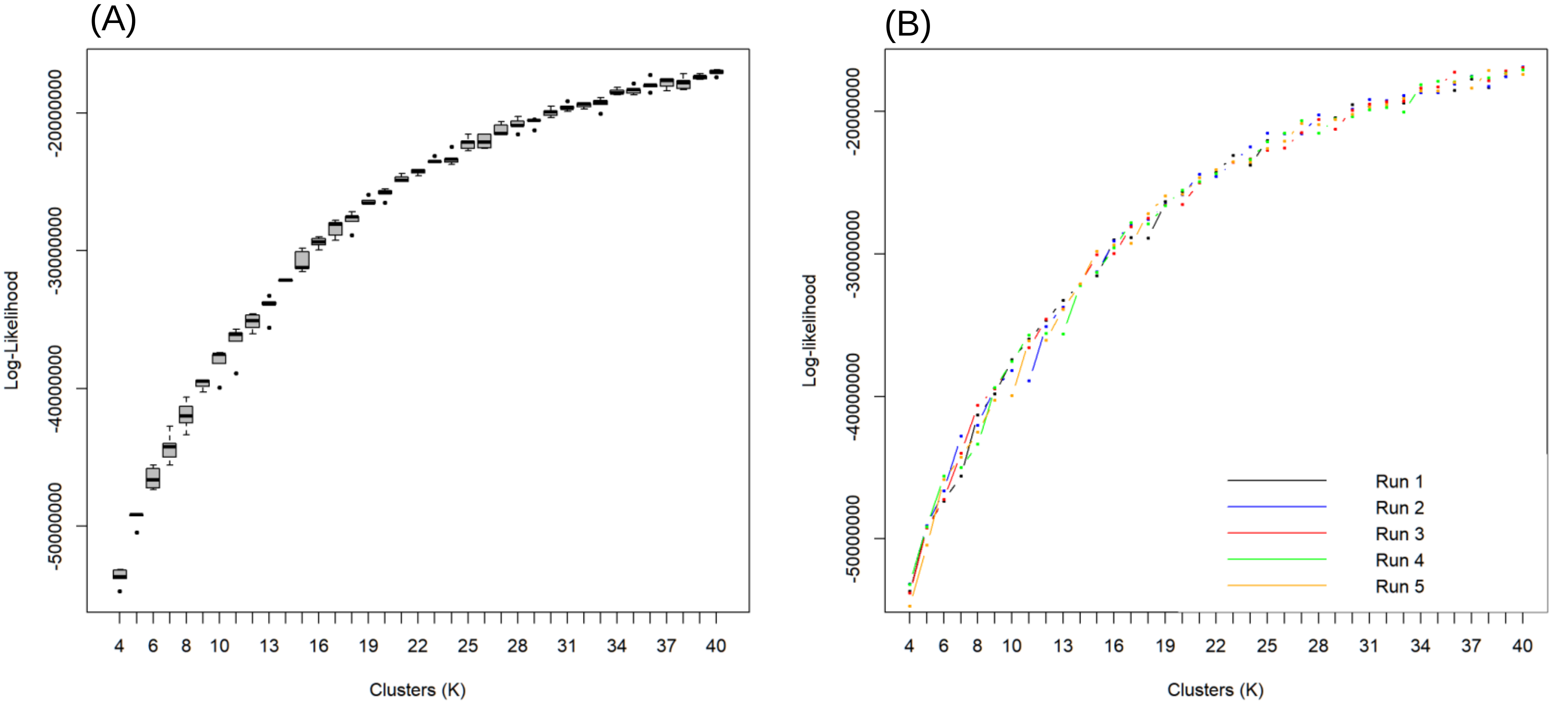
(A)
Log-Likelihood
-20000000
-30000000
-40000000
-50000000
4 6 8 10 13 16 19 22 25 28 31 34 37 40
Clusters (K)
(B)
Log-likelihood
Run 1
Run 2
Run 3
Run 4
Run 5

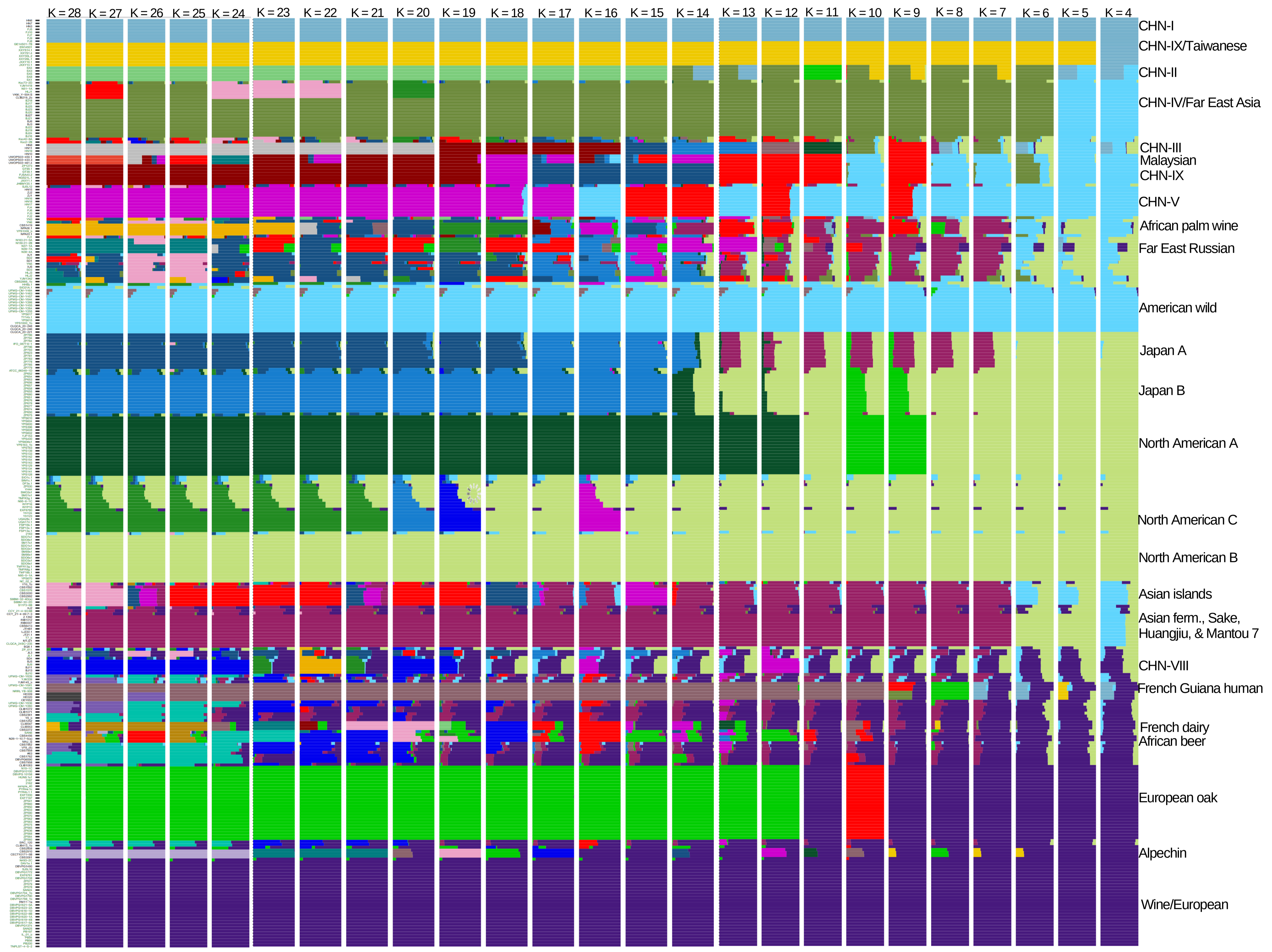
K = 28
K = 27
K = 26
K = 25
K = 24
K = 23
K = 22
K = 21
K = 20
K = 19
K = 18
K = 17
K = 16
K = 15
K = 14
K = 13
K = 12
K = 11
K = 10
K = 9
K = 8
K = 7
K = 6
K = 5
K = 4
CHN-I
CHN-IX/Taiwanese
CHN-II
CHN-IV/Far East Asia
CHN-III
Malaysian
CHN-IX
CHN-V
African palm wine
Far East Russian
American wild
Japan A
Japan B
North American A
North American C
North American B
Asian islands
Asian ferm., Sake,
Huangjiu, & Mantou 7
CHN-VIII
French Guiana human
French dairy
African beer
European oak
Alpechin
Wine/European

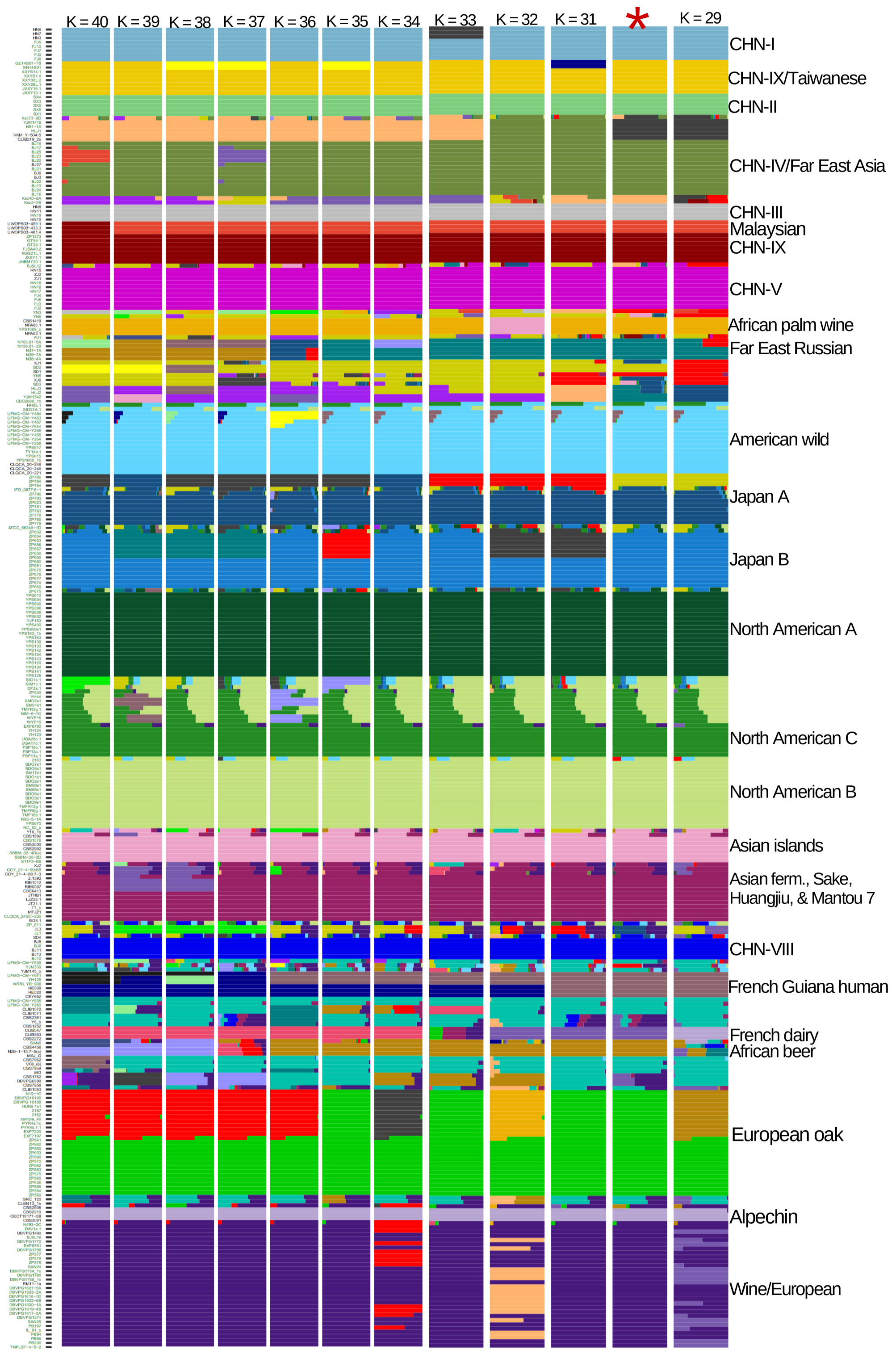
K = 40
K = 39
K = 38
K = 37
K = 36
K = 35
K = 34
K = 33
K = 32
K = 31
*
K = 29
CHN-I
CHN-IX/Taiwanese
CHN-II
CHN-IV/Far East Asia
CHN-III
Malaysian
CHN-IX
CHN-V
African palm wine
Far East Russian
American wild
Japan A
Japan B
North American A
North American C
North American B
Asian islands
Asian ferm., Sake,
Huangjiu, & Mantou 7
CHN-VIII
French Guiana human
French dairy
African beer
European oak
Alpechin
Wine/European

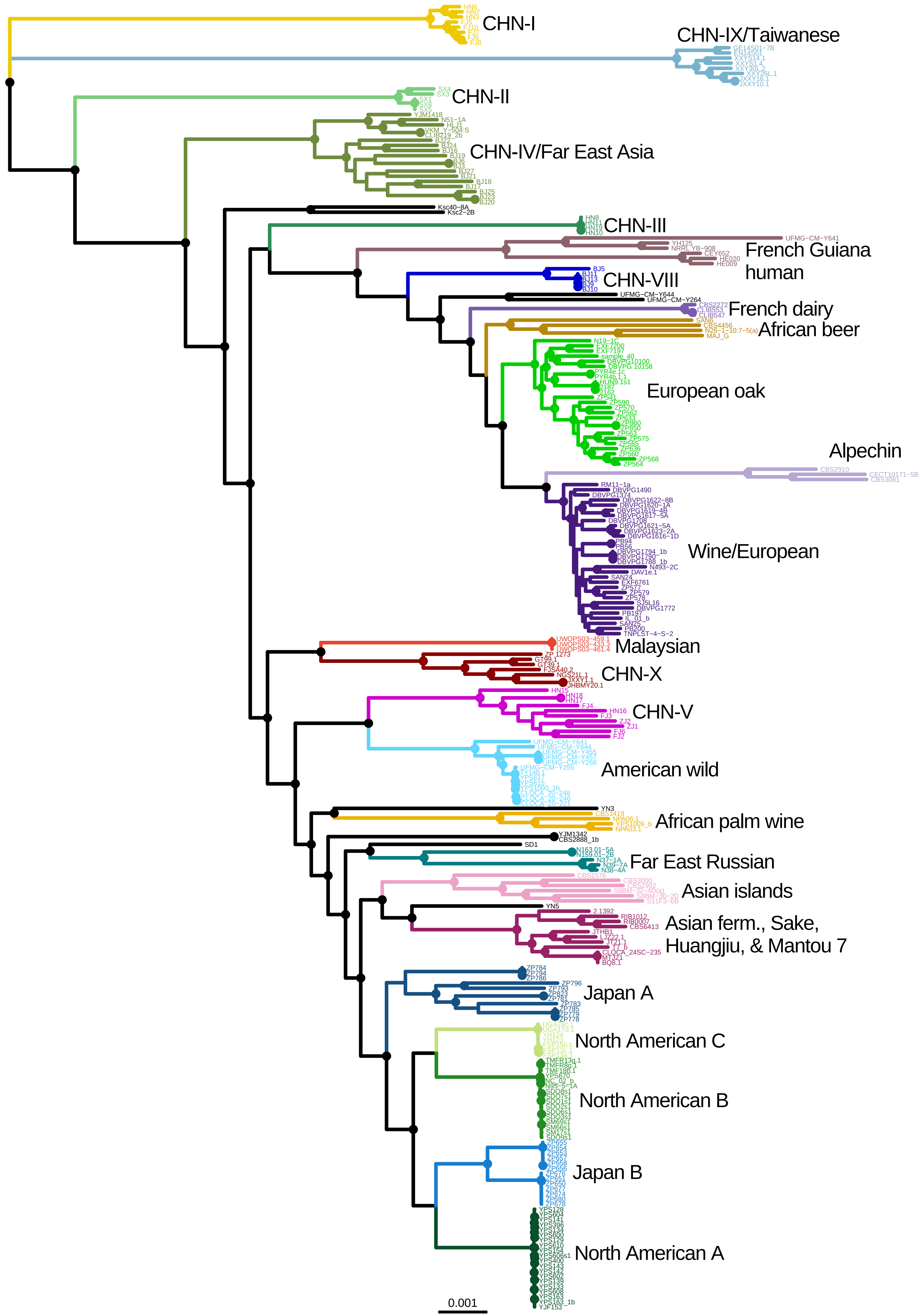
CHN-I
CHN-IX/Taiwanese
CHN-II
CHN-IV/Far East Asia
CHN-III
French Guiana human
CHN-VIII
French dairy
African beer
European oak
Alpechin
Wine/European
Malaysian
CHN-X
CHN-V
American wild
African palm wine
Far East Russian
Asian islands
Asian ferm., Sake, Huangjiu, & Mantou 7
Japan A
North American C
North American B
Japan B
North American A
0.001

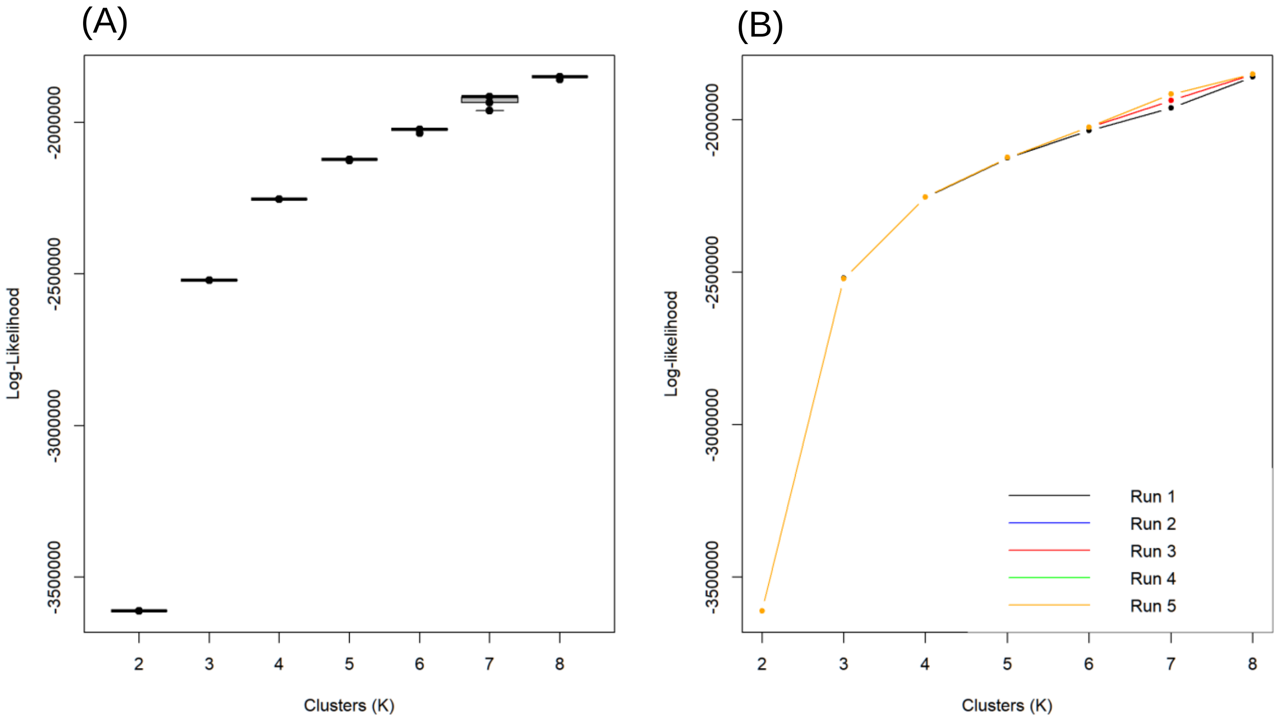
(A)
Log-Likelihood
-2000000
-2500000
-3000000
-3500000
2
3
4
5
6
7
8
Clusters (K)
(B)
Log-likelihood
-2000000
-2500000
-3000000
-3500000
Run 1
Run 2
Run 3
Run 4
Run 5
2
3
4
5
6
7
8
Clusters (K)

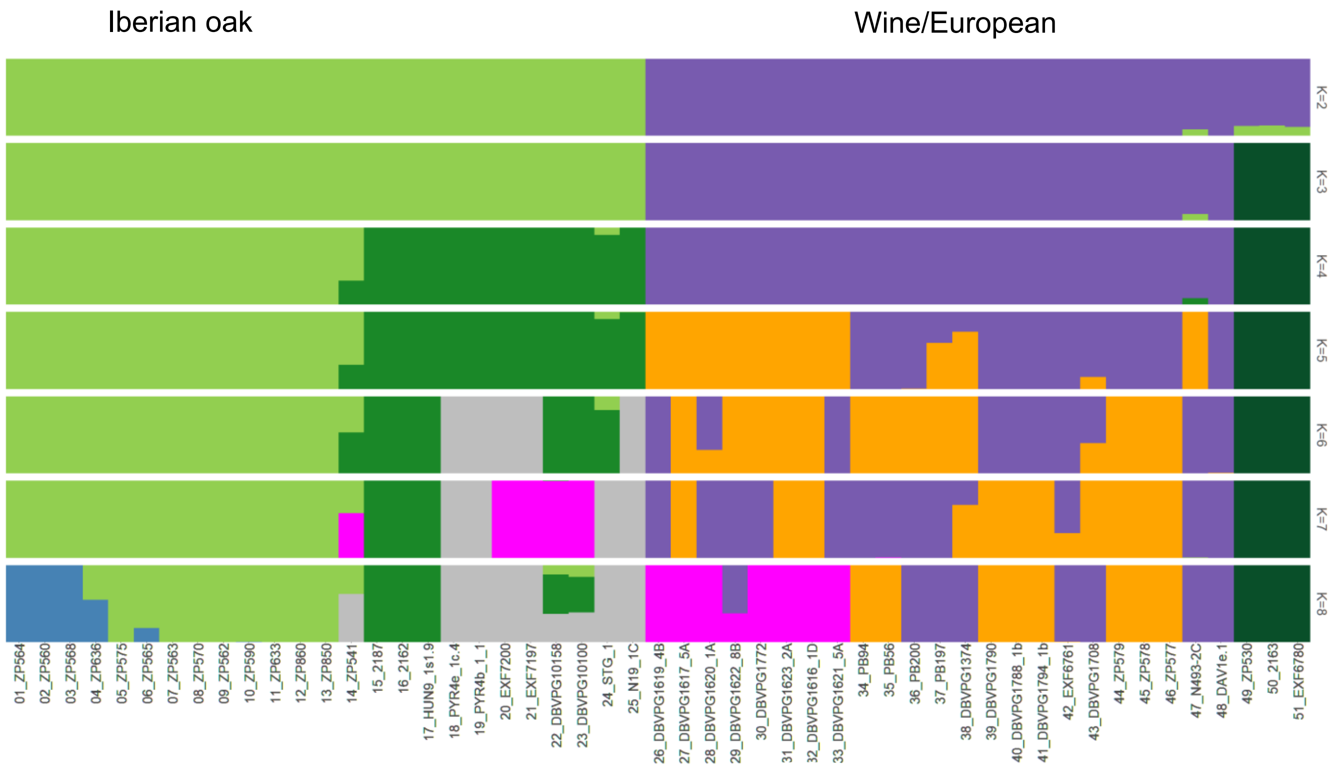

Iberian oak
Wine/European
K=2
K=3
K=4
K=5
K=6
K=7
K=8
01_ZP564
02_ZP560
03_ZP568
04_ZP636
05_ZP575
06_ZP565
07_ZP563
08_ZP570
09_ZP562
10_ZP590
11_ZP633
12_ZP860
13_ZP850
14_ZP541
15_2187
16_2162
17_HUN9_1s1.9
18_PYR4e_1c.4
19_PYR4b_1_1
20_EXF7200
21_EXF7197
22_DBVPG10158
23_DBVPG10100
24_STG_1
25_N19_1C
26_DBVPG1619_4B
27_DBVPG1617_5A
28_DBVPG1620_1A
29_DBVPG1622_8B
30_DBVPG1772
31_DBVPG1623_2A
32_DBVPG1616_1D
33_DBVPG1621_5A
34_PB94
35_PB56
36_PB200
37_PB197
38_DBVPG1374
39_DBVPG1790
40_DBVPG1788_1b
41_DBVPG1794_1b
42_EXF6761
43_DBVPG1708
44_ZP579
45_ZP578
46_ZP577
47_N493-2C
48_DAV1e.1
49_ZP530
50_2163
51_EXF6780

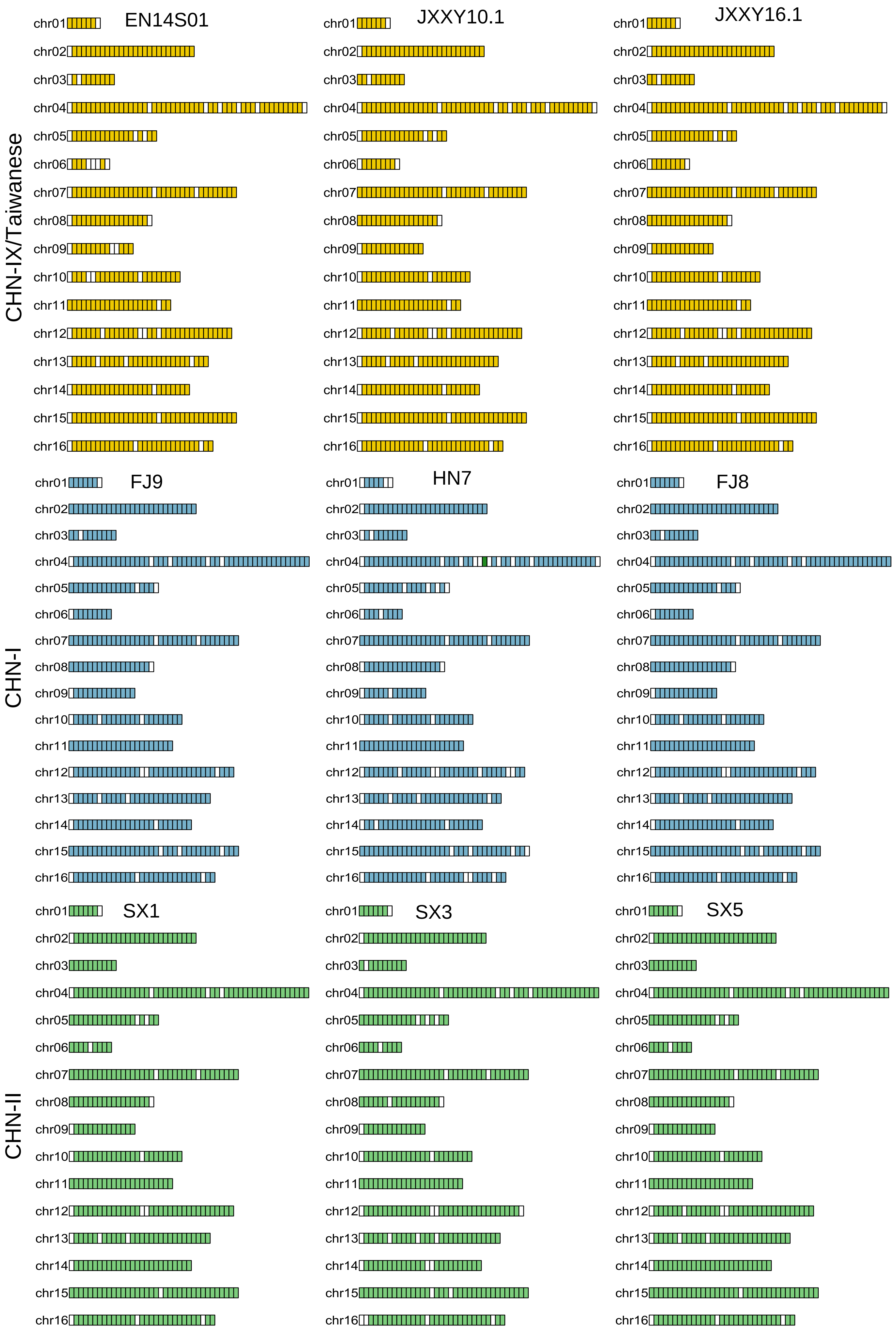

CHN-IX/Taiwanese
EN14S01
JXXY10.1
JXXY16.1
CHN-I
FJ9
HN7
FJ8
CHN-II
SX1
SX3
SX5
chr01
chr02
chr03
chr04
chr05
chr06
chr07
chr08
chr09
chr10
chr11
chr12
chr13
chr14
chr15
chr16

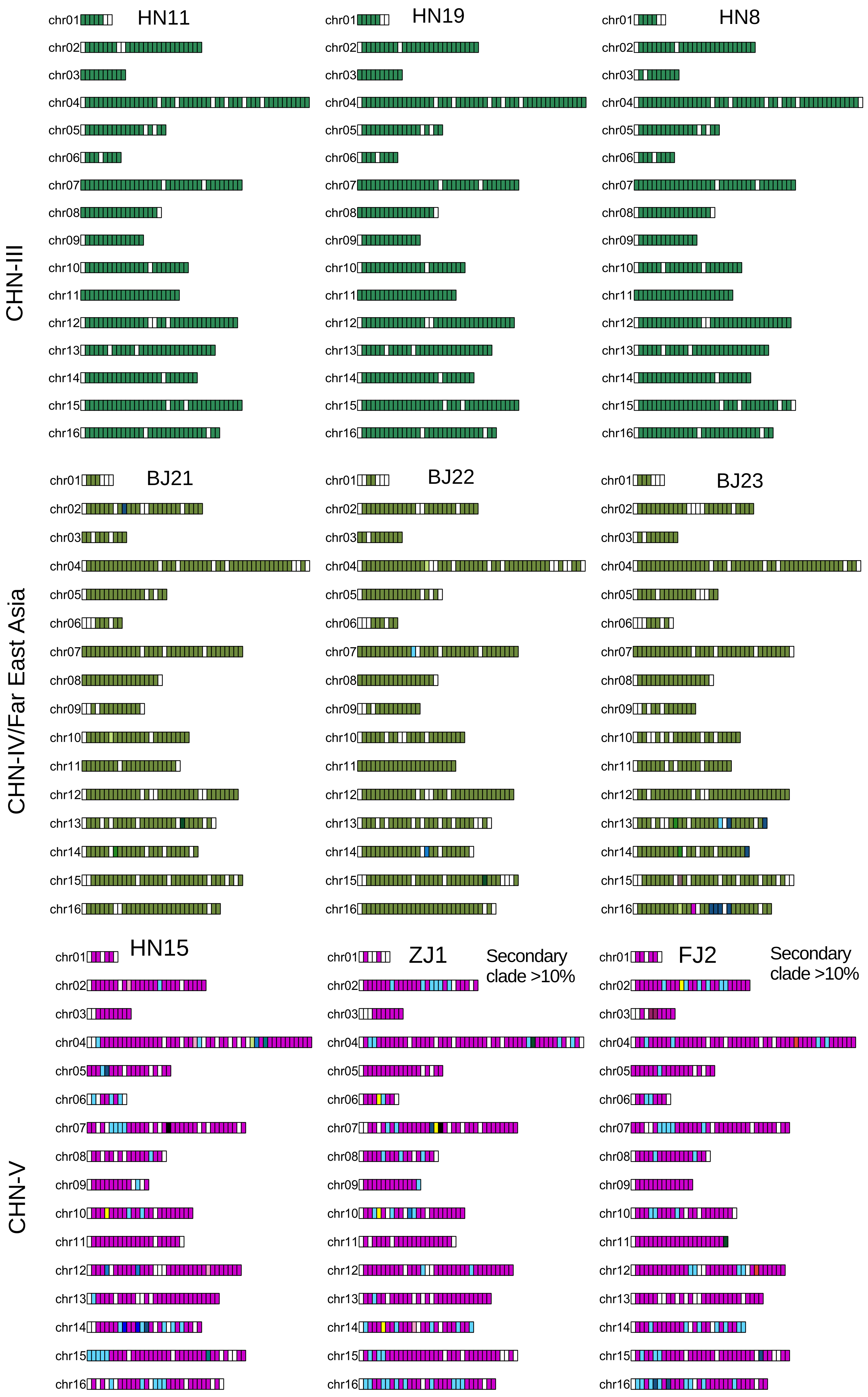

CHN-III
HN11
HN19
HN8
CHN-IV/Far East Asia
BJ21
BJ22
BJ23
CHN-V
HN15
ZJ1
Secondary clade >10%
FJ2
Secondary clade >10%
chr01
chr02
chr03
chr04
chr05
chr06
chr07
chr08
chr09
chr10
chr11
chr12
chr13
chr14
chr15
chr16

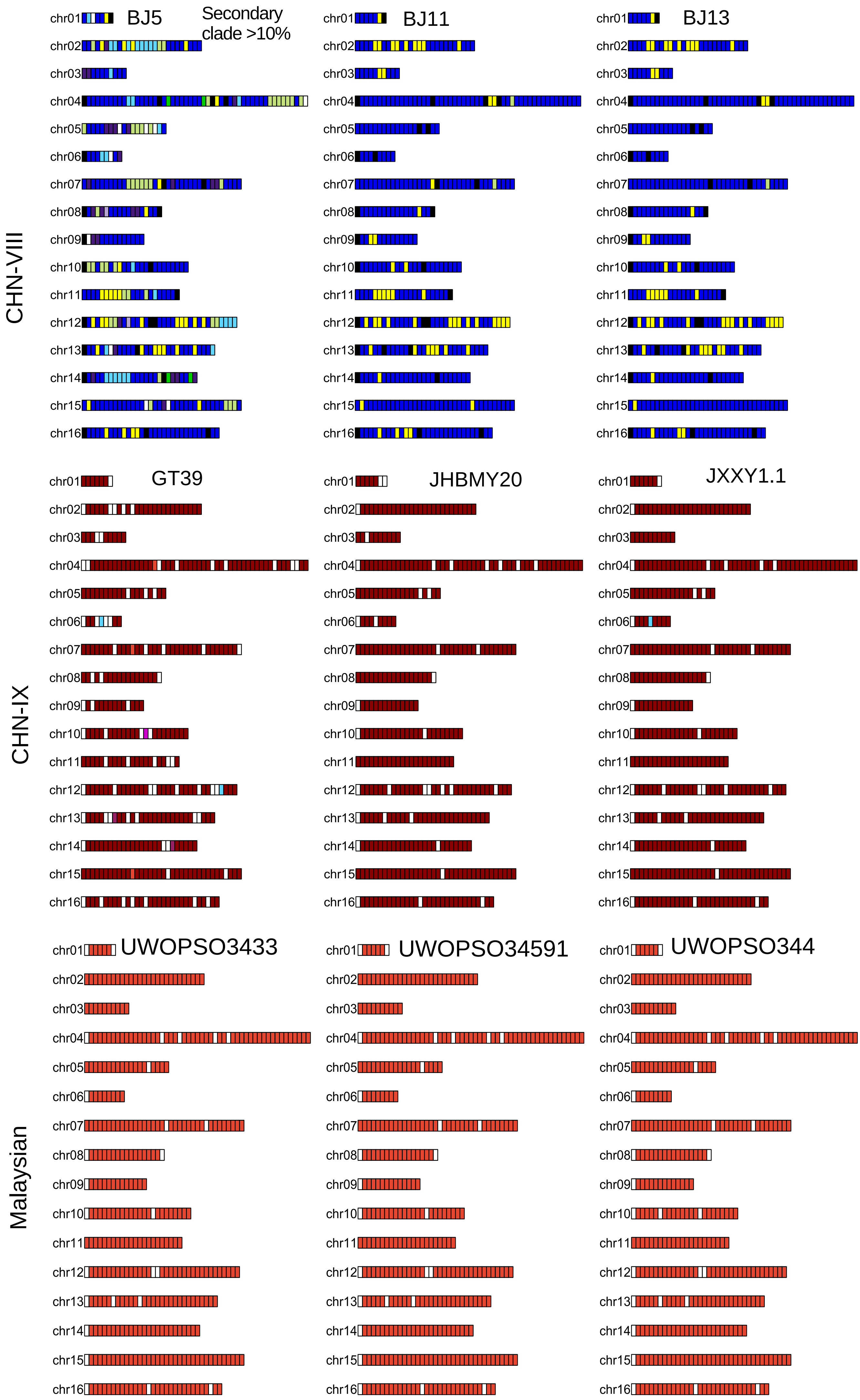
CHN-VIII
BJ5
Secondary clade >10%
BJ11
BJ13
CHN-IX
GT39
JHBMY20
JXXY1.1
Malaysian
UWOPSO3433
UWOPSO34591
UWOPSO344
chr01
chr02
chr03
chr04
chr05
chr06
chr07
chr08
chr09
chr10
chr11
chr12
chr13
chr14
chr15
chr16

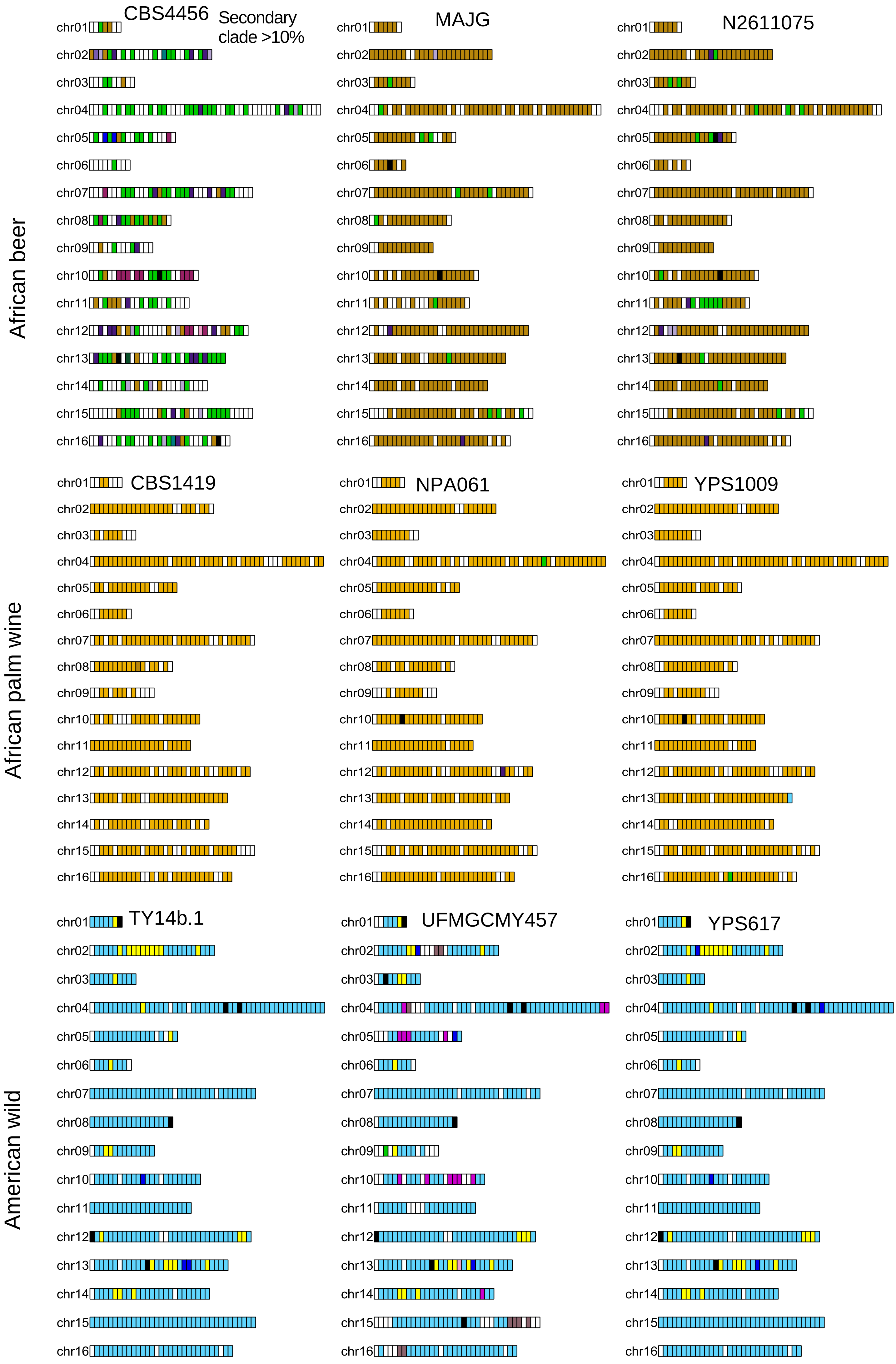
African beer
African palm wine
American wild
CBS4456
Secondary clade >10%
MAJG
N2611075
CBS1419
NPA061
YPS1009
TY14b.1
UFMGCMY457
YPS617
chr01
chr02
chr03
chr04
chr05
chr06
chr07
chr08
chr09
chr10
chr11
chr12
chr13
chr14
chr15
chr16

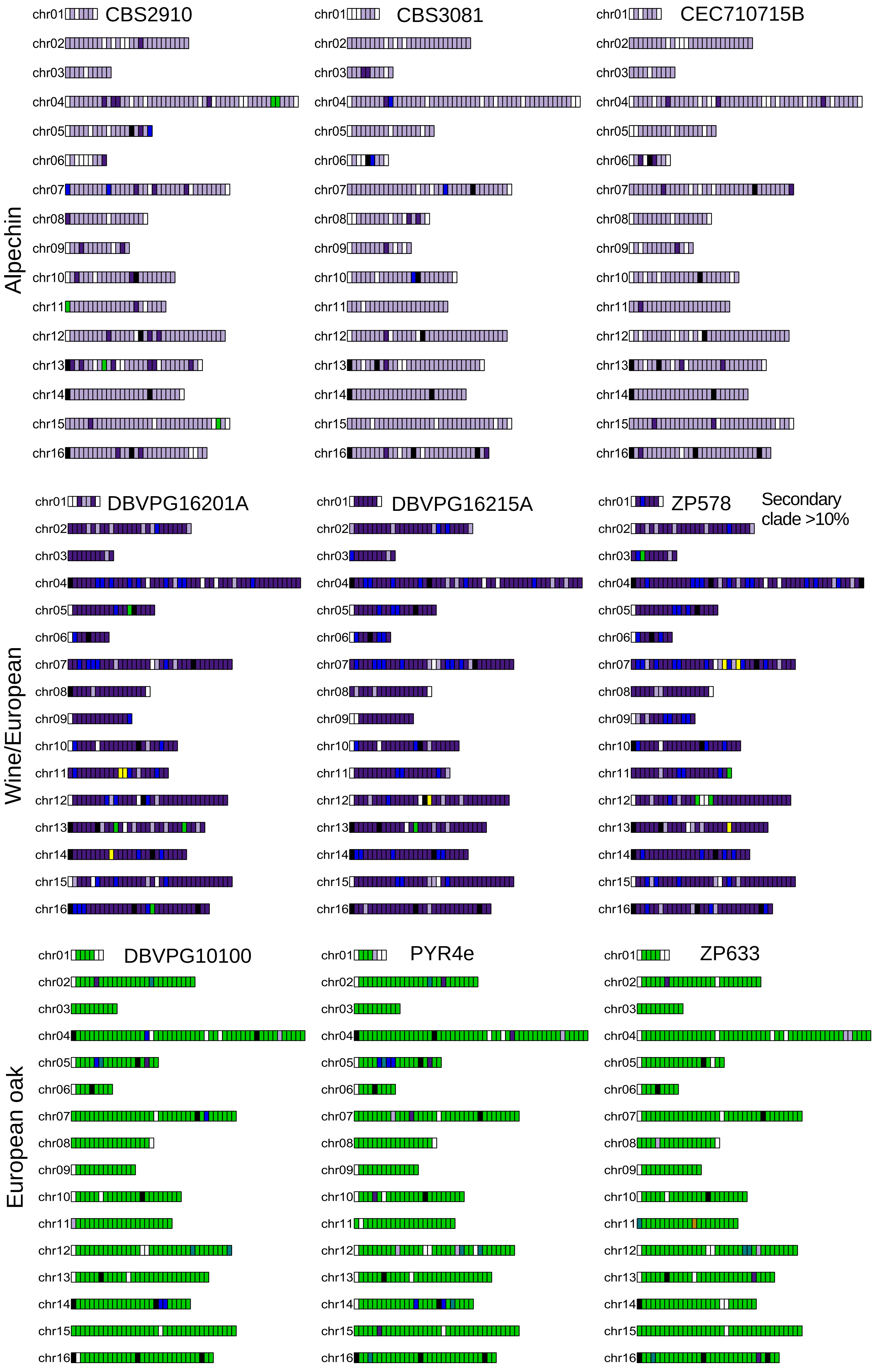
Alpechin
CBS2910
CBS3081
CEC710715B
Wine/European
DBVPG16201A
DBVPG16215A
ZP578
Secondary clade >10%
European oak
DBVPG10100
PYR4e
ZP633
chr01
chr02
chr03
chr04
chr05
chr06
chr07
chr08
chr09
chr10
chr11
chr12
chr13
chr14
chr15
chr16

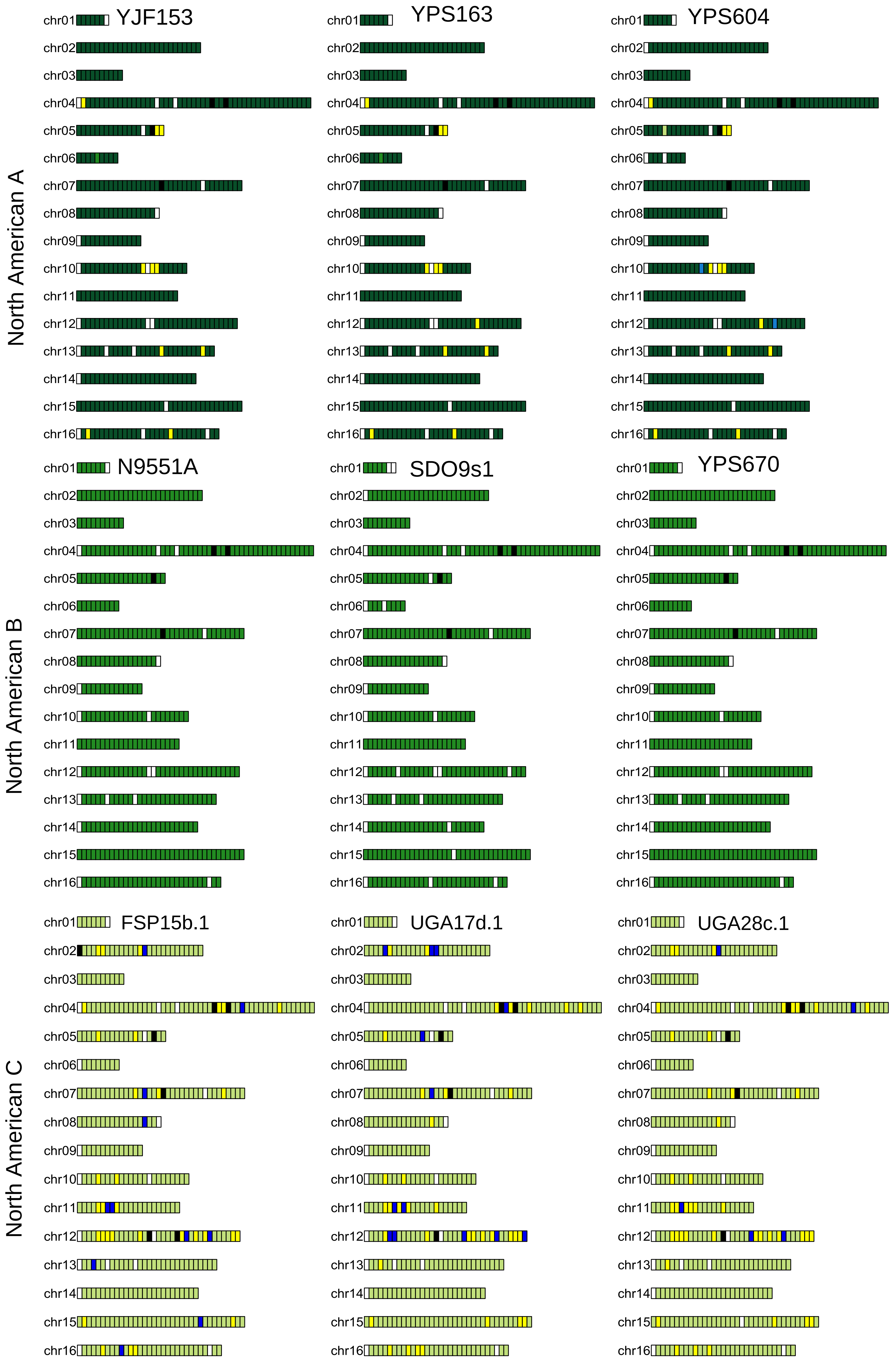
North American A
YJF153
YPS163
YPS604
North American B
N9551A
SDO9s1
YPS670
North American C
FSP15b.1
UGA17d.1
UGA28c.1
chr01
chr02
chr03
chr04
chr05
chr06
chr07
chr08
chr09
chr10
chr11
chr12
chr13
chr14
chr15
chr16

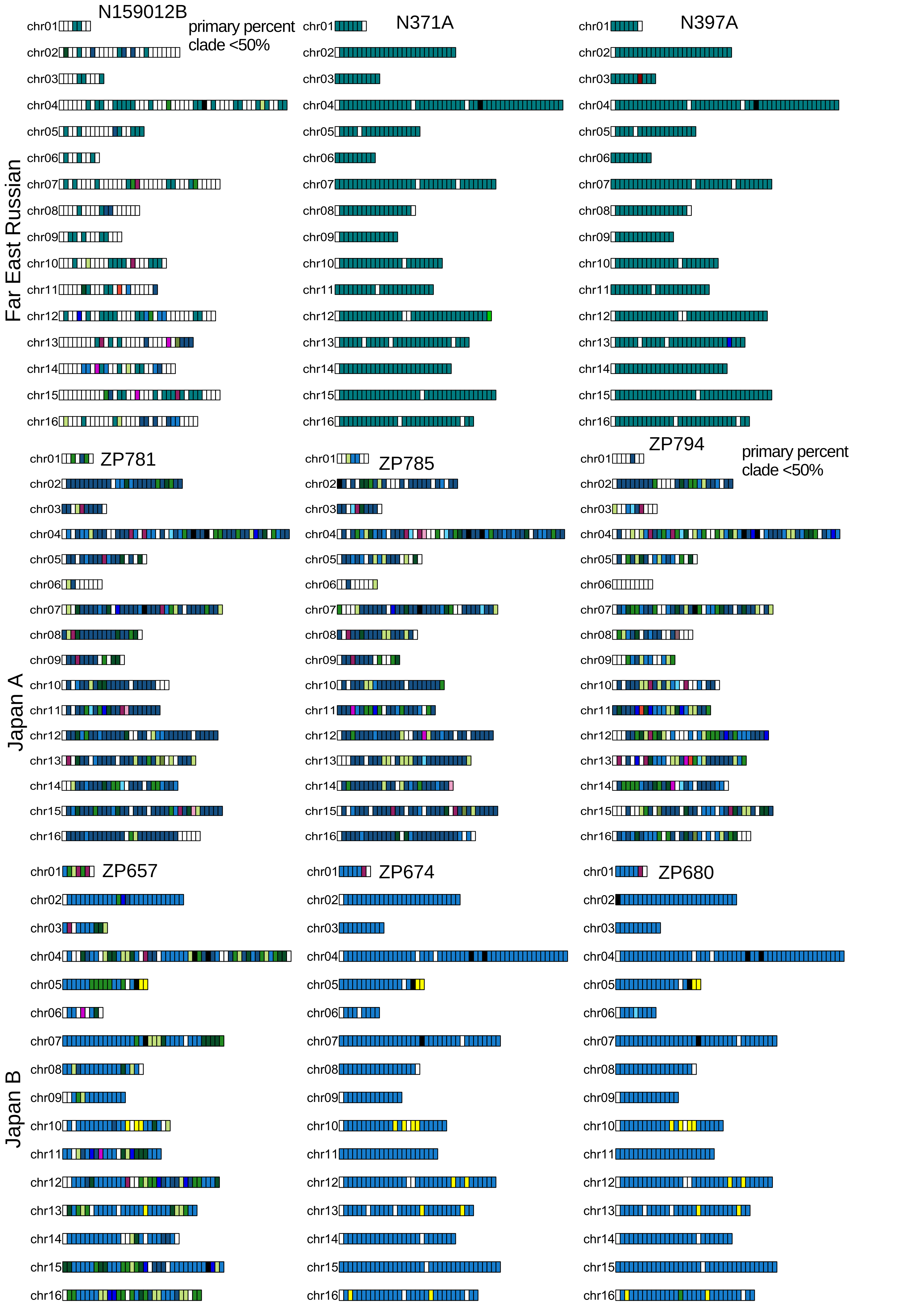
Far East Russian
N159012B
primary percent clade <50%
N371A
N397A
Japan A
ZP781
ZP785
ZP794
primary percent clade <50%
Japan B
ZP657
ZP674
ZP680
chr01
chr02
chr03
chr04
chr05
chr06
chr07
chr08
chr09
chr10
chr11
chr12
chr13
chr14
chr15
chr16

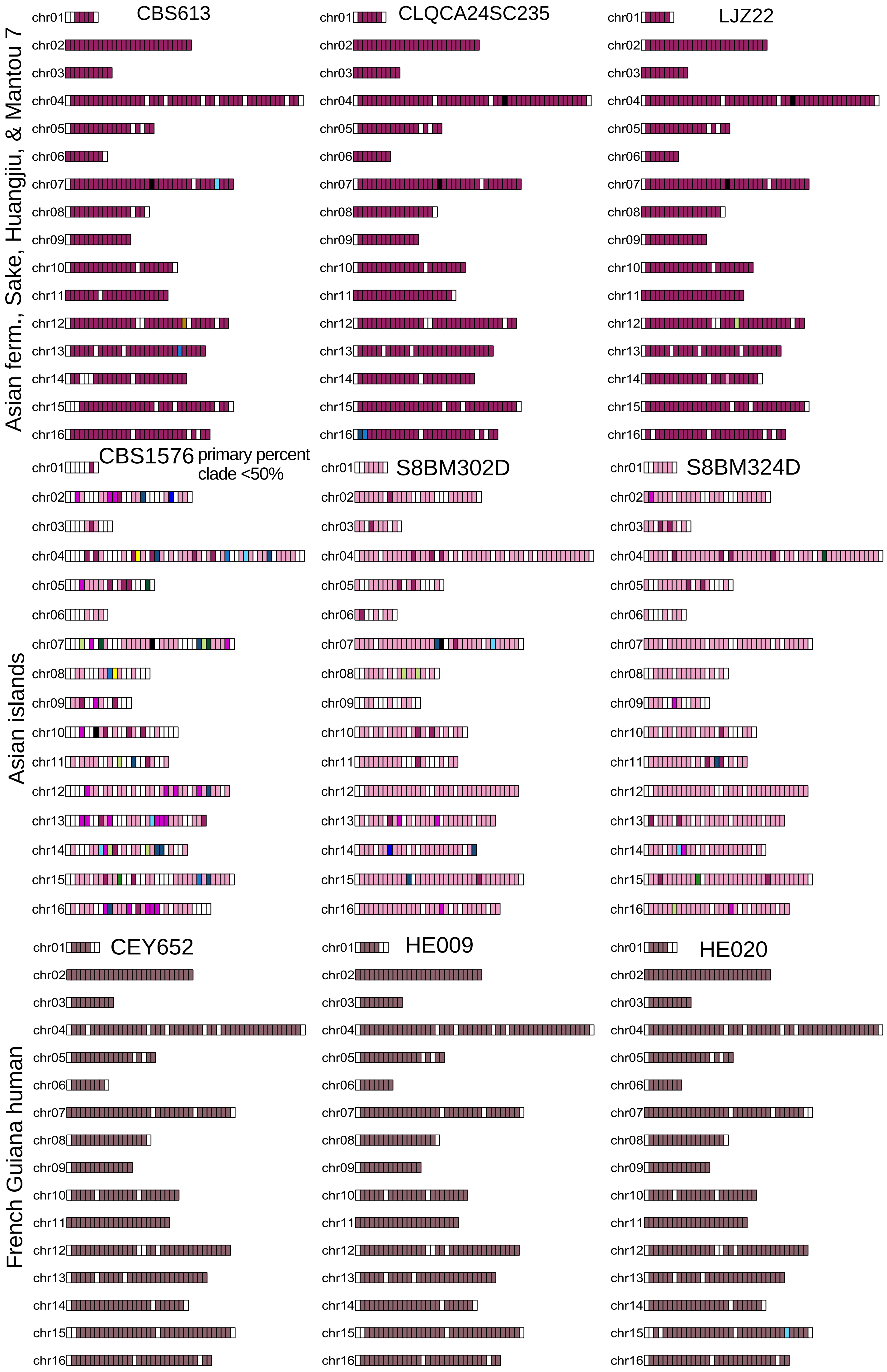
Asian ferm., Sake, Huangjiu, & Mantou 7
CBS613
CLQCA24SC235
LJZ22
CBS1576
primary percent clade <50%
S8BM302D
S8BM324D
Asian islands
CEY652
HE009
HE020
French Guiana human
chr01
chr02
chr03
chr04
chr05
chr06
chr07
chr08
chr09
chr10
chr11
chr12
chr13
chr14
chr15
chr16

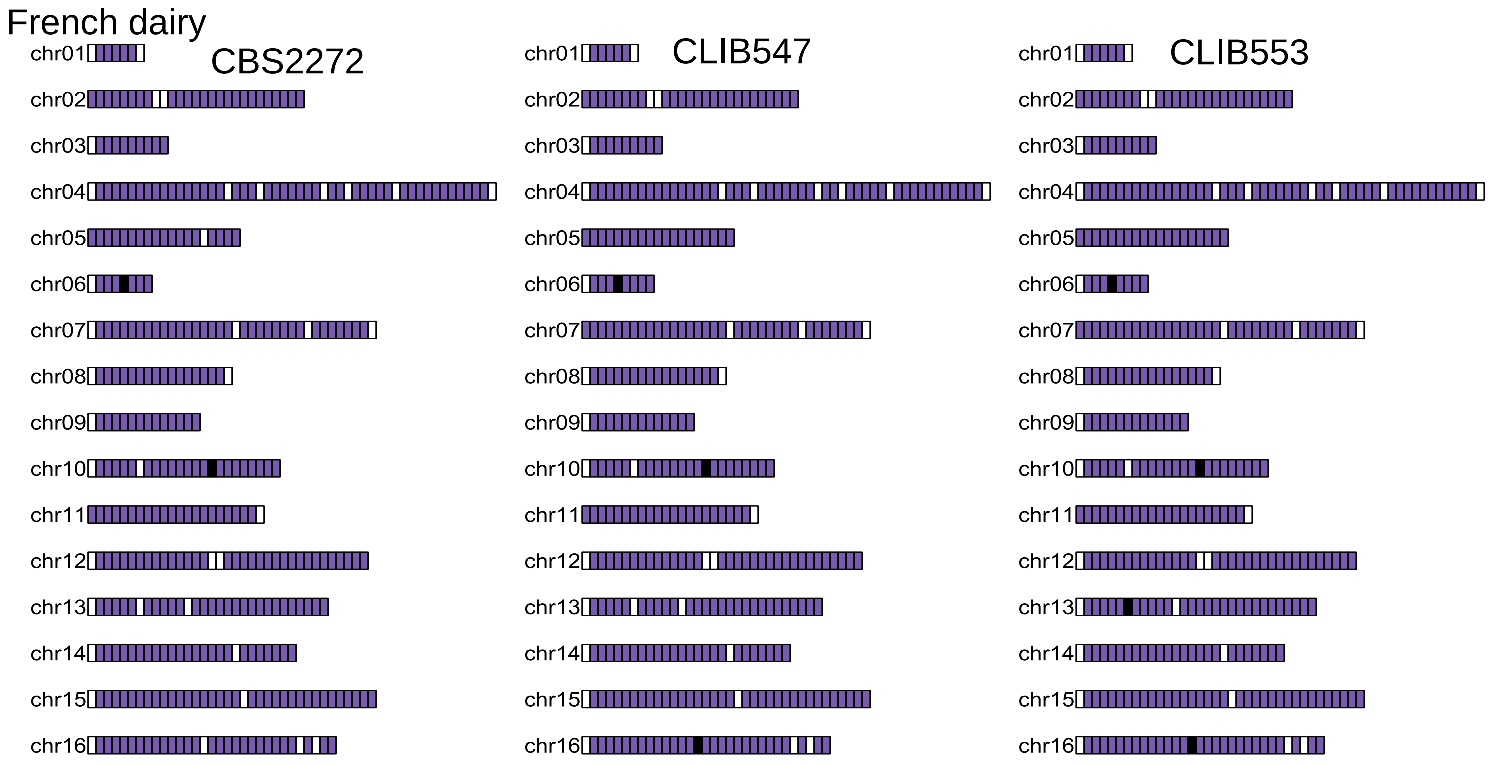
French dairy
CBS2272
CLIB547
CLIB553
chr01
chr02
chr03
chr04
chr05
chr06
chr07
chr08
chr09
chr10
chr11
chr12
chr13
chr14
chr15
chr16

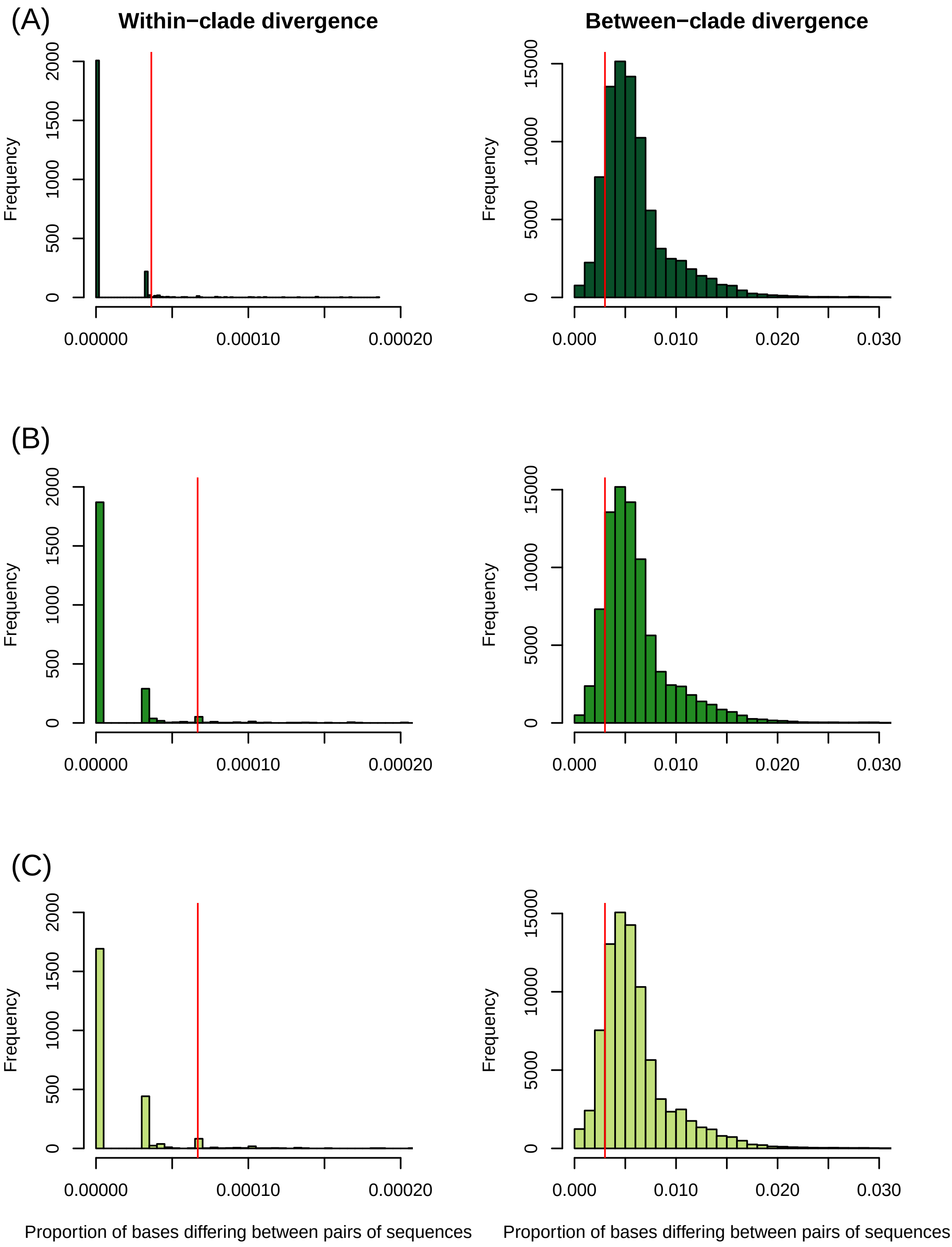

(A)
Within−clade divergence
Between−clade divergence
Frequency
0
500
1000
1500
2000
0.00000
0.00010
0.00020
5000
10000
15000
0.000
0.010
0.020
0.030
(B)
(C)
Proportion of bases differing between pairs of sequences
Proportion of bases differing between pairs of sequences

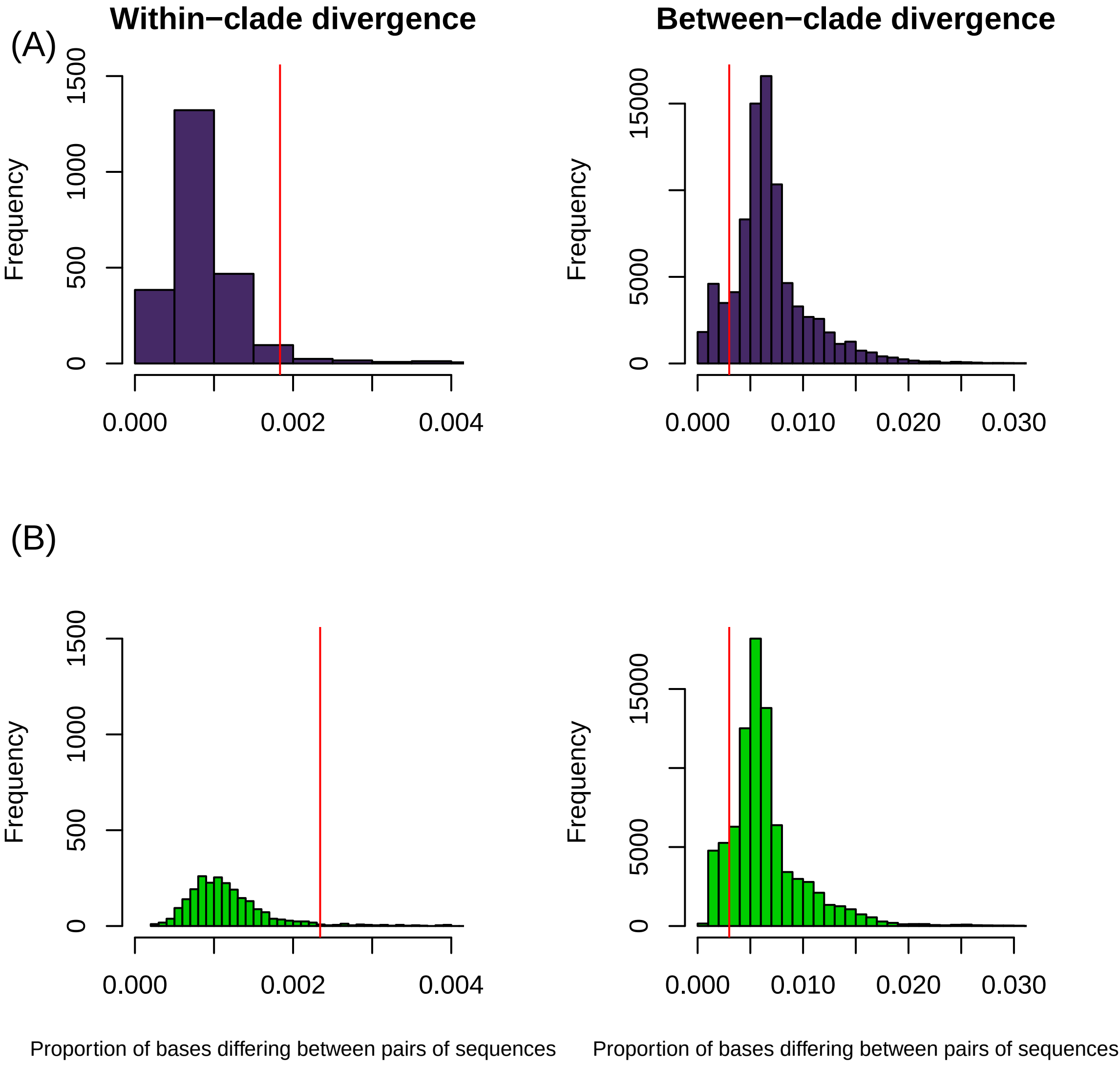
Within−clade divergence
Between−clade divergence
(A)
(B)
Frequency
0
500
1000
1500
0
5000
15000
0.000
0.002
0.004
0.000
0.010
0.020
0.030
Proportion of bases differing between pairs of sequences
Proportion of bases differing between pairs of sequences

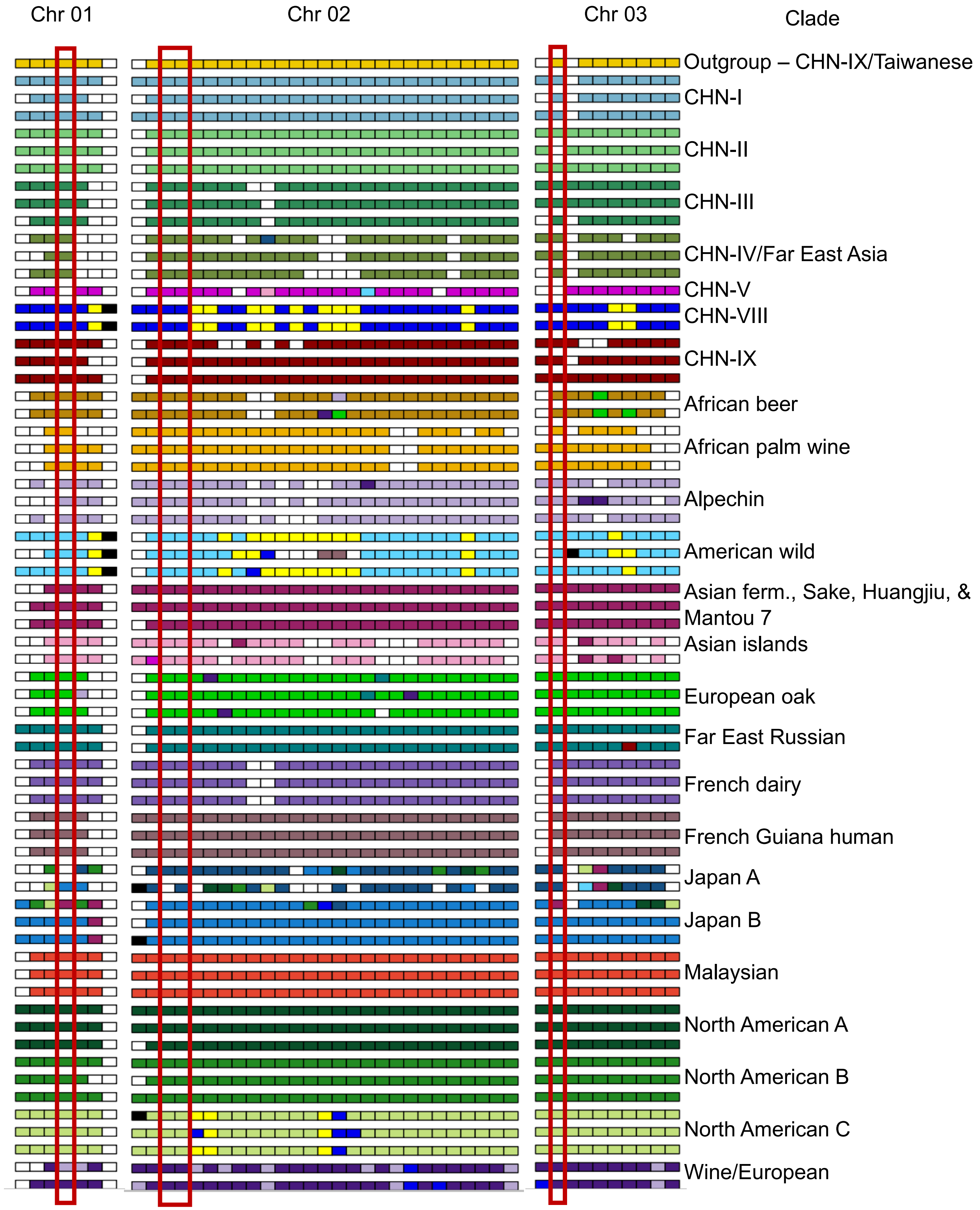
Chr 01
Chr 02
Chr 03
Clade
Outgroup – CHN-IX/Taiwanese
CHN-I
CHN-II
CHN-III
CHN-IV/Far East Asia
CHN-V
CHN-VIII
CHN-IX
African beer
African palm wine
Alpechin
American wild
Asian ferm., Sake, Huangjiu, &
Mantou 7
Asian islands
European oak
Far East Russian
French dairy
French Guiana human
Japan A
Japan B
Malaysian
North American A
North American B
North American C
Wine/European

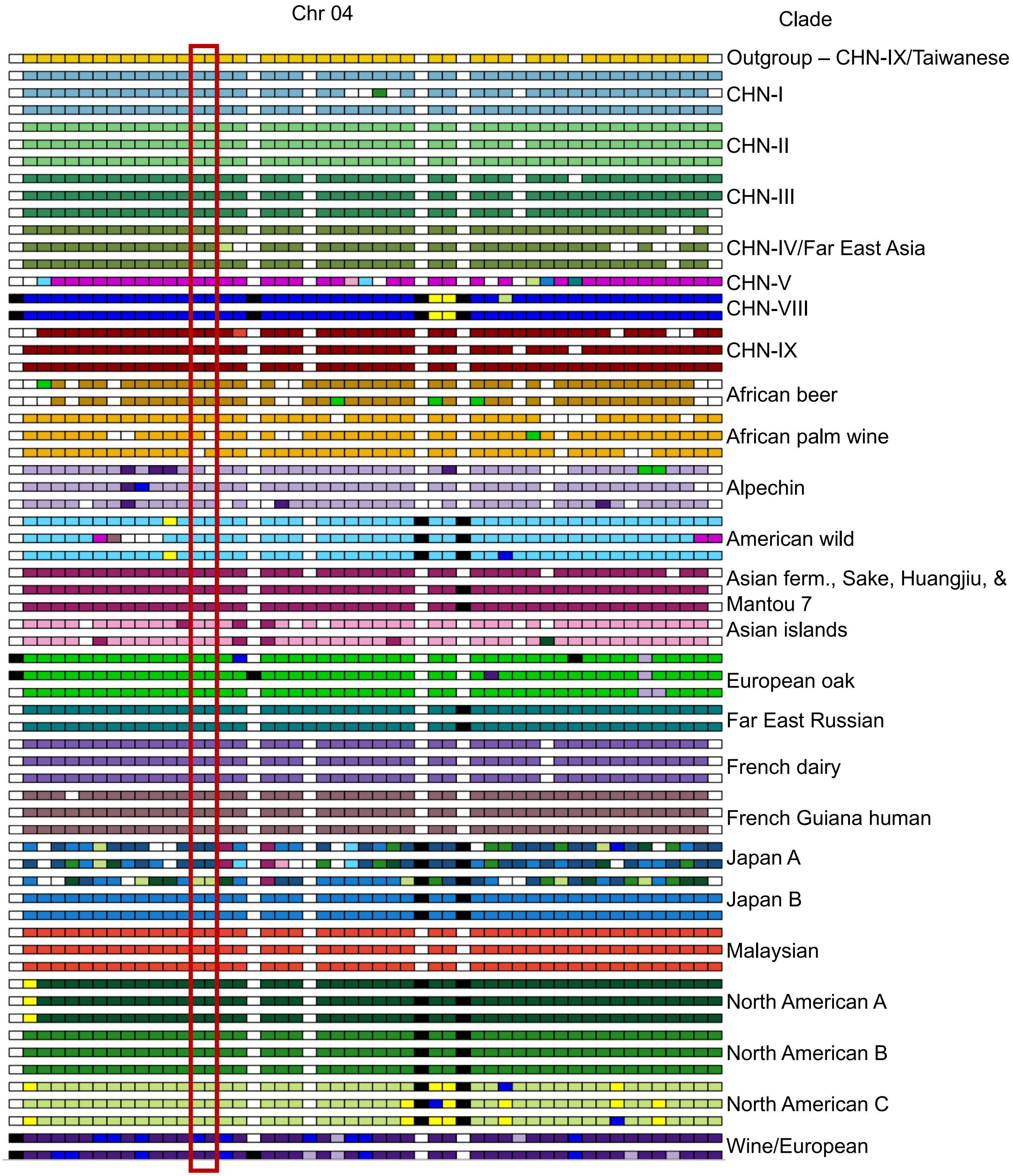
Chr 04
Clade
Outgroup – CHN-IX/Taiwanese
CHN-I
CHN-II
CHN-III
CHN-IV/Far East Asia
CHN-V
CHN-VIII
CHN-IX
African beer
African palm wine
Alpechin
American wild
Asian ferm., Sake, Huangjiu, & Mantou 7
Asian islands
European oak
Far East Russian
French dairy
French Guiana human
Japan A
Japan B
Malaysian
North American A
North American B
North American C
Wine/European

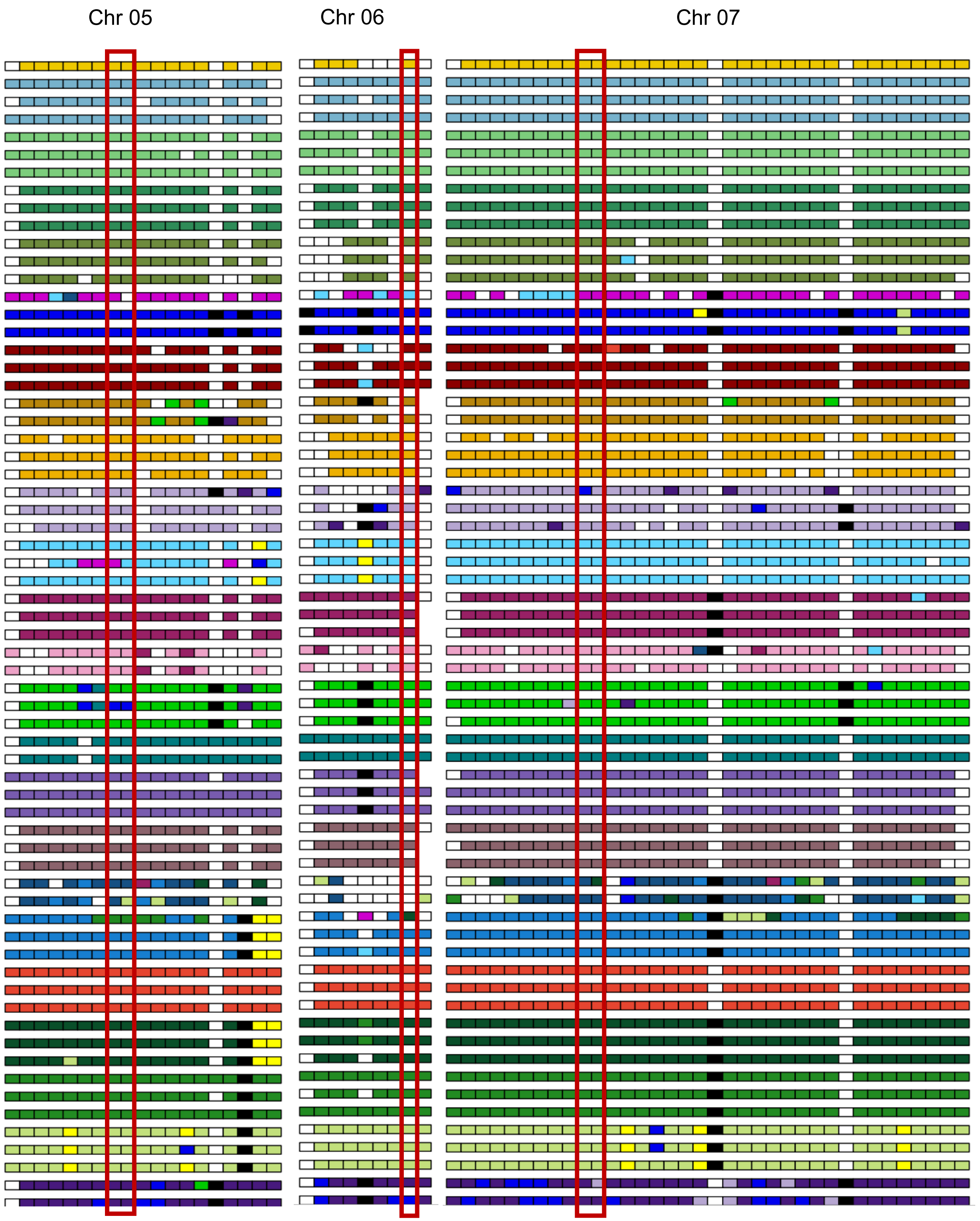
Chr 05
Chr 06
Chr 07

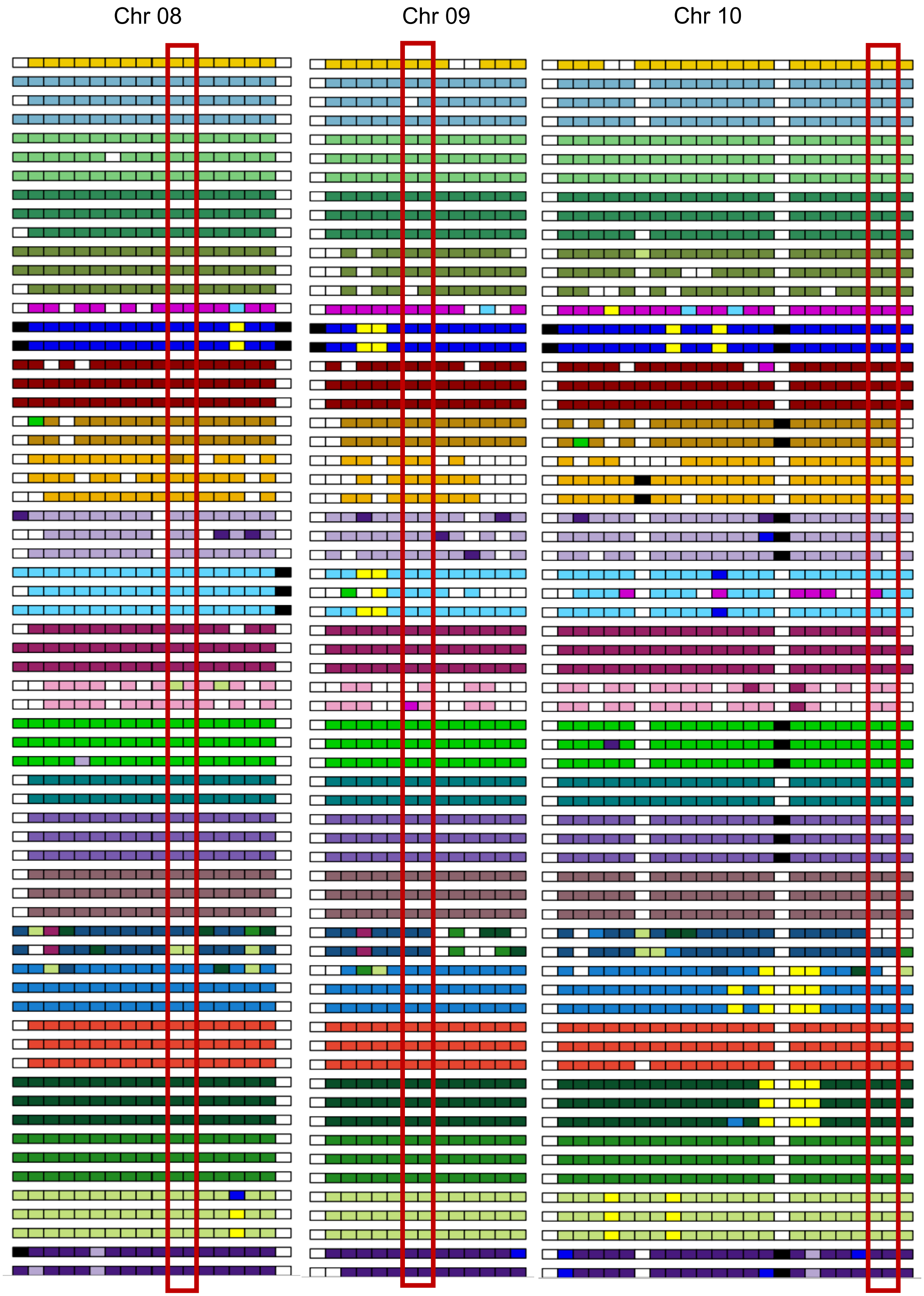
Chr 08
Chr 09
Chr 10

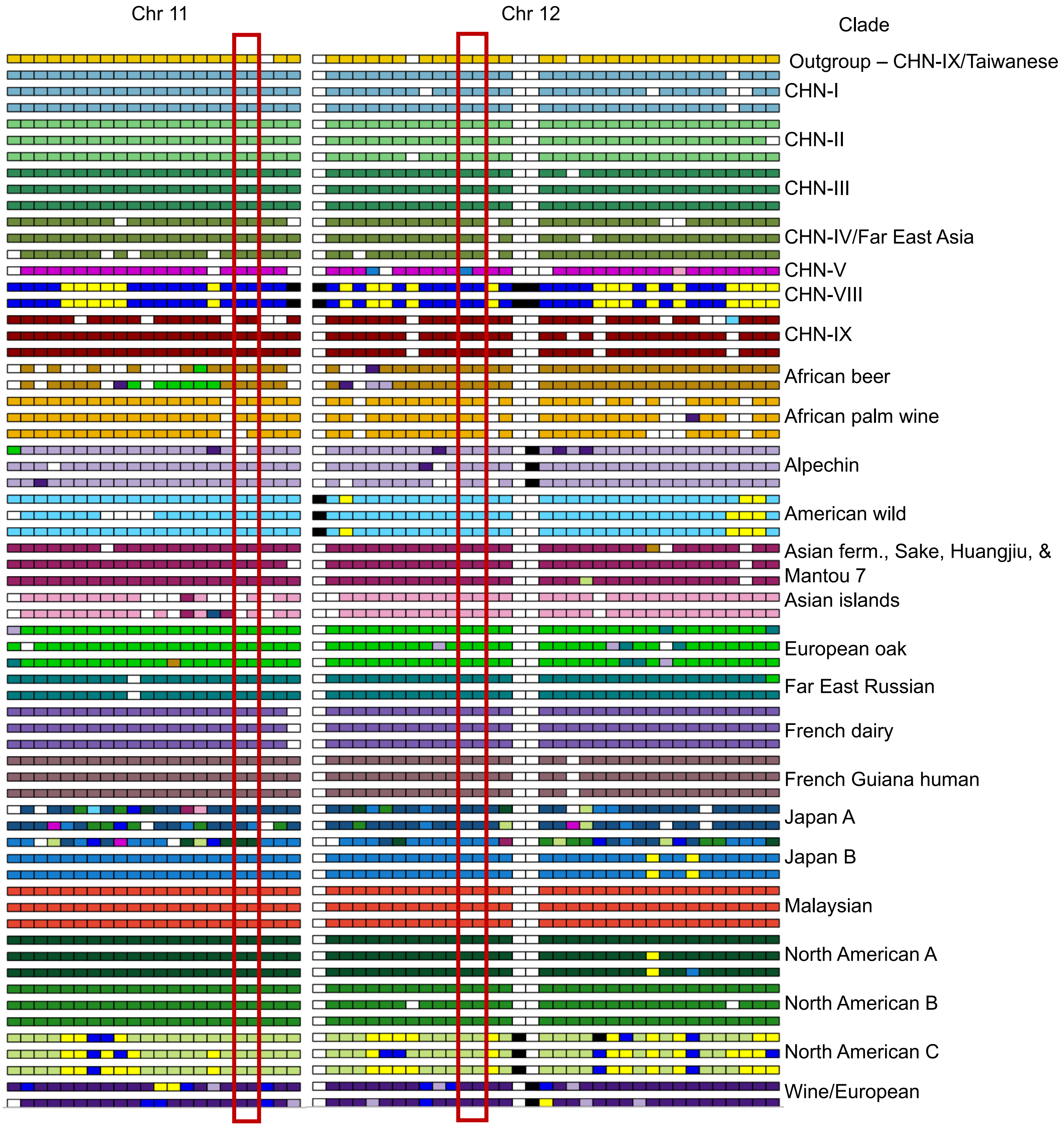
Chr 11
Chr 12
Clade
Outgroup – CHN-IX/Taiwanese
CHN-I
CHN-II
CHN-III
CHN-IV/Far East Asia
CHN-V
CHN-VIII
CHN-IX
African beer
African palm wine
Alpechin
American wild
Asian ferm., Sake, Huangjiu, &
Mantou 7
Asian islands
European oak
Far East Russian
French dairy
French Guiana human
Japan A
Japan B
Malaysian
North American A
North American B
North American C
Wine/European

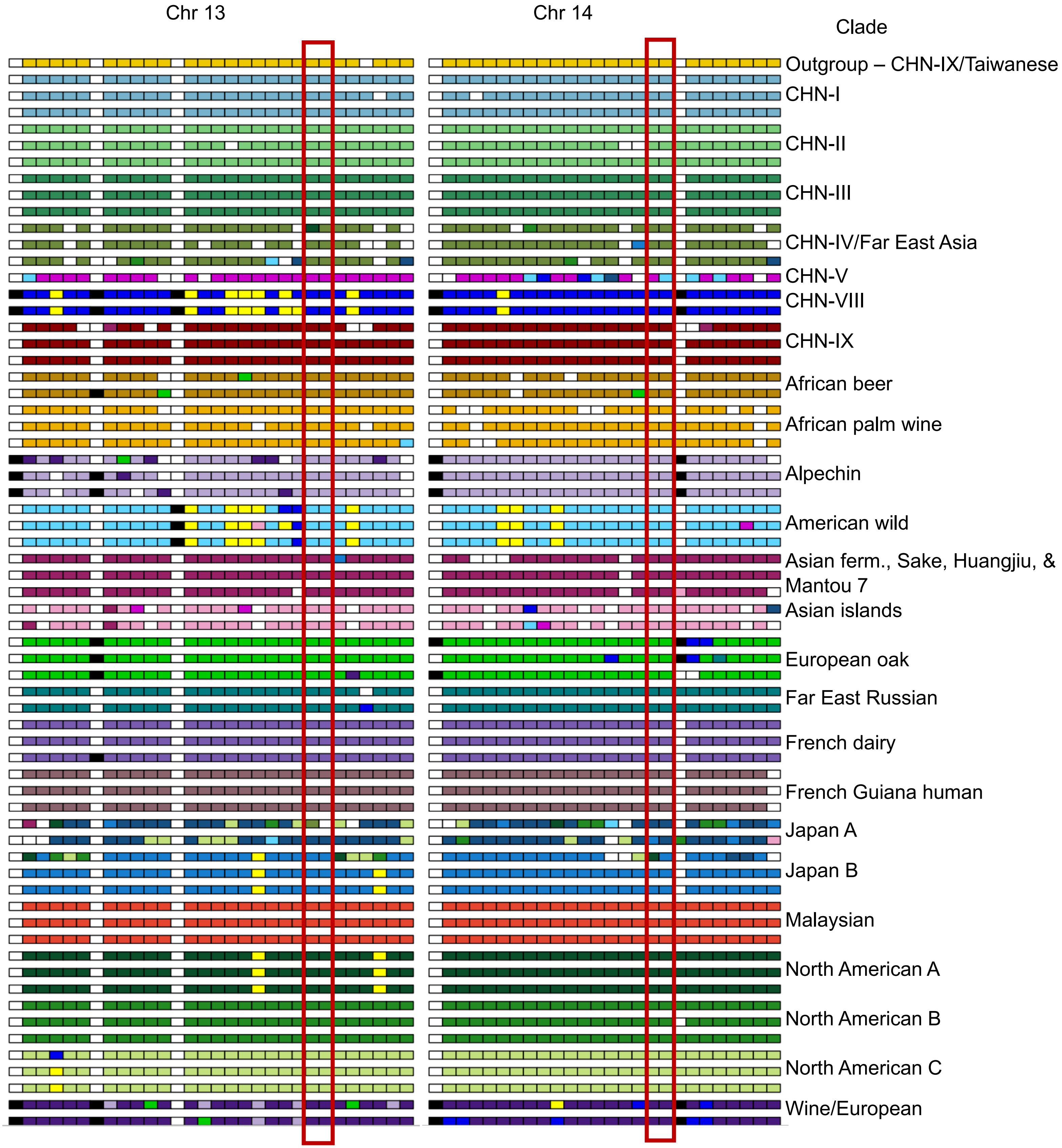

Chr 13
Chr 14
Clade
Outgroup – CHN-IX/Taiwanese
CHN-I
CHN-II
CHN-III
CHN-IV/Far East Asia
CHN-V
CHN-VIII
CHN-IX
African beer
African palm wine
Alpechin
American wild
Asian ferm., Sake, Huangjiu, & Mantou 7
Asian islands
European oak
Far East Russian
French dairy
French Guiana human
Japan A
Japan B
Malaysian
North American A
North American B
North American C
Wine/European

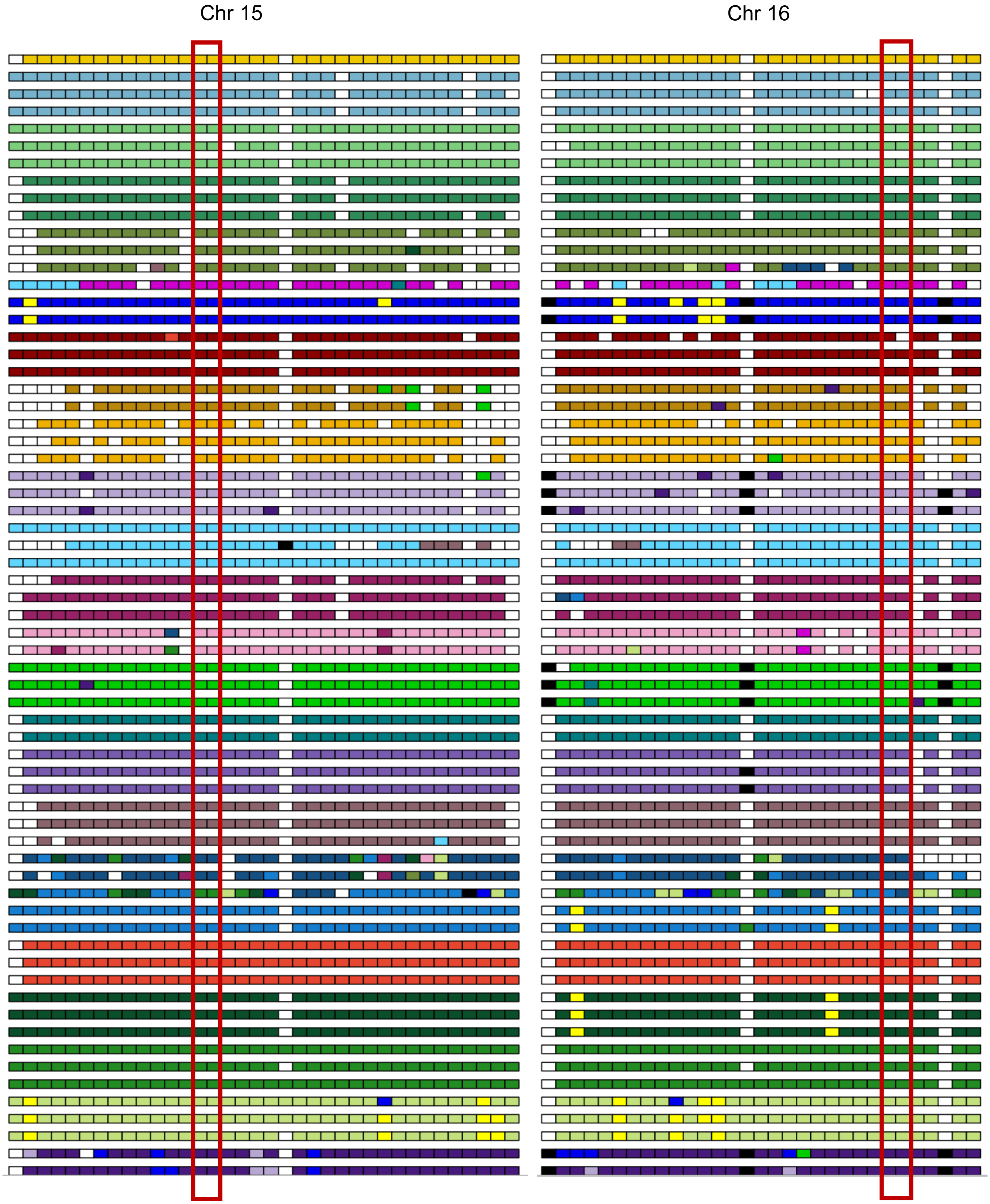
Chr 15
Chr 16

Chromosome 1

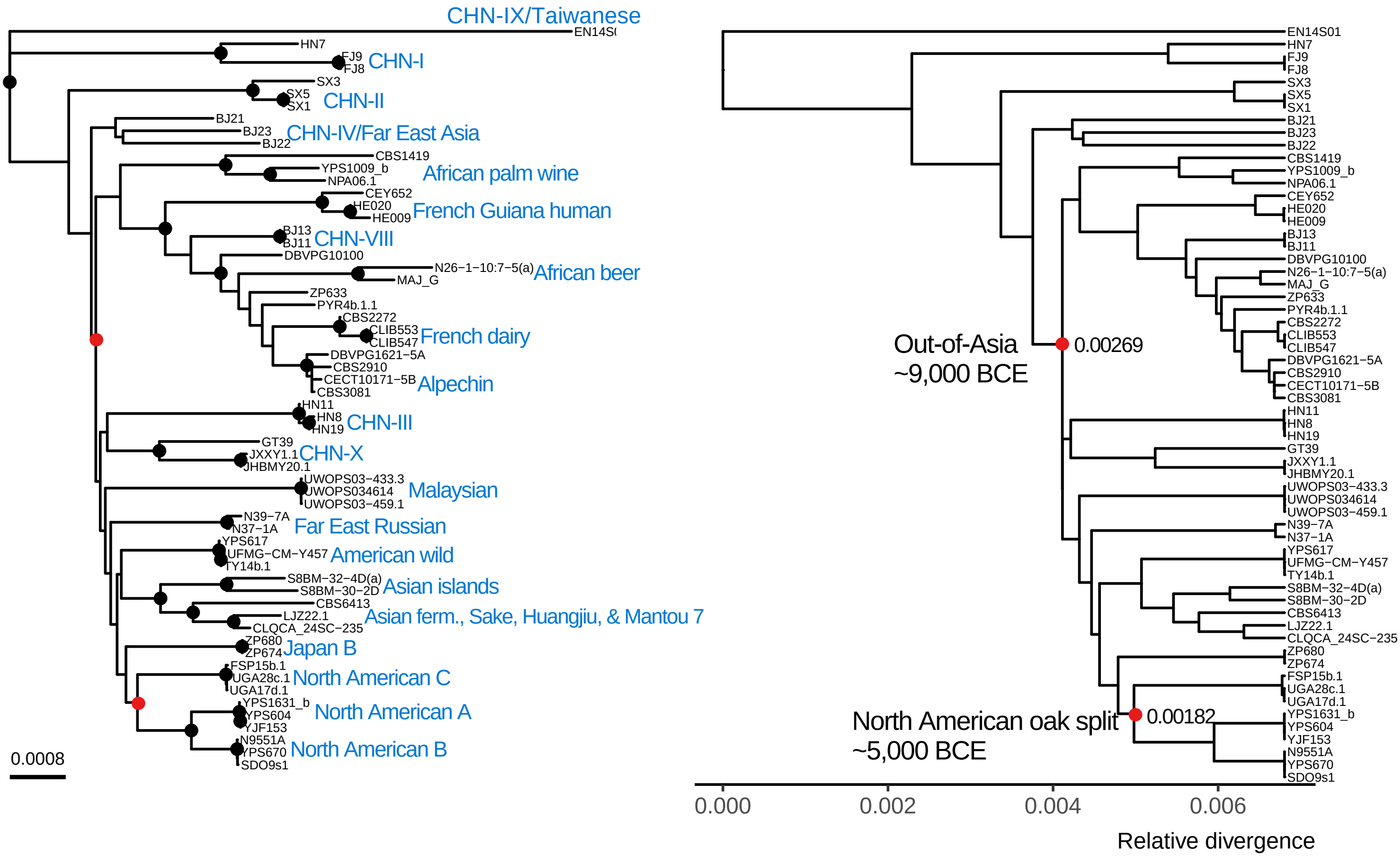


Chromosome 2

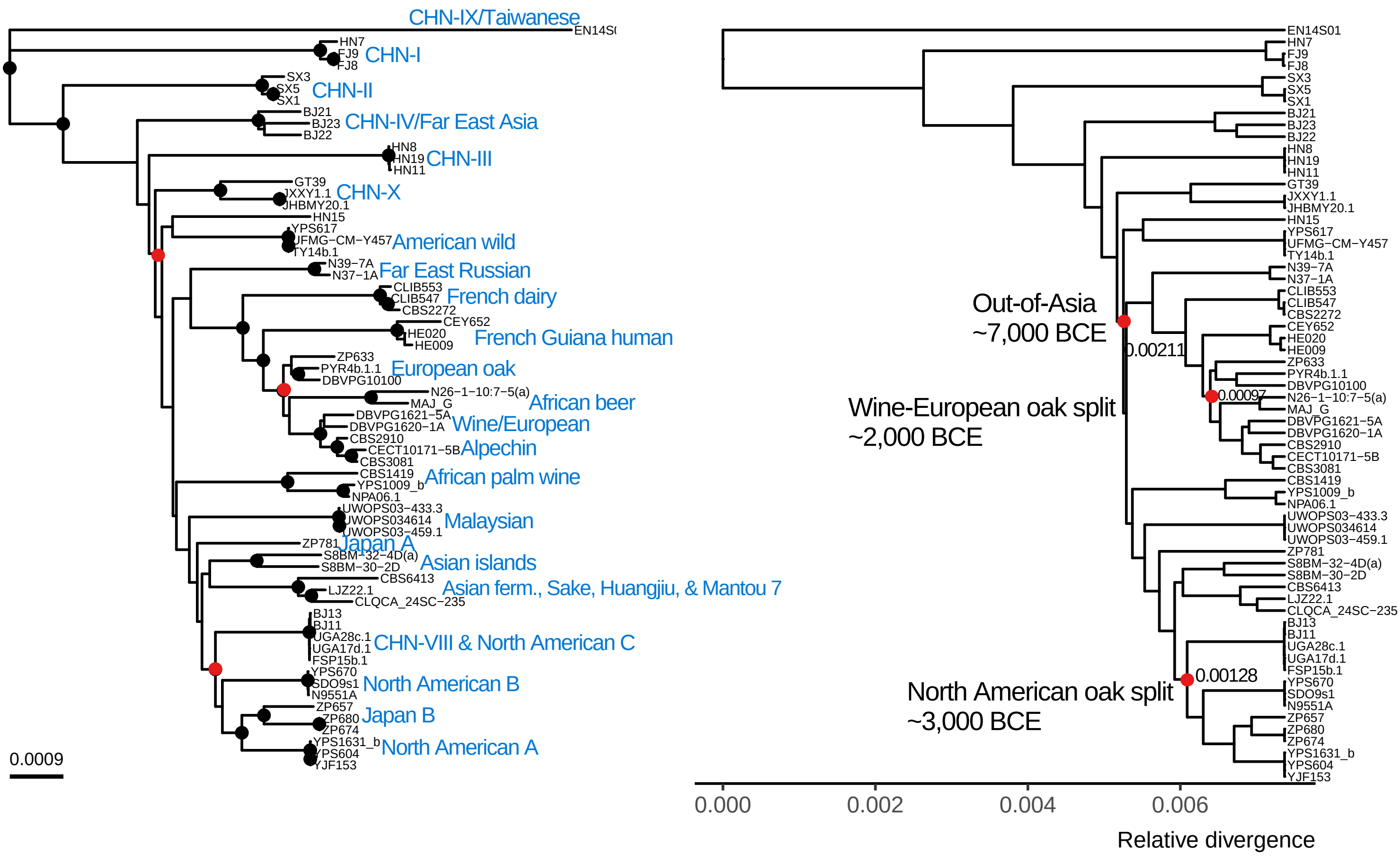

Chromosome 3

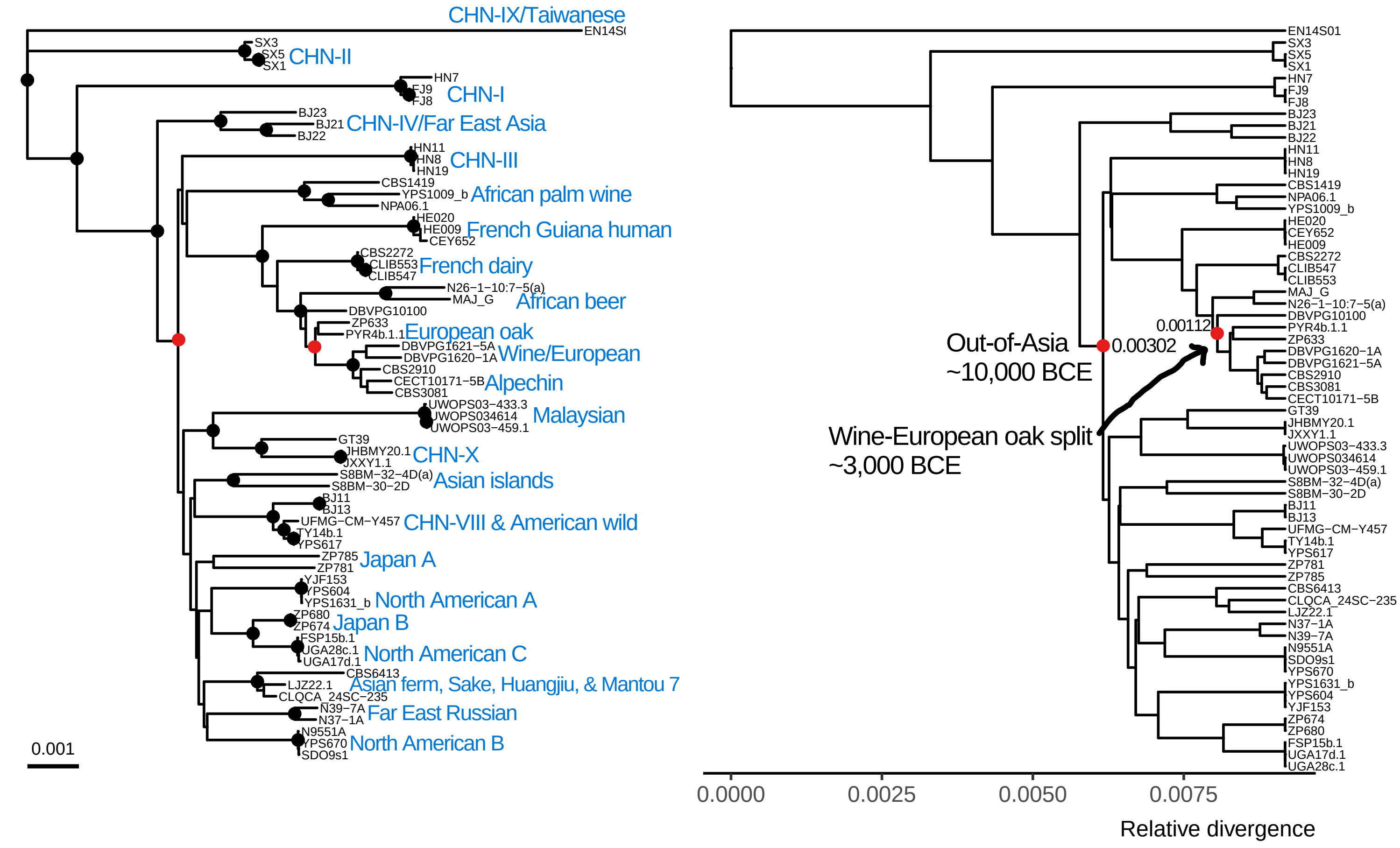


Chromosome 4

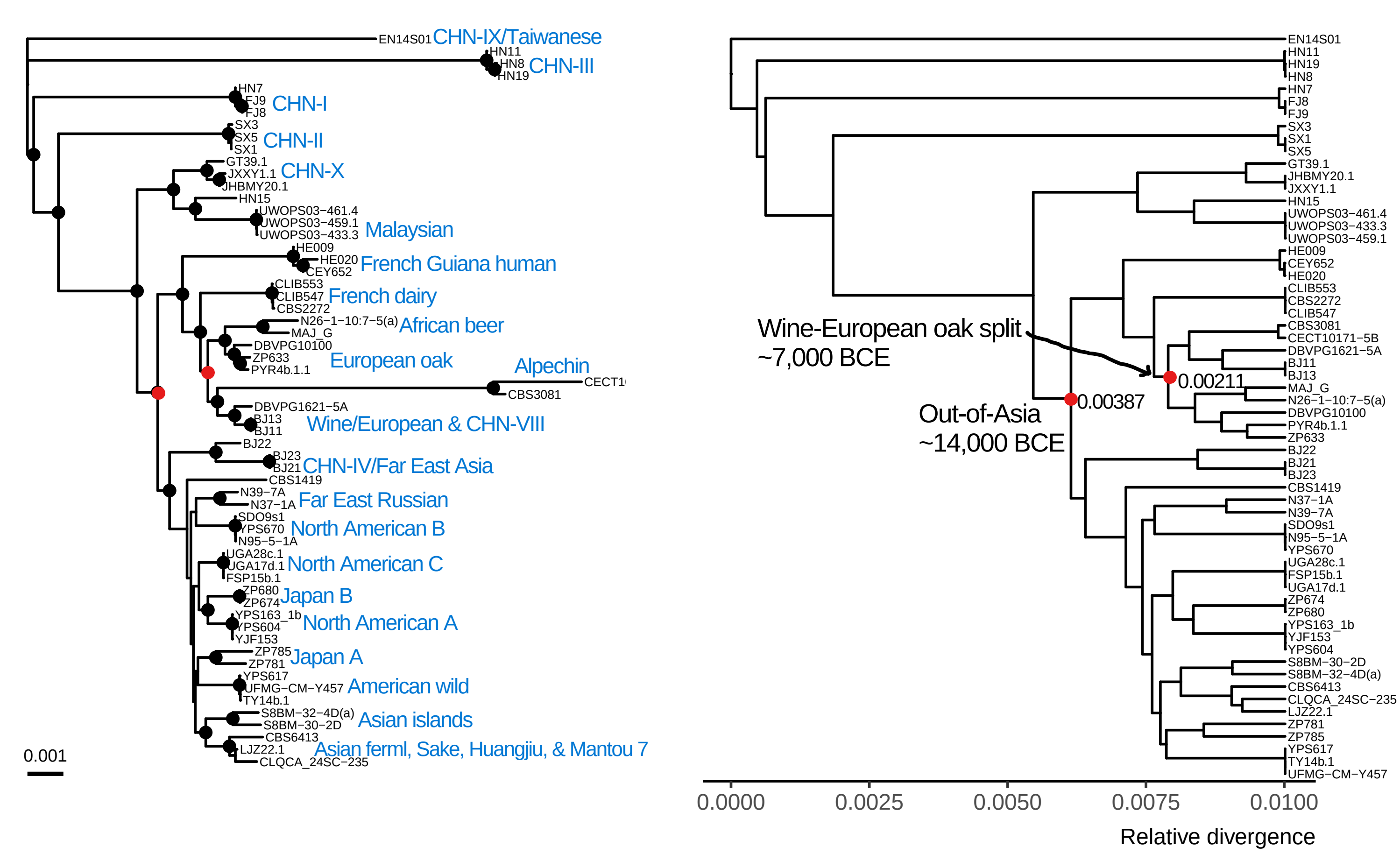

Chromosome 5

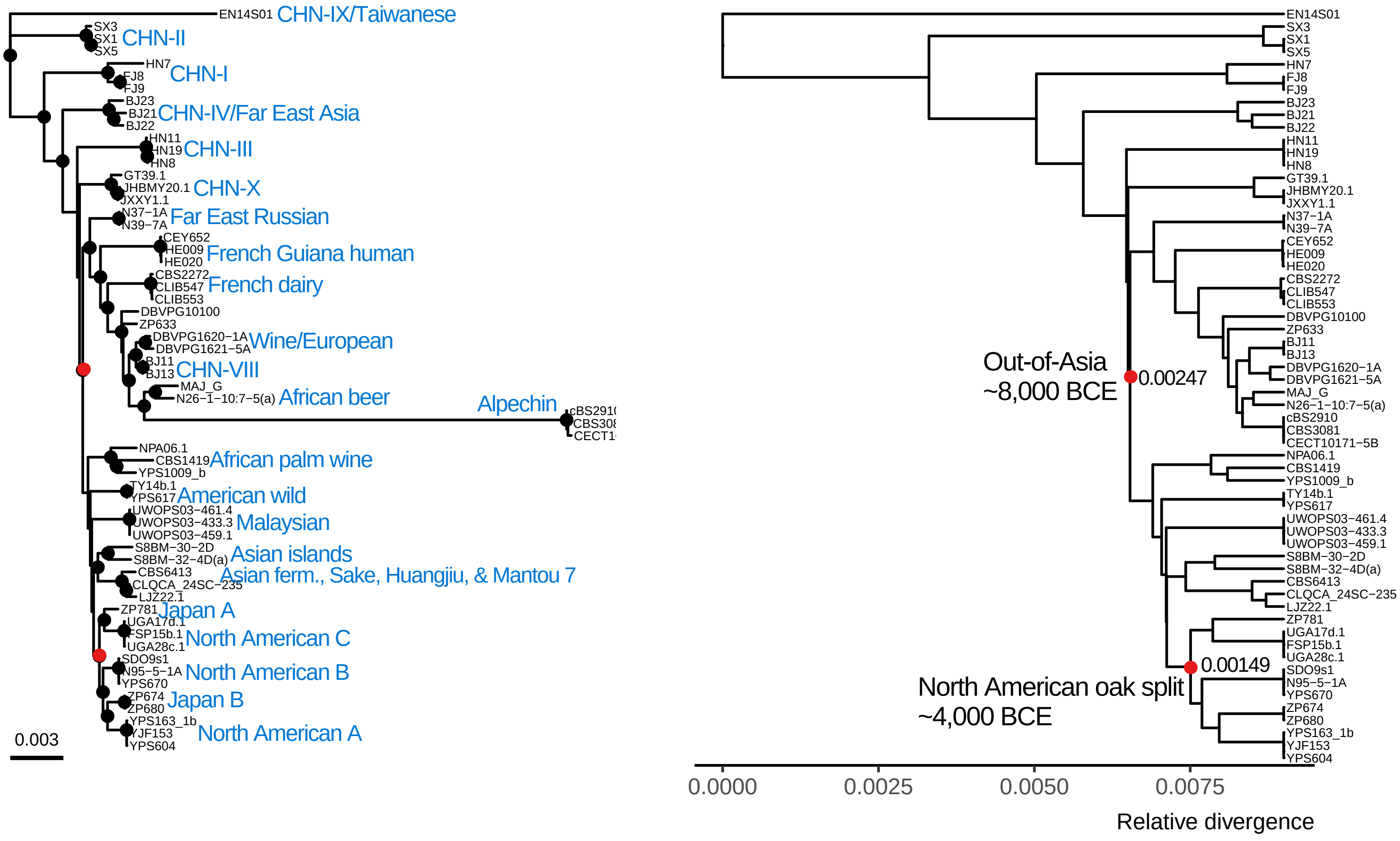


Chromosome 6

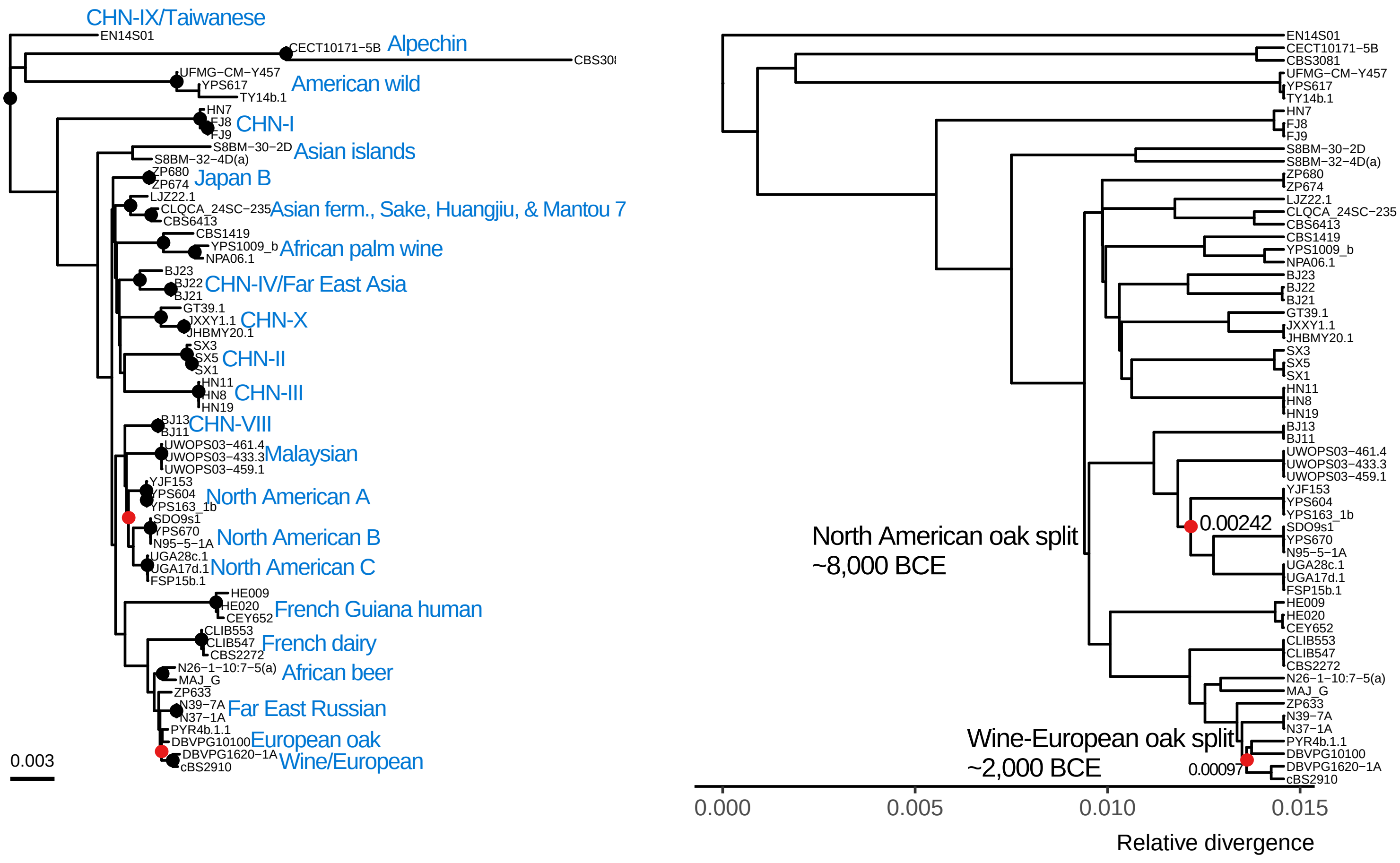

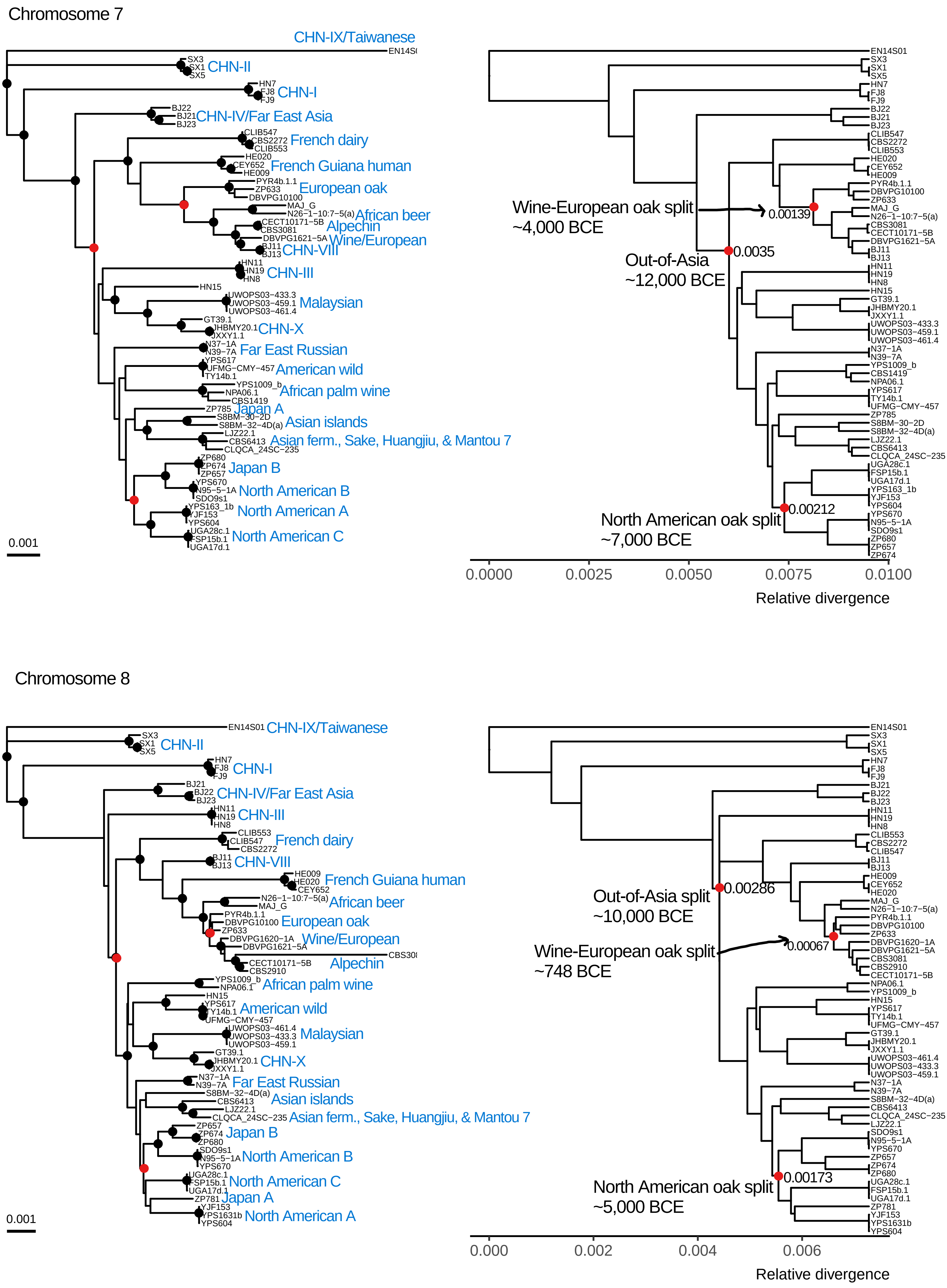

Chromosome 7
CHN-IX/Taiwanese
CHN-II
CHN-I
CHN-IV/Far East Asia
French dairy
French Guiana human
European oak
African beer
Alpechin
Wine/European
CHN-VIII
CHN-III
Malaysian
CHN-X
Far East Russian
American wild
African palm wine
Japan A
Asian islands
Asian ferm., Sake, Huangjiu, & Mantou 7
Japan B
North American B
North American A
North American C
0.001
Wine-European oak split
~4,000 BCE
0.00139
Out-of-Asia
~12,000 BCE
0.0035
North American oak split
~7,000 BCE
0.00212
0.0000
0.0025
0.0050
0.0075
0.0100
Relative divergence
Chromosome 8
CHN-IX/Taiwanese
CHN-II
CHN-I
CHN-IV/Far East Asia
CHN-III
French dairy
CHN-VIII
French Guiana human
African beer
European oak
Wine/European
Alpechin
African palm wine
American wild
Malaysian
CHN-X
Far East Russian
Asian islands
Asian ferm., Sake, Huangjiu, & Mantou 7
Japan B
North American B
North American C
Japan A
North American A
0.001
Out-of-Asia split
~10,000 BCE
0.00286
Wine-European oak split
~748 BCE
0.00067
North American oak split
~5,000 BCE
0.00173
0.000
0.002
0.004
0.006
Relative divergence

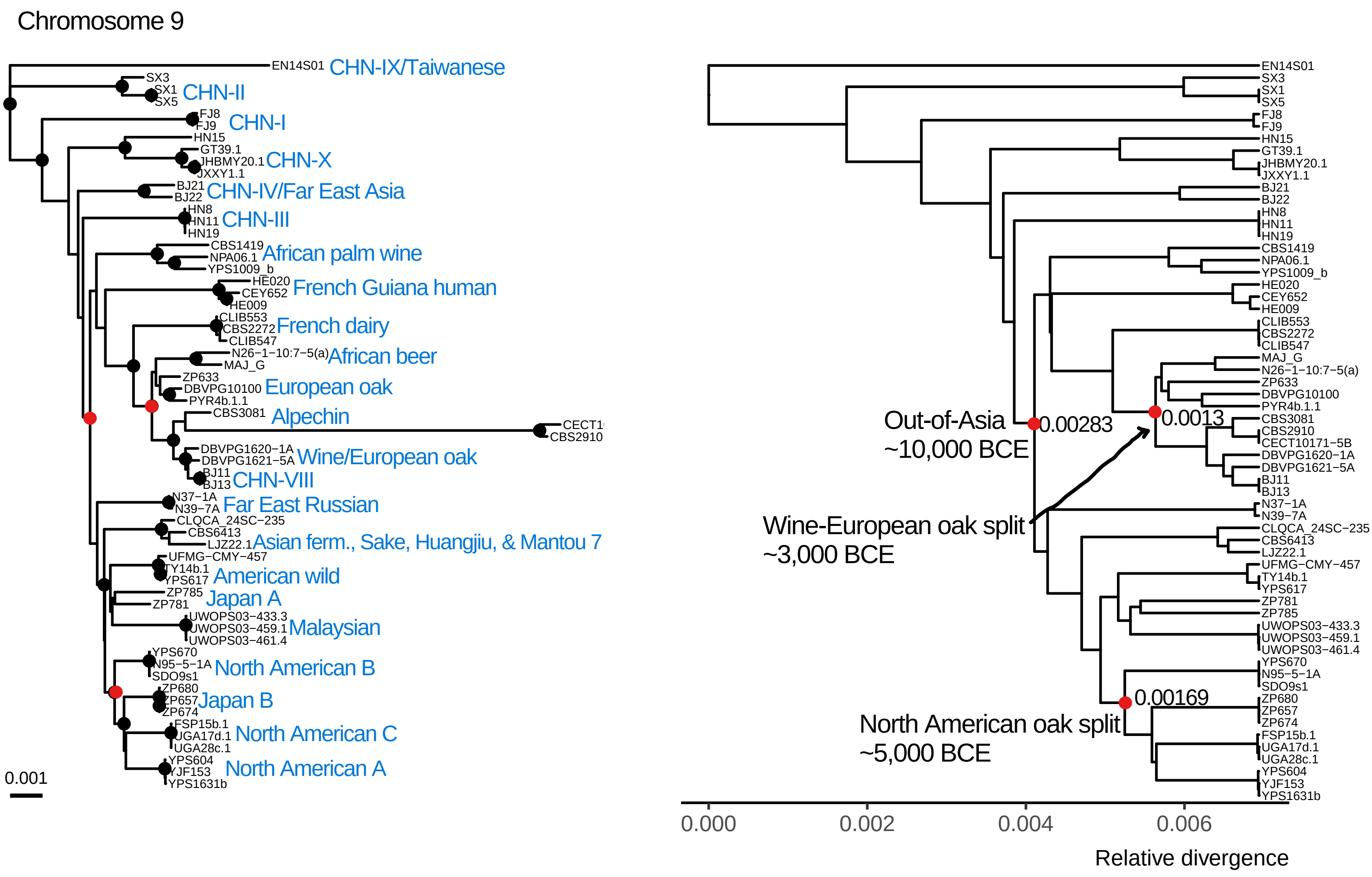

Chromosome 9
CHN-IX/Taiwanese
CHN-II
CHN-I
CHN-X
CHN-IV/Far East Asia
CHN-III
African palm wine
French Guiana human
French dairy
African beer
European oak
Alpechin
Wine/European oak
CHN-VIII
Far East Russian
Asian ferm., Sake, Huangjiu, & Mantou 7
American wild
Japan A
Malaysian
North American B
Japan B
North American C
North American A
0.001
Out-of-Asia
~10,000 BCE
0.00283
0.0013
Wine-European oak split
~3,000 BCE
0.00169
North American oak split
~5,000 BCE
0.000
0.002
0.004
0.006
Relative divergence


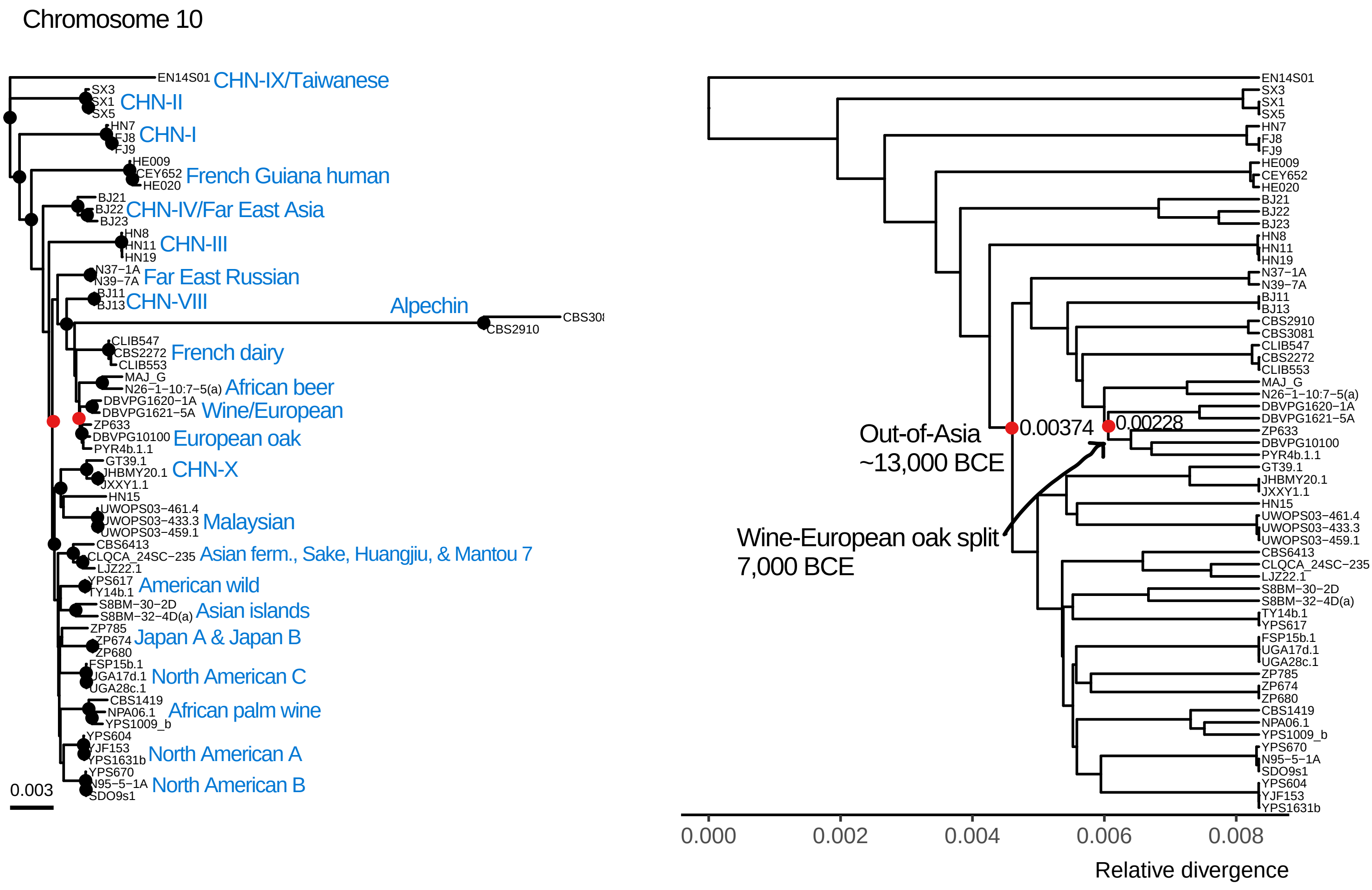

Chromosome 10
CHN-IX/Taiwanese
CHN-II
CHN-I
French Guiana human
CHN-IV/Far East Asia
CHN-III
Far East Russian
CHN-VIII
Alpechin
French dairy
African beer
Wine/European
European oak
CHN-X
Malaysian
Asian ferm., Sake, Huangjiu, & Mantou 7
American wild
Asian islands
Japan A & Japan B
North American C
African palm wine
North American A
North American B
0.003
Out-of-Asia
~13,000 BCE
0.00374
0.00228
Wine-European oak split
7,000 BCE
0.000
0.002
0.004
0.006
0.008
Relative divergence

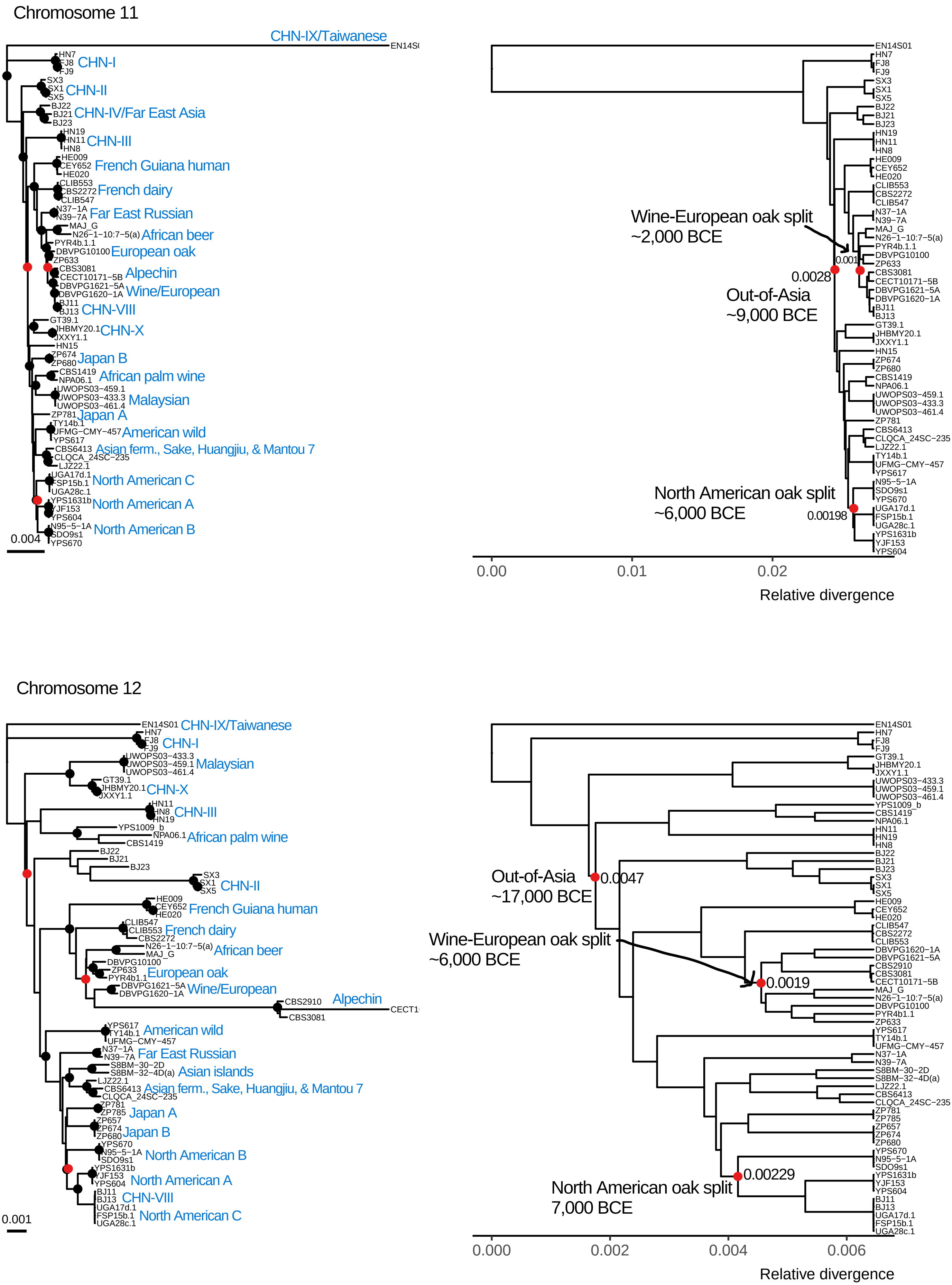
Chromosome 11
CHN-IX/Taiwanese
CHN-I
CHN-II
CHN-IV/Far East Asia
CHN-III
French Guiana human
French dairy
Far East Russian
African beer
European oak
Alpechin
Wine/European
CHN-VIII
CHN-X
Japan B
African palm wine
Malaysian
Japan A
American wild
Asian ferm., Sake, Huangjiu, & Mantou 7
North American C
North American A
North American B
0.004
Wine-European oak split
~2,000 BCE
0.001
0.0028
Out-of-Asia
~9,000 BCE
North American oak split
~6,000 BCE
0.00198
0.00
0.01
0.02
Relative divergence
Chromosome 12
CHN-IX/Taiwanese
CHN-I
Malaysian
CHN-X
CHN-III
African palm wine
CHN-II
French Guiana human
French dairy
African beer
European oak
Wine/European
Alpechin
American wild
Far East Russian
Asian islands
Asian ferm., Sake, Huangjiu, & Mantou 7
Japan A
Japan B
North American B
North American A
CHN-VIII
North American C
0.001
Out-of-Asia
~17,000 BCE
0.0047
Wine-European oak split
~6,000 BCE
0.0019
North American oak split
7,000 BCE
0.00229
0.000
0.002
0.004
0.006
Relative divergence

Chromosome 13

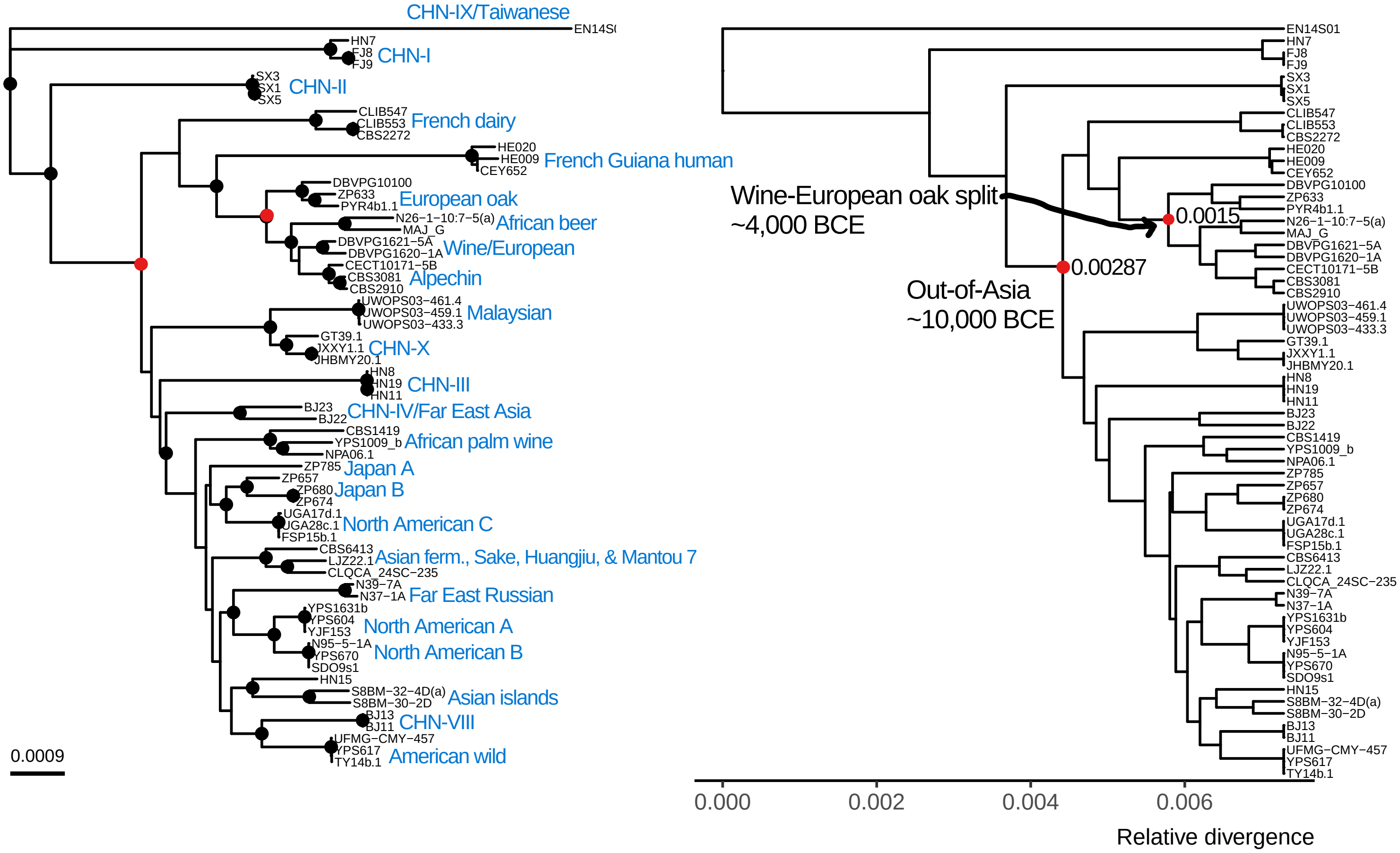


Chromosome 14

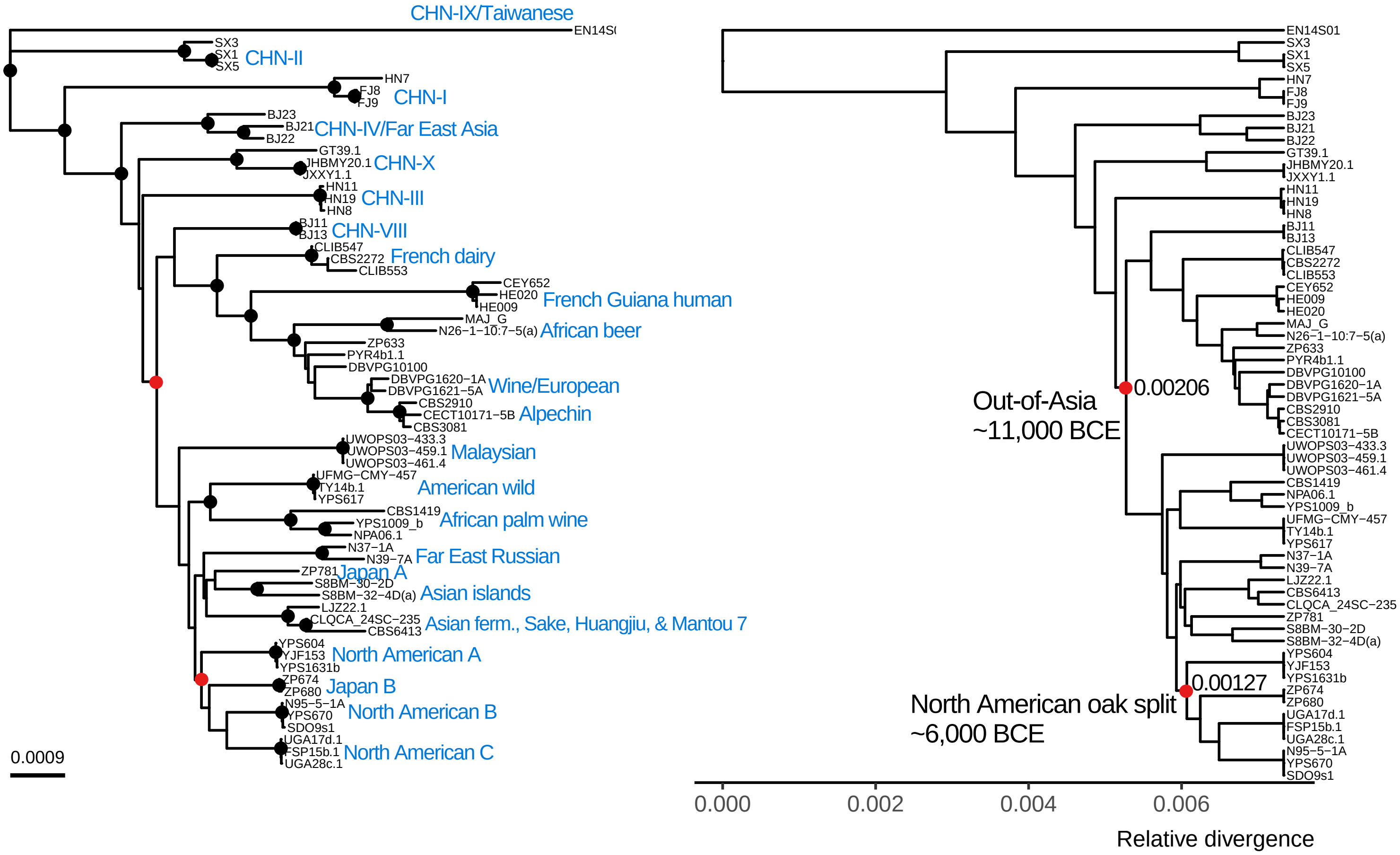

Chromosome 15

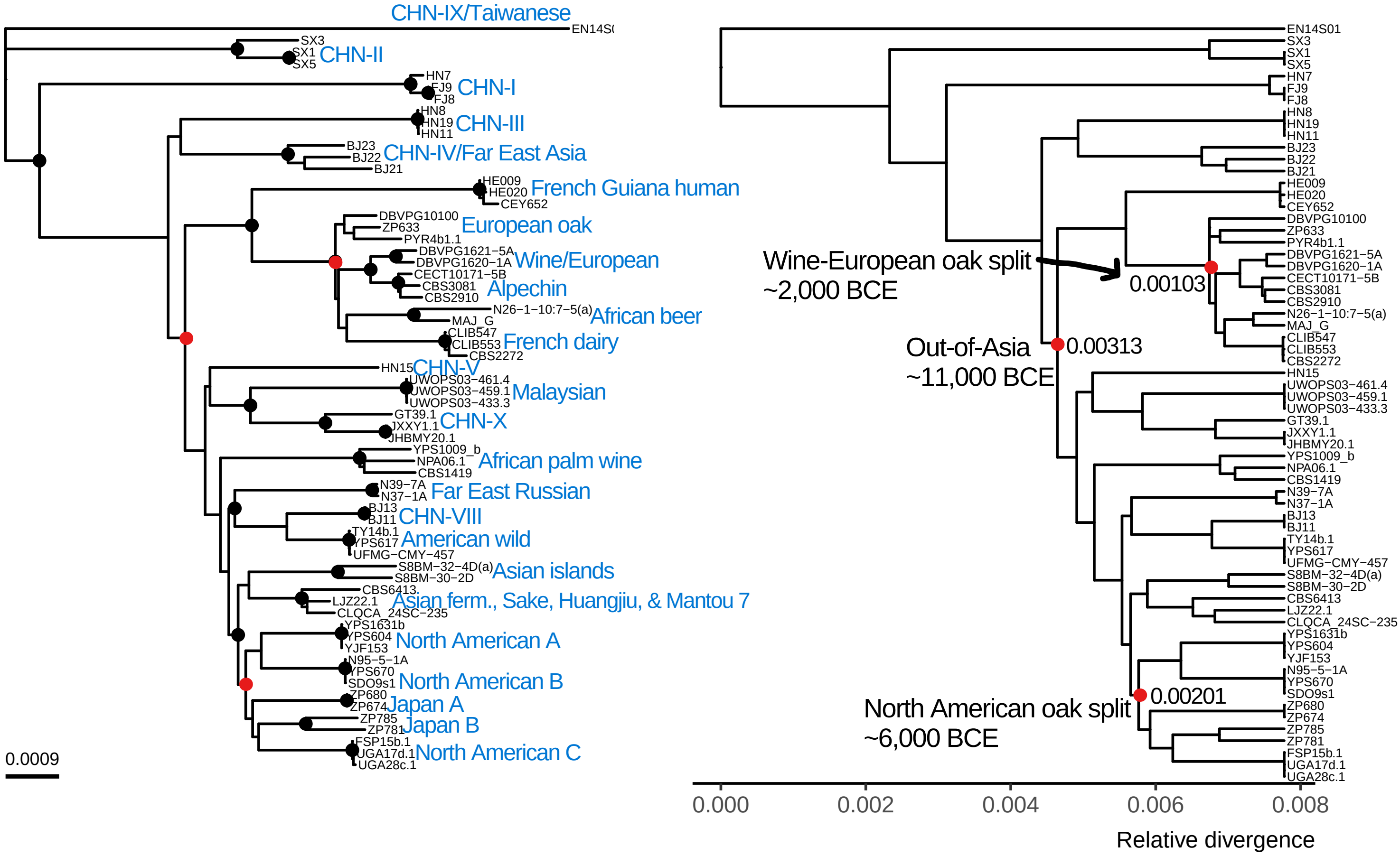


Chromosome 16

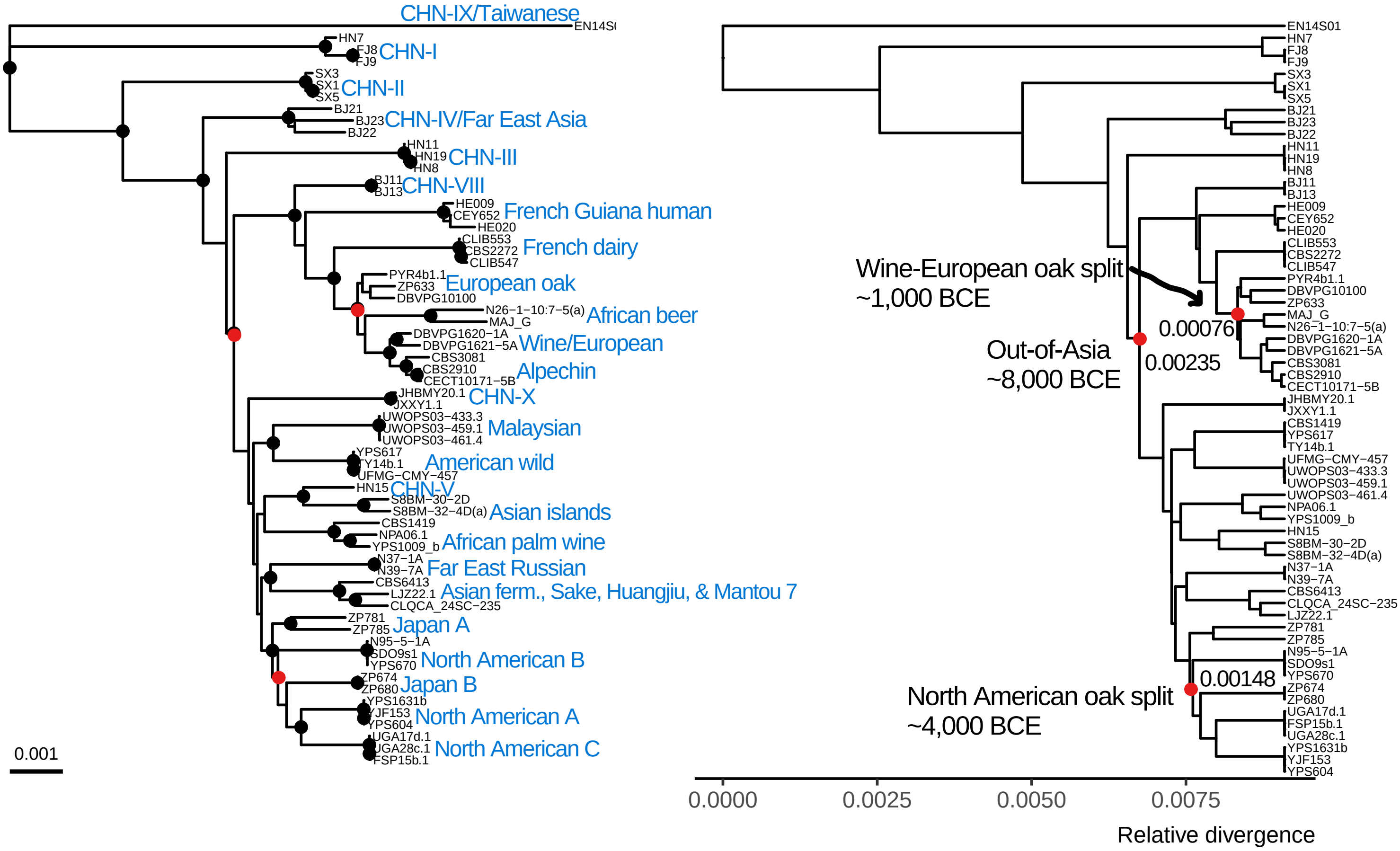
